# Specific Codons Control Cellular Resources and Fitness

**DOI:** 10.1101/2022.09.21.508913

**Authors:** Aaron M. Love, Nikhil U. Nair

## Abstract

As cellular engineering progresses from simply overexpressing proteins to imparting complex metabolic and regulatory phenotypes through multigene expression, judicious appropriation of cellular resources is essential. Since there is degeneracy in codons and their use is biased, codons may control cellular resources at a translational level. We investigate how partitioning tRNA resources by incorporating dissimilar codon usage can drastically alter interdependence of expression level and burden on the host. By isolating the effect of individual codons’ use during elongation, while eliminating confounding factors like mRNA structure, GC content, transcript level, and translation initiation rates, we show that codon choice can *trans*-regulate fitness of the host and expression of other heterologous genes. We correlate specific codon usage patterns with host fitness, and derive a coding scheme for multi-gene expression called the Codon Health Index (CHI, χ). This empirically derived coding scheme (χ) enables the design of multi-gene expression systems that avoid catastrophic cellular burden and is robust across multiple growth conditions.

## Introduction

The genetic code is degenerate with 61 codons and only 20 amino acids, creating an astronomically high level of mRNA sequence space for most protein coding genes. However, it is well accepted that synonymous codons are not equivalent (*1*, *2*), as numerous reports of *cis* and *trans* effects have been documented (*3–11*) – from mRNA structure and co-translational protein folding (*12–15*) to tRNA and ribosome competition (*16–18*). Re-coding genes typically proceeds through use of a codon adaptation index (CAI), which facilitates re-coding towards the codon usage bias (CUB) of a reference, often a set of highly expressed genes (*19*). This strategy may generally correlate CUB with protein expression, but it ignores the role CUB can play in partitioning translational resources such as tRNA and ribosomes. Several recent studies have demonstrated the ability of heterologous genetic CUB to *trans-*regulate host gene expression through translational resource completion (*20*, *21*). There are also established theoretical models (*22*, *23*) and experimental studies to support specific codon use (*24*) and tRNA availability (*25*) as the primary determinant of elongation rate and fidelity. Unfortunately, there are few accounts describing how specific CUB alters host fitness given that cellular resources are invariably limited. Re-coding strategies such as the tRNA adaptation index (tAI) (*7*, *26*) and normalized translational efficiency (nTE) (*6*) are attempts to address tRNA related translational supply-demand constraints, but they are limited by how predictive natural CUB and/or tRNA levels are for recombinant protein expression.

It is particularly important to consider translational resource competition in the context of multi-gene expression (e.g., in the case of metabolic engineering and synthetic biology), where the objective is often for overall strain performance rather than high expression level alone. When optimizing microbial systems, tradeoffs in expression level can be highly consequential for pathway or genetic circuit function and robustness (*27*). Overexpression of a new heterologous gene will often reduce levels of existing proteins (*28*), creating difficulty in engineering predictable cellular behavior. The contextual and variable performance of biological systems is well documented, as many have published how resource competition limits applied research in synthetic biology (*29–31*). Despite a growing body of work, this area remains underexplored.

Many studies to date focus on feedback control mechanisms (*28*, *32*) and resource partitioning (*33*, *34*) between host and engineered components to improve the function of synthetic constructs. Approaches focused on improving gene design have generated a lot of data, but often attempt to draw inferences about elongation in larger genes from libraries limited to the 5’ sequence of a reporter (*3*, *35*), and experiments that do not isolate translation elongation from initiation effects (*10*). While a role for CUB in the partitioning of cellular resources has been reported (*36*, *37*), identification of specific codons that present excess translational capacity could provide a novel avenue for harnessing underutilized resources that are insofar ignored.

In this study, we systematically isolate the role of codon choices during translational elongation and identify supply-demand constraints imposed on tRNA and ribosomal resources in *Escherichia coli*. We demonstrate that tRNA supply and demand imbalances lead to competition between overexpressed genes as well as with the host’s resource needs and that results cannot be explained by other factors like mRNA structure, GC content, transcript levels, RNA toxicity, or translation initiation rates. We find that select codons that are over-represented in native, highly expressed genes, are found to cause severe fitness costs when present in overexpressed coding sequences.

While the traditional method of codon-optimization through maximizing CAI may promote use of these codons, our data reveal their demand and supply are delicately balanced. We define a new metric called “Codon Health Index” (CHI, χ) that quantitatively ranks codons by their capacity to remain orthogonal to host demands. We also posit using this metric as a new codon optimization scheme to mitigate competition with host demands and avoid growth defects. Genes characterized by high CHI scores demonstrate relatively high expression and minimal burden on the host cells across multiple growth conditions, allowing effective multigene expression and cellular growth.

## Results

### Fitness costs are incurred due to translation elongation limitation

Genetic burden is frequently observed in microbial systems as a growth defect upon the overexpression of recombinant proteins (*32*). While the cause of this effect varies, it is often attributed to resource competition at the level of translation (*38*), not transcription. In a fast-growing culture of *E. coli*, the availability of free ribosomes can limit mRNA translation, especially in a system with overexpressed protein (*39*) (**Fig. 1A**). Elongation speed determines the rate at which free ribosomes are made available, hence sub-optimally coded mRNA that are poorly translated have higher ribosome occupancy, which is something that re-coding often attempts to resolve. Such elongation limited mRNA sequences will sequester more ribosomes and return them to the free pool at a slower rate, thus reducing ribosome availability. Translational resource competition has been modeled in several ways (*40*), including the ribosome flow model (RFM) (*41*), which can be useful in examining translation rate as a function of elongation time that varies depending on the supply and demand of tRNAs in the cell. Applying a previously published RFM (*42*) to cyan and yellow fluorescent proteins (CFP and YFP, respectively) used in this study with high or low CAI values (where CAI is in reference to highly expressed *E. coli* genes) illustrates the increase in mRNA ribosome occupancy that occurs when codons with longer elongation times (*43*) are used. The model further indicates that elongation-limited sequences are less sensitive to changes in the rate of translation initiation (**fig. S1, Data S1**). Elongation limited sequences are therefore predicted to vary less in expression as a function of free ribosome availability, while also creating higher genetic burden by sequestering ribosomes across the mRNA.

**Fig. 1.**
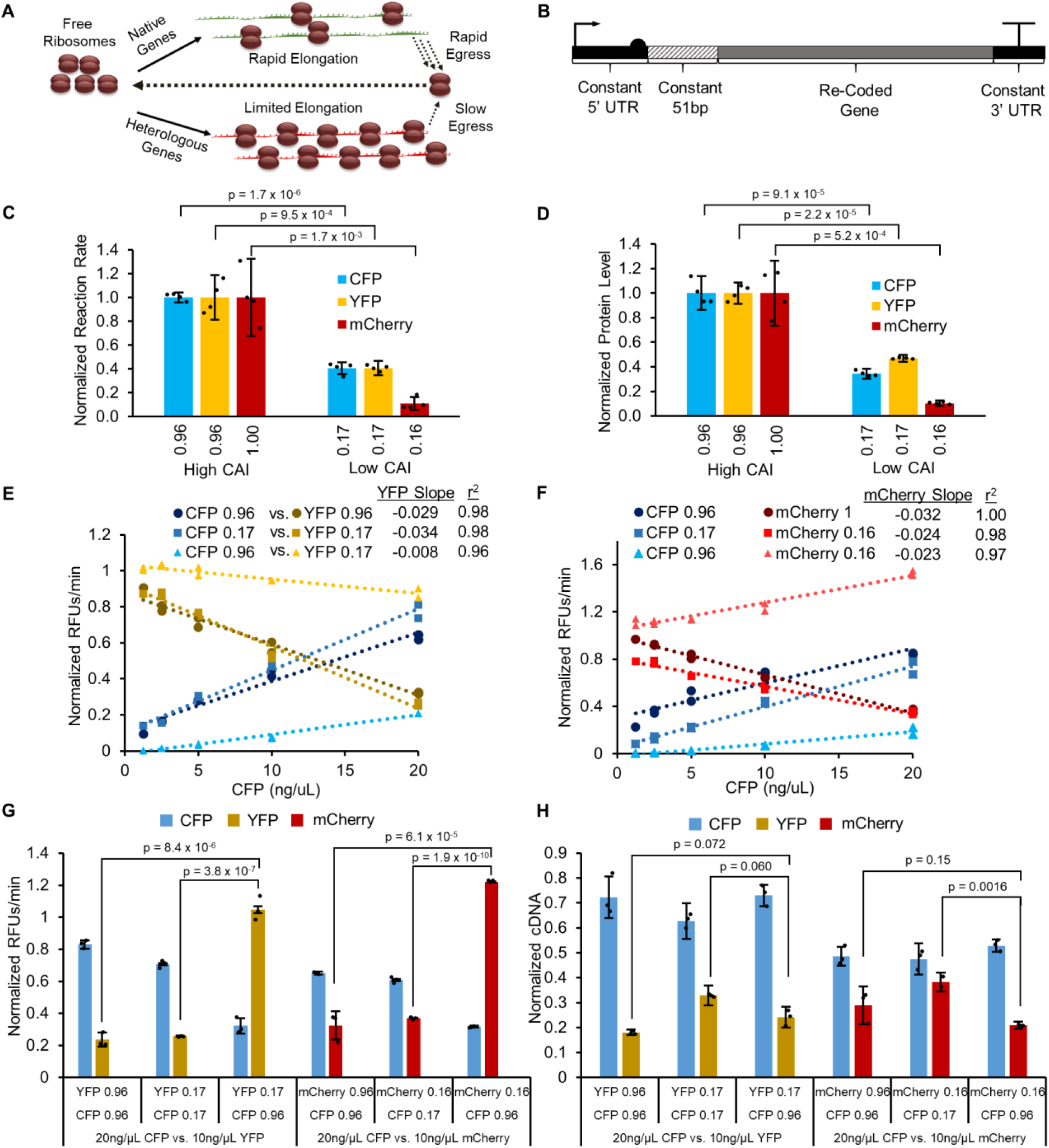
Elongation-limited TxTL system exhibits competition for translational resources. (**A**) Schematic of two mRNAs competing for ribosomes. Genes with low adaptation to tRNA availability exhibit higher ribosomal occupancy and sequester more translational resources than those with optimized codon bias. (**B**) Gene design for expression in TxTL assay. A T7 promoter with a strong RBS (BCD7) drives expression of re-coded genes, where the first 51 bases are constant, as well as the 5’ and 3’ UTR. (**C**). TxTL expression rates for individual genes (*n* = 4 replicates of a single re-code) normalized to respective high CAI variants. CAI calculations reference highly expressed *E. coli* genes. (**D**) Endpoint protein level from the same reactions shown in panel C, quantified using LC/MS targeted proteomics. (**E and F**). Competitive in vitro TxTL results between pairs of CFP and YFP (**E**) or CFP and mCherry (**F**). Protein synthesis rates are normalized to each gene in isolation. YFP and mCherry cassettes (DNA) were each added to a constant 10 ng/µL whereas the CFP cassette was titrated from 1.25 – 20 ng/µL (*n* = 2 replicates of single re-codes for each concentration level, linear regression slope and r^2^ values are calculated from a total of 10 data points for each series). (**G**) Competition data for sequence pairs in panels E and F repeated with 20ng/µL CFP and 10ng/µL YFP or mCherry (*n* = 3). Protein synthesis rates are again normalized to individual gene expression in isolation. (**H**) qRT-PCR results for sequence pairs shown in G. Absolute cDNA levels were measured and normalized to cDNA from individual genes expressed in isolation. Yellow, blue, or red lines and bars indicate the production of YFP, CFP, or mCherry proteins respectively. All p values represent two tailed t tests. All bars represent means ± SD.

With the ribosome flow model as a guide, we first sought to investigate the impact of resource competition during translation elongation using a well-controlled in vitro transcription-translation (TxTL) model. A significant challenge to investigating translational resource competition is the difficulty in isolating any single sequence parameter experimentally, as any synonymous mutation can have a multitude of effects on initiation, elongation, and mRNA structure (*2*). A TxTL system allows for better physical control over the genetic expression environment by holding available resources (i.e., ribosomes, tRNAs, aminoacyl tRNA synthetases, RNA polymerase etc.) constant, and allowing precise titration of genes of interest in the reaction. We developed a resource competition assay for elongation limitation by leveraging the unique amino acid sequence similarity between CFP and YFP derived from a super-folder green fluorescent protein (*44*), which only differ by 2 amino acids (*45*). This virtually eliminates variability in protein structure and amino acid demand, thus enabling the interrogation of sequence designs using effectively identical proteins. These proteins should also be minimally susceptible to variation in co-translational protein folding due to their rapid folding and high stability. We also include mCherry in the study, which is <30% identical to CFP-YFP and serves as a comparison to find trends independent of amino acid sequence (see supplementary data for sequences). The implemented TxTL assay is based off the *E. coli* MRE600 strain, which has a nearly identical CUB as the closely related *E. coli* K12 MG1655 strain used later in our work, and is therefore assumed represent the expected tRNA profile in vivo (**fig. S2**). Reactions were driven by a T7 promoter using a bicistronic domain (BCD) in place of a traditional ribosome binding site to minimize interactions between the 5’ untranslated region and gene of interest that could lead to differential expression (*46*). To further isolate translation elongation as the primary variable in sequence design, we chose to keep the 5’ and 3’ untranslated regions (UTRs) as well as the first 51 base pairs (17 codons) constant to mitigate any effect sequence changes may have on translation initiation (**Fig. 1B**).

Utilizing our optimized TxTL competition assay, we evaluated baseline expression rates from CFP, YFP, and mCherry re-codes with extreme CAI values (0.96 or 0.16) (**Fig. 1C, Data S2**). The CFP and YFP CAI value of 0.96 (as opposed to 1.0) results from starting with an existing sequence lower than CAI = 1 and holding the first 51 bases constant. Individual expression cassettes (DNA) were each added to 10 ng/µL, and data are normalized for each fluorescent reporter to the high CAI variant expressed in isolation. We find that protein expression rates for CFP, YFP, and mCherry correspond well with CAI value. Low CAI values (rare codon use) reduce protein synthesis rate across all tested genes. This significant correspondence supports our assumption that the relative tRNA concentrations in the TxTL assay are similar to previously reported tRNA levels (*18*). We further validated that the observed differences in fluorescent measurements correspond to changes in protein level (as opposed to misfolded protein) using LC-MS/MS (**Fig. 1D**). This supports that TxTL can simulate translation elongation limitation – i.e., genes with low CAI that use low abundance tRNAs show low protein synthesis rates. Next, we examined competition between different pairs of genes. As in the RFM, we expected elongation-limited sequences with lower CAI to disrupt expression of other genes through the sequestration of free ribosomes. We titrated CFP template DNA against constant YFP or mCherry DNA using the same re-codes with either very high or very low CAI (**Fig. 1 E and F, Data S3**). Reactions were run for each pair of sequences tested with five concentration levels of CFP. Competitive reaction rates for each protein were normalized respectively to reaction rates of proteins in isolation. Linear regression was performed to calculate slope values using 10 replicates per series. More negative slopes for normalized YFP and mCherry expression rates indicate higher competition for resources. For instances of two identically re-coded sequences with any CAI tested, YFP and mCherry synthesis rates are inversely correlated with CFP DNA concentration, irrespective of their baseline expression, indicating strong competition for limiting resources (e.g., tRNA). This indicates that while an apparent excess protein synthesis capacity exists in the TxTL system for low CAI sequences, they are clearly resource-limited, likely due to lower availability of tRNA.

More interesting observations are seen when dissimilar CAI re-codes are under competition upon co-expression (CAI 0.96 vs. CAI 0.16 or 0.17). In these cases, low CAI YFP and mCherry synthesis rates are not very sensitive to increasing resource consumption by high CAI CFP synthesis. Conversely, the relative CFP expression is much lower than we observed either in isolation or when competing with a high CAI sequence. The observed results are consistent for both sequence pairs, indicating that this phenomenon is independent of protein sequence. CFP with low CAI was also titrated against YFP or mCherry with high CAI and was found to monopolize resources at low DNA concentrations, and caused non-linear reduction in YFP or mCherry expression (**fig. S3**). We further confirmed that resource competition between sequence pairs occurs at the level of translation and not transcription with quantitative reverse transcription PCR (qRT-PCR) by repeating the experiment under the highest level of competition (20 ng/µL CFP vs. and 10 ng/µL YFP or mCherry) (**Fig. 1G and H, Data S4**). Both YFP and mCherry are again negatively affected by CFP with identical codon use, but the rare codon rich sequences are significantly less affected under competition with high CAI CFP (**Fig. 1G**). It is unclear why expression for low CAI sequences under competition exceeds the expression rate in isolation (value >1), but this could be due to the presence of additional total DNA in the reaction having a stabilizing effect. We performed qRT-PCR on endpoint samples (**Fig. 1H, fig. S4**) to determine differences in mRNA level and quantified absolute cDNA levels with standard curves to enable analysis across different gene amplicons and found that mRNA level does not explain the differences we see in protein expression rate between pairs of re-coded sequences. For both YFP and mCherry, low CAI sequences competing against high CAI CFP have either lower or similar mRNA levels to the other conditions, despite exhibiting higher relative translation rates.

When examined in the context of an RFM, the model suggests that rare codon enriched YFP and mCherry sequences sequester ribosomes to such a degree that even excess CFP template DNA does not yield high protein synthesis rates. In this scenario, YFP and mCherry protein synthesis rates are expected to be unaffected by changes in free ribosome concentration, and translation initiation rates would not be affected due to severe elongation limitation (**fig. S1**). While we do not measure ribosome occupancy directly, this model is supported by our experimental design and several observations. Our results are not explained by differences in transcription (**Fig. 1H**), and we have designed our expression cassettes to minimize any influence the different gene coding has on translation initiation (**Fig. 1B**). This model is further supported by our observation that both YFP and mCherry rates are reduced due to *trans* effects when competing between two low CAI CFP sequences, which is a likely consequence of competition for scarce tRNAs based on reported tRNA levels in *E. coli* (*18*). Overall, our data indicate that proteins coded with similar CAI (high or low) are strongly competitive due to demand for translational resources. Conversely, genes coded under low CAI regimes are constrained by the availability of tRNA and can dominate the availability of protein translation resources. Our TxTL data strongly support the argument that translation elongation limitation could play an important role in cellular resource competition and highlights the impact to global translational resources (e.g., free ribosomes, tRNA) in multigene expression environments.

We next set out to optimize an in vivo assay for *E. coli* protein expression to efficiently interrogate the effect that mRNA coding has on gene expression and host fitness. Our system generally consists of a strong constitutively expressed YFP reporter gene (CAI = 0.96 unless otherwise noted) integrated into the *E. coli* chromosome paired with an inducible CFP or mCherry gene on a plasmid driven by the inducible promoter P_trc_ with a strong RBS (**Fig. 2A**). As before, we held the first 51 bases and the 5’ and 3’ UTRs constant for all re-codes to prevent unwanted effects on translation initiation. Cells grown in rich medium with a common pre-culture were passaged under inducing or non-inducing conditions. The area under the curve (AUC) is used to measure each of the signals (growth and fluorescence), which captures the aggregate effects of lag phase, expression and growth rate, and carrying capacity into a single value (**fig. S5**). We define fitness as the ratio of AUC induced vs. uninduced, which ranges from 0 to 1 for low and high fitness, respectively (or conversely, high and low burden). Fitness can be in terms of Growth Fitness based on OD_600_, or Co-expression Fitness based on YFP fluorescence (chromosomal reporter), while Expression Level is based on CFP or mCherry fluorescence (i.e., the overexpressed protein) (**Fig. 2B–D**). Throughout our work we focus mainly on Co-Expression Fitness, since the single term captures differences in growth and cell specific protein production, and best enables selection for designs that enable improved multi-gene expression with reduced resource competition. We also validated that protein level correlates very well with our fluorescence measurements using LC/MS based proteomics (**fig. S6**).

**Fig. 2.**
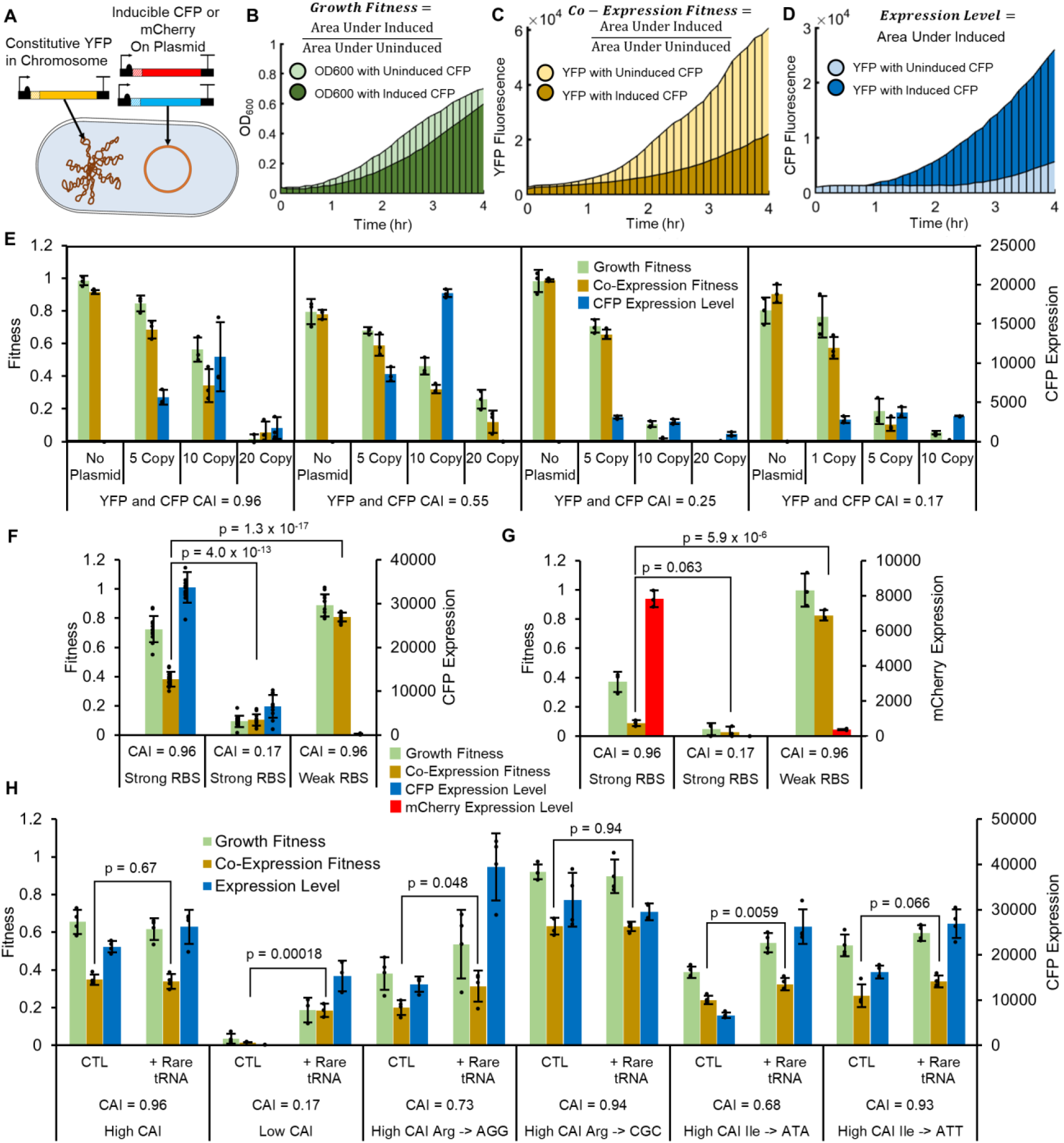
Optimization of *in vivo* fitness assay. (**A**) Schematic of the in vivo system, with constitutive YFP integrated into the chromosome (CAI = 0.96) of *E. coli*, and inducible CFP or mCherry (varying CAI) expressed from plasmid. (**B**) Example OD600 data with a Growth Fitness cost due to increased CFP expression. (**C**) Example YFP fluorescence data with a Co-expression Fitness cost due to induced CFP. (**D**) Example fluorescence data used to determine Expression Level of CFP. (**E**) Fitness and expression data for re-coded CFP-YFP pairs with co-varying CAI, where CFP is expressed from different copy number plasmids (*n* = 3). Matching CFP and YFP CAI values are indicated in the figure. 1 copy = f1 origin, 5 copy = pSC101 origin, 10 copy = p15A origin, 20 copy = pBR322 origin. (**F** and **G**) Analysis of fitness and Expression Level for control CFP (*n* = 12) and mCherry (*n* = 3) sequences with strong and weak RBS supports that burden created by gene expression is dependent on translation. CFP or mCherry are expressed from a 10-copy vector with indicated CAI, whereas YFP (CAI = 0.96) is constitutive and chromosomally integrated. (**H**) Analysis of fitness and Expression Level for CFP sequences re-coded with rare codons and complemented with corresponding rare tRNA on a single copy vector pBAC-RARE2 (*n* = 3). The presence of multiple rare codons for one or several amino acids leads to translational resource limitation and Growth and Co-Expression Fitness costs (along with decreased Expression Level). Fitness and Expression Level can be recovered by supplementing rare tRNAs in *trans*. All points are biological replicates of individual re-codes. All p values represent two tailed t tests, and all bars represent means ± SD.

Upon varying plasmid copy number with several pairs of CFP and YFP with different matching CAI levels, we found that fitness costs (Co-expression and Growth) were strongly dependent on copy number and are further exacerbated by low CAI (**Fig. 2E, Data S5**). High CAI CFP generally expresses well but elicits a significant fitness cost in terms of Growth Fitness and Co-expression Fitness. For a CFP re-coded with rare codons, the result is catastrophic, and cultures are unable to grow at all with high-copy vectors, which has also been reported elsewhere (*20*). Based on these results, we picked the 10-copy vector (p15A origin) with the YFP CAI = 0.96 reporter as the platform for further studies to investigate re-coding schemes that may reduce fitness costs. Using a multi-copy vector is expected to simulate the fitness costs from multiple genes expressed at lower levels on multiple vectors or from the chromosome, while simplifying the experimental design. Evaluating controls in this system for CFP (**Fig. 2F, Data S6**) and mCherry (**Fig. 2G, Data S7**), we observe significant reduction in Co-Expression fitness with a high CAI coding for both genes. While RNA toxicity has been reported elsewhere (*47*), the effect appears mediated by translation (not transcription) since using the same promoter with a very weak RBS for CFP and mCherry results in low protein but high fitness.

We characterized tRNA limitation in our experimental system by changing arginine and isoleucine codons in the high CAI CFP sequence to rare codons. Holding the first 51 bases constant, we re-coded all 8 arginines from CGT to AGG or 10 isoleucines from ATT to ATA, expecting these very rare codons to produce a significant effect on Growth Fitness and Co-Expression Fitness. Each sequence was tested with or without complementation with pBAC-RARE2, which overexpresses 12 rare tRNAs, including those in the CFP variants (**fig. S7**). We also included arginine and isoleucine re-codes with frequent codons as controls, expecting a minimal impact on fitness since their corresponding tRNAs are not limiting. We found that incorporation of rare codons compared to frequent codons reduces Expression Level and Co-Expression Fitness (**Fig. 2H**). Upon complementation of pBAC-RARE2, we see that tRNA supplementation significantly improves Co-Expression Fitness, particularly in rare-codon re-coded sequences. This result provides direct evidence that tRNA limitation can reduce co-expression of multiple genes in vivo.

### Systematic analysis of codon use reveals supply and demand constraints in translational resources

Prior to designing re-coded genes that moderate translation elongation resources, we first thoroughly investigated CUB in the *E. coli* transcriptome. CAI calculations are typically based on the natural CUB in highly expressed genes. CUB can be represented as a 64-dimensional space (total number of codons) using RSCU values (observed vs. expected frequency) for each protein coding gene. Initial analysis using hierarchal clustering revealed that groups of genes within the *E coli* transcriptome cluster according to distinct CUB schemes (**fig. S8**). We focused on a consolidated set of this sequence space by analyzing all operons with at least 2 protein coding genes, given that functionally related genes that naturally cluster have similar CUB (**fig. S9**), offering a convenient method to reduce the size of the analysis. The resulting 64 dimensions of codon usage across 773 operons can be represented in 2 dimensions accounting for 41.2% of total variance (**fig. S10**) using principal component analysis (PCA) as shown in **Fig. 3A**. The loading vectors mapped onto the plot represent the 10 codons that contribute most significantly to codon bias across the 773 operons. A similar result was found when the analysis was performed on individual genes instead of operons (**fig. S11**), but we found the operon analysis to be more informative due to lower dimensionality, and the natural clustering of functionally related genes.

**Fig. 3.**
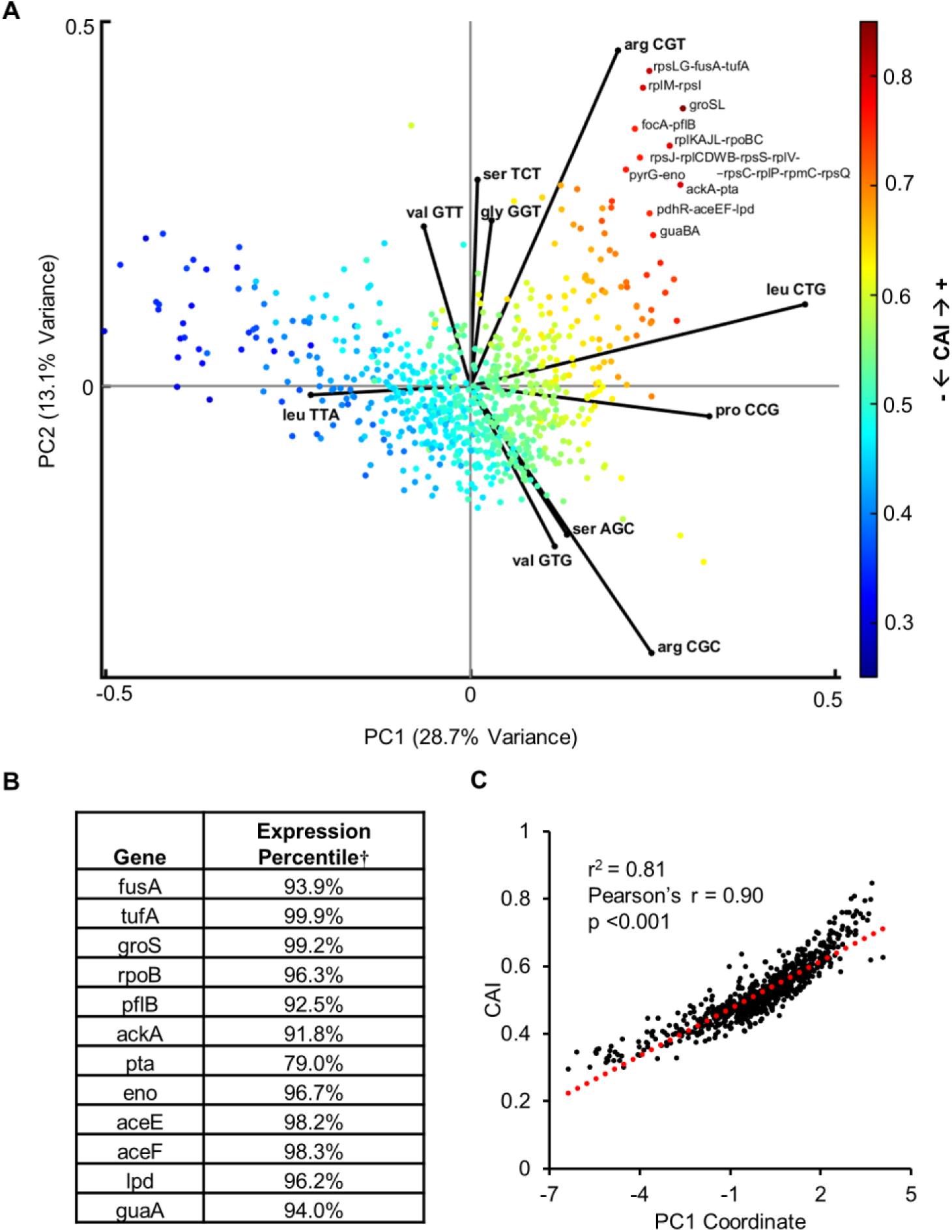
Codon usage bias in highly expressed *E. coli* genes. (**A**) PCA analysis of RSCU in 773 *E. coli* operons with loadings mapped for the 10 codons with the highest contribution to variance. CAI is mapped onto individual operons and indicated in the color bar. (**B**) Select genes from the most extremely biased operons and their expression percentile. ^†^Expression data from Taniguchi et al. (*48*) (**C**) PC1 is largely explained by CAI with a very strong and significant Pearson correlation. Pearson’s r, and linear regression r^2^ values are calculated from 733 data points.

This analysis captures the CUB naturally observed in the *E. coli* transcriptome and highlights a positive correlation between CAI and expression. This is expected because here CAI is calculated by optimizing towards CUB in highly expressed genes (*19*) (see methods) (**fig. S12**). Consistent with previous studies, we corroborate that genes in the most extremely biased CUB space are some of the most highly expressed genes in the *E. coli* proteome that often serve essential functions (**Fig. 3B**). The natural bias leading to the CAI scale is very well explained by PC1 (**Fig. 3C**), suggesting that CAI is the most dominant descriptor of natural codon bias in *E. coli*, however, PC1 only describes 28.7% of total variance in CUB. Despite the apparent correlation between CAI and expression, studies have reported that CAI often does not reliably predict higher gene expression (*10*), which could be due to the 71.3% of natural codon use it fails to model, as well as other sequence features reviewed elsewhere including secondary structure, codon co-occurrence bias, and codon pair bias to name a few (*2*). Additionally, other types of CUB have likely evolved towards objectives other than high expression, such as the pressure to balance cellular resources. Importantly, the CAI paradigm of re-coding proteins to match the CUB of highly expressed genes ignores potential resource competition that can occur at the tRNA level.

For 18 of 20 amino acids, multiple codons exist, and 10 of 18 of those can be coded to use different tRNAs in *E. coli* K12 MG1655 (**fig. S13**). Upon examining the PCA loadings, there are clearly particular codons that are very overrepresented in highly expressed proteins (e.g., arg CGT, leu CTG, and pro CCG), and account for the majority of observed CUB. For such high-demand codons, using alternative codon/tRNA pairs, or even codons that recruit tRNAs with weaker affinity, have the potential to reduce translation elongation-based resource competition between overexpressed proteins and native essential and/or highly expressed genes.

Using our optimized in vivo assay, we sought to experimentally determine the contribution of individual codons to gene Expression Level and Co-Expression Fitness. The synonymous codon sequence space that could be explored in even a small gene such as CFP is experimentally intractable. Holding the first 51 bp constant and co-varying all possible synonymous codons would produce a massive library size of 1.8 × 10^104^. While a more constrained codon library is possible, we chose a focused experimental approach by interrogating individual codon contributions to gene Expression Level and Co-Expression Fitness. Starting with a CFP or mCherry sequence having a high CAI (0.96 − 1.0) and using a single codon for each amino acid where the effective number of codons (ENC) = 20 (for details on ENC, see methods), for each amino acid we re-coded every instance to another synonymous codon, resulting in a total of 41 possible re-coded sequences for each gene (64 possible codons – 20 high CAI codons already in use – 3 stop codons not changed) (**Fig. 4A**). Results were normalized in terms of both Expression Level and Co-expression Fitness (defined in **Fig. 2B**) relative to the high CAI parent control and indicate wide ranging benefits or costs (**Fig. 4B**). In several instances, alternative codons improve the normalized mean Co-Expression Fitness across both mCherry and CFP, with a moderate but significant positive correlation. Arginine, proline, and leucine are emphasized since they drive a significant portion of natural CUB and appear to generally improve Co-expression Fitness when selecting alternative codons to those found in highly expressed genes.

**Fig. 4.**
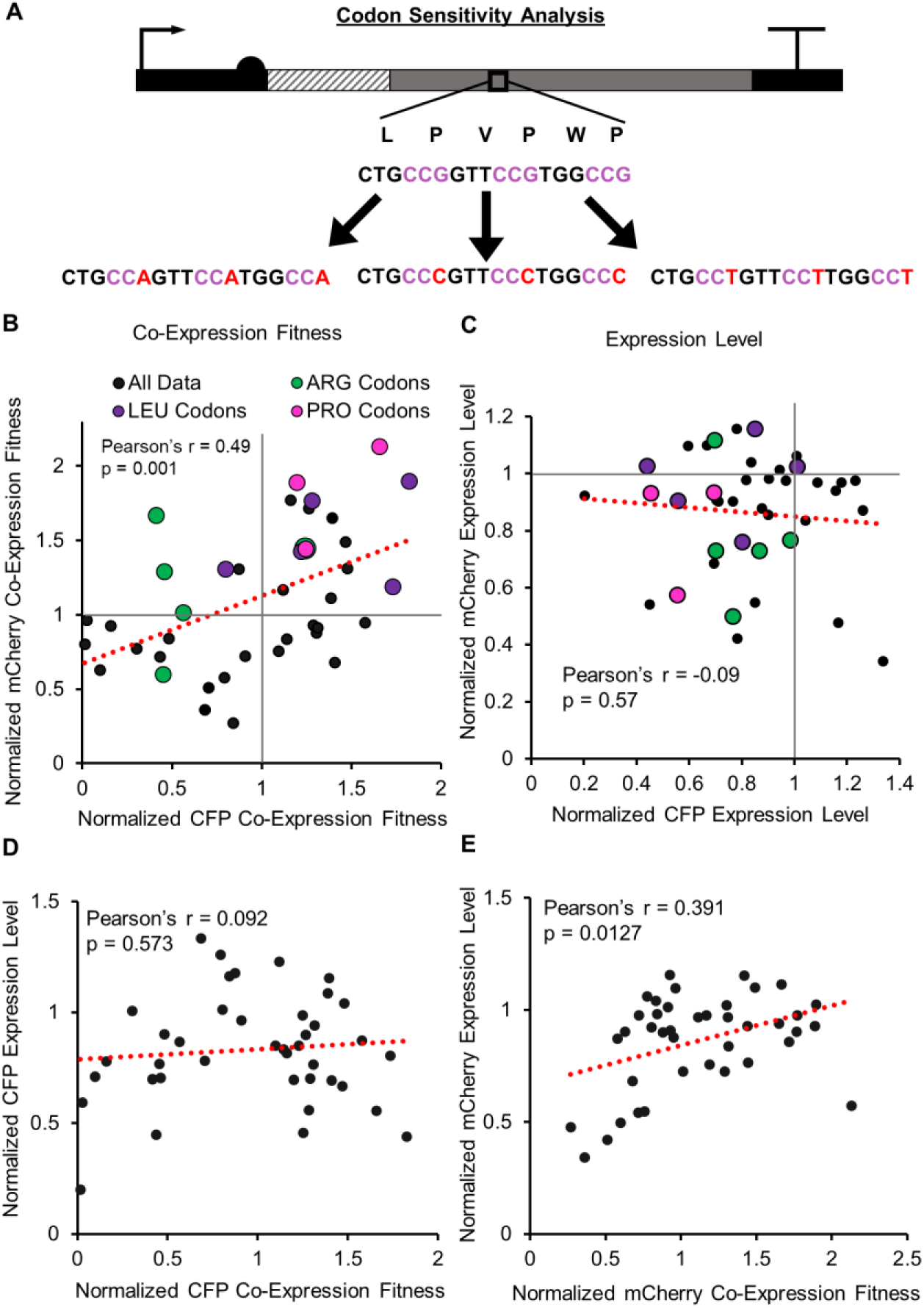
Systematic codon sensitivity analysis. (**A**) Schematic of how genes are re-coded for every amino acid. Starting with the highest CAI weighted codon for every instance of each amino acid, they are re-coded to alternative synonymous codons. Example shown is for Proline. (**B**) Mean fold change in Co-expression Fitness (YFP) upon CFP or mCherry co-expression, normalized to the parental (high CAI) control. (**C**) Mean fold change in CFP or mCherry Expression Level relative to parental control. (**D** and **E**). Poor correlations (Pearson’s r) between fold change in Expression Level of CFP or mCherry re-codes with Co-expression Fitness. Pearson correlation coefficients were calculated from *n* = 40 codons in each plot, data points are means of *n* = 3 biological replicates.

Discrepancies in the effect between analogous re-codes of CFP and mCherry could in part be due to different amino acid composition, as the number of re-coded amino acids was not held constant between genes (**fig. S14**). We chose to re-code all instances of each amino acid to maximize the effect size, and to not limit the number of altered codons to the amino acid with the fewest instances. Most of the re-codes do not improve expression (**Fig. 4C**), which is expected since they were derived from (and normalized to) high CAI sequences that emulate highly expressed genes. CFP and mCherry re-codes correlate better in terms of Co-Expression Fitness than Expression Level, reflecting a higher degree of variability between genes in *cis* compared to *trans* effects.

Expression Level and Co-expression Fitness do not correlate well for the re-codes (**Fig. 4D and E**), indicating that while there may be general tradeoffs between expression and fitness, there are many instances where specific codons improve fitness while maintaining high expression, suggesting they possess excess translational capacity.

### Novel recoding scheme yields genes with robustly improved fitness

Next, we developed a new recoding index derived from Co-expression Fitness values for individual codons in **Fig. 4B**. We chose to focus on fitness rather than expression since our primary aim was to investigate how re-coding schemes can modulate resource competition during translation elongation. To convert the Co-expression Fitness data for CFP and mCherry re-codes into generalized codon weights, we took the Euclidean distance from the origin to the coordinates of each data point shown in **Fig. 4B** as a raw score for each sequence, where each parent codon held a normalized coordinate value of (1,1) (**fig. S15**). This method scores codons higher that improve Co-Expression Fitness for both CFP and mCherry. Similar to calculating CAI, relative adaptiveness (W_i_) scores were then determined by normalizing the raw weights from each amino acid codon set to the codon with the highest fitness (see methods and **Data S8**). We refer to this new metric as the Codon Health Index (CHI or χ).

A comparative analysis between CUB in the overall *E. coli* genome, CAI (using highly expressed genes as a reference), and χ reveals that χ favors very different codon use than CAI and discourages use of codons enriched in highly expressed genes (**Fig. 5A**). In this analysis, RSCU is calculated for a perfectly adapted gene to each of the 3 metrics being compared. The most notable differences between χ and CAI are for Arg CGT, Leu CTG, and Pro CCG. There are instances where χ and CAI do correspond well (e.g., Gly GGA, GGC, GGG), but many codons show inverse trends between the two scales. Generally, amino acids with multiple available tRNAs (including Arg, Leu, and Pro as shown in **fig. S13**) correspond with significantly larger deviations between CAI and χ (**fig. S16**), suggesting that recruitment of different tRNAs is playing a role in determining Co-Expression Fitness. Interestingly, χ favored codons do not always correspond to amino acids with multiple available tRNAs, indicating tRNA availability may not alone account for the observed effect, which could also be in part due to different translation efficiencies created by interactions of tRNA codon-anticodon pairs.

**Fig. 5.**
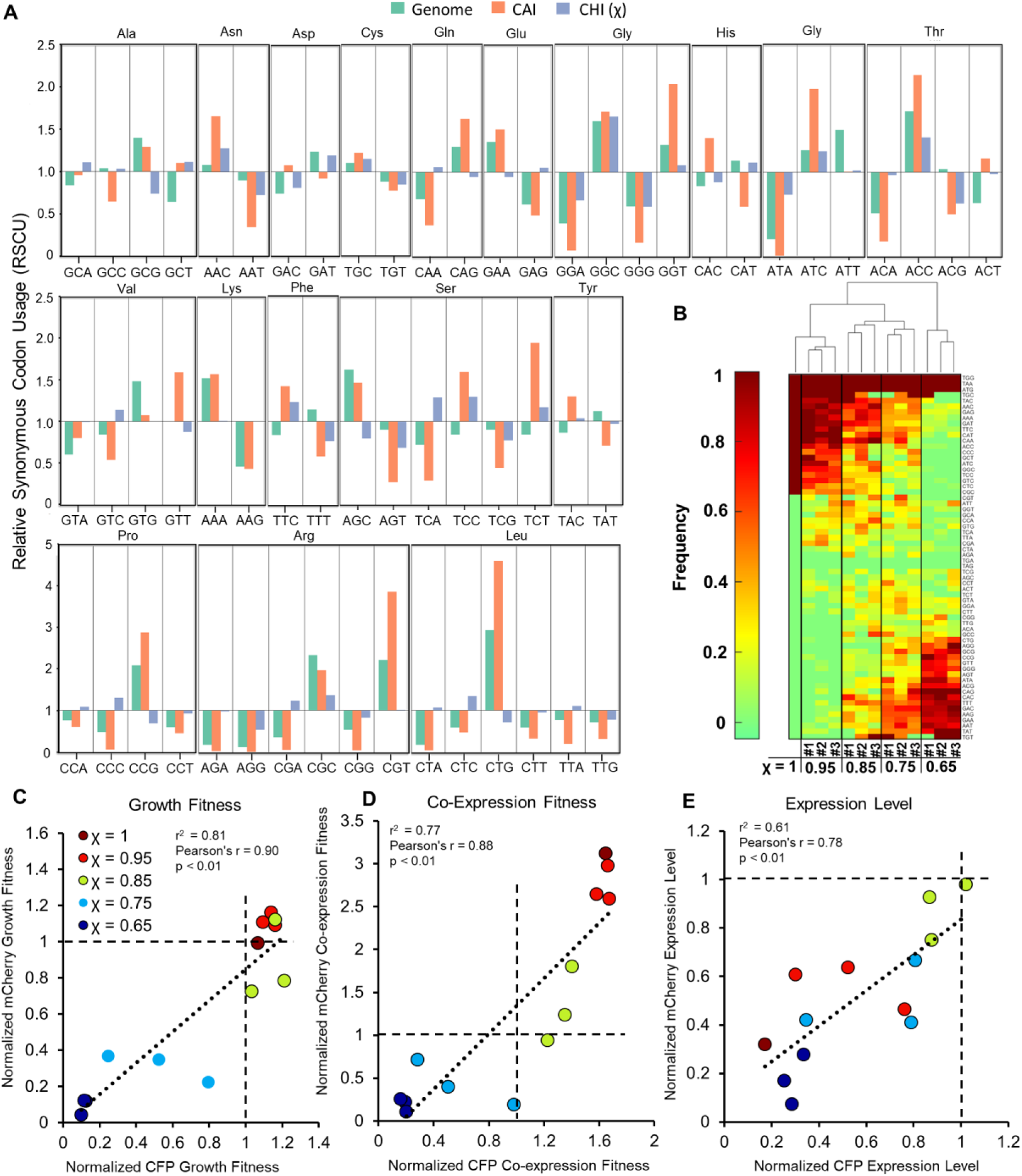
Codon Health Index (CHI, χ) used to design and test sequences for CFP and mCherry. (A) Relative synonymous codon usage (RSCU) observed in the *E. coli* genome or calculated for weighted CAI and χ scales. Calculated RSCU values represent the expected RSCU for a gene perfectly adapted to each scale. **(B)** Codon frequency of CFP or mCherry re-coded sequences using variable χ values illustrated on a clustered heat map. Three unique sequences were tested for each χ value other than for χ = 1. **(C−E).** Growth Fitness, Co-expression Fitness, and Expression Level data for CFP and mCherry re-coded using χ. In each case, results were normalized relative to the high CAI parent control. Pearson correlation coefficients and linear regression r^2^ values were calculated from *n* = 13 re-codes in each plot, data points are means of *n* = 3 biological replicates.

Utilizing the new χ weights, we created several CFP and mCherry sequences that were optimized to varying degrees (**Fig. 5B**). Specifically, we created a χ = 1, ENC = 20 sequence, along with 4 sets of 3 different sequences each holding χ constant at 0.95, 0.85, 0.75, and 0.65 for both CFP and mCherry by using a greedy algorithm (**fig. S17**). Importantly, these triplicate cases vary in primary sequence space and individual codon usage bias while maintaining consistent χ values. The lower end of the χ scale for the CFP/mCherry genes was approximately 0.6, which is dictated by the protein sequence, and lowest W_ij_ values for each set of codons (see methods). When the χ re-coded sequences were assayed for fitness and expression (**Fig. 5 C**–**E**), there was a strong positive correlation between CFP and mCherry analogous re-codes for fitness and expression, indicating that these synonymous coding schemes are a primary determinant for how a gene performs regardless of amino acid sequence. Remarkably, we also observe a strong positive correlation between χ, and both Growth Fitness and Co-Expression Fitness—indicating that the weights derived from the individual codon assay are additive to improve the fitness of various globally-recoded sequences (**fig. S18**). The benefit of high χ is consistent across different sequences, indicating the primary predictor of Co-Expression Fitness is codon use and not any other unintended sequence feature(s). The High χ sequences clearly provide reduced competition for host resources and improved fitness. The χ scale is less predictive of expression, which is expected as it was not part of the criteria used to create the codon weights. Despite this, there is a good correlation between CFP and mCherry re-coded sequences in terms of Expression Level, indicating that codon usage bias can predict expression. Importantly, there are several sequences with reduced burden that retain relatively high expression, which represents an excess translational capacity for sequences re-coded using high χ values.

Since χ was derived empirically from a specific set of conditions, we evaluated how robust the best performing re-coded sequences were at improving fitness for CFP and mCherry, and investigated their *trans*-acting effects on background gene expression. Using the CFP or mCherry χ = 0.95 #3 codon weights (**Fig. 5B**), we tested M9 minimal media, as well as 30°C and 42°C conditions. Varying nutrient availability and growth rate can change cellular resource capacity (*31*), and are thus important variables to consider when evaluating the broader utility of χ. We found that Co-Expression Fitness for the χ re-codes significantly outperformed the high CAI control under all conditions tested (**Fig. 6A, fig. S19**). Furthermore, at 42°C the mCherry re-code had higher fitness and expression than the high CAI counterpart. To confirm that Co-Expression Fitness variation was due to translational resource allocation, we performed qRT-PCR on YFP and CFP or mCherry using the same variants at the 37°C baseline condition (**Fig. 6B, fig. S19E**). These results indicate that while YFP Co-Expression Fitness is higher using χ instead of CAI, the cDNA levels are not significantly different, demonstrating that mRNA levels do not account for the observed differences in YFP protein levels. Lower cDNA levels for CFP and mCherry weak RBS controls (**fig. S19E**) are likely due to higher degradation rates of poorly translated mRNA, a phenomenon that has been well documented (*49*). We do not observe lower cDNA levels of the equivalent weak RBS construct expressed in vitro (**fig. S4**), which further supports this notion.

**Fig. 6.**
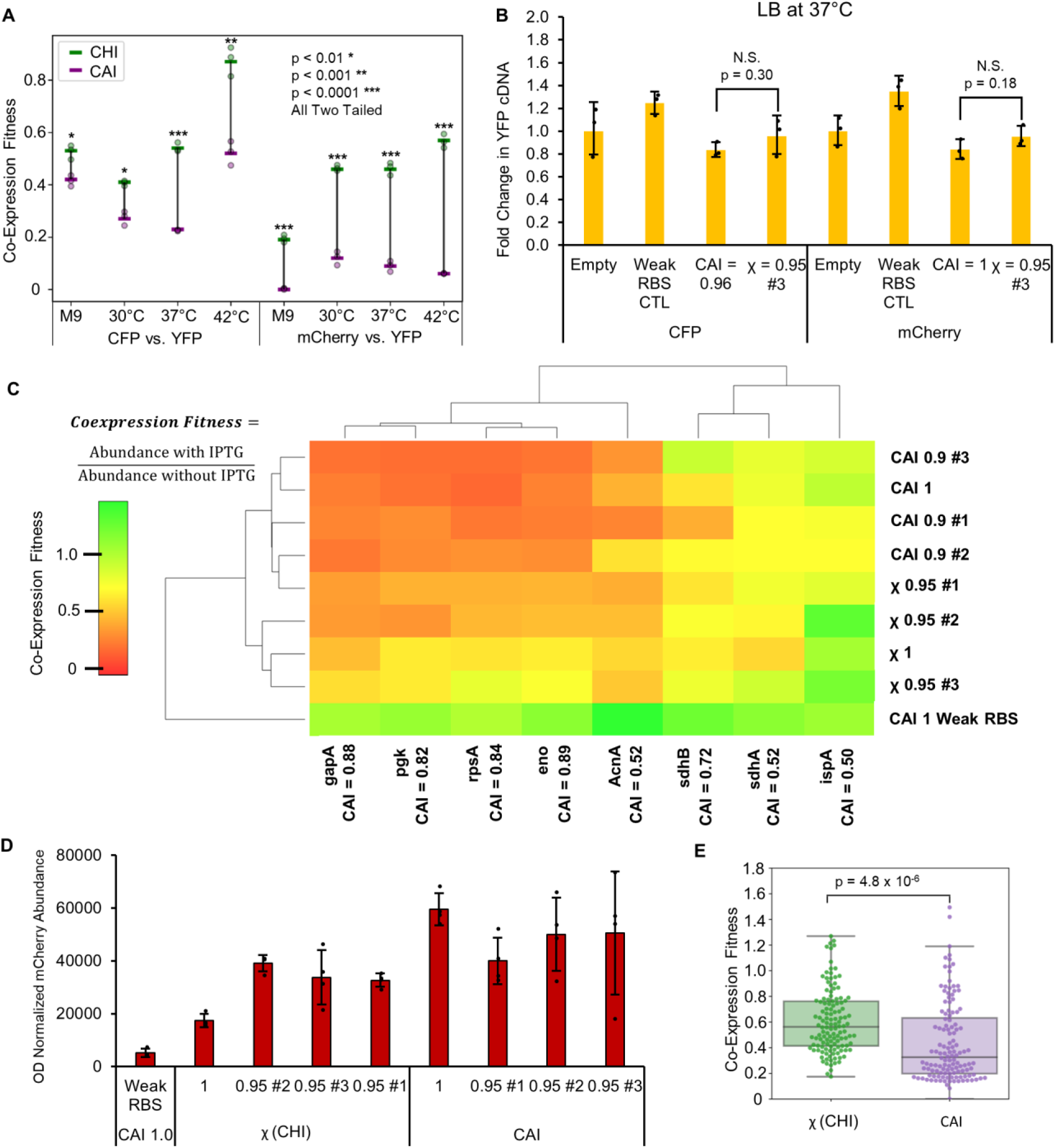
Investigating the effect of high χ sequences. (**A**) Testing the χ = 0.95 #3 sequences for CFP or mCherry compared to the high CAI controls validates that Co-Expression Fitness is robustly improved regardless of growth conditions (M9 = minimal medium, 30°C, 37°C, and 42°C indicate growth temperature in LB). *n* = 3 biological replicates of single re-coded sequences, bars represent means ± SD. (**B**) qRT-PCR results indicate that transcription of co-expressed YFP is consistent between high CAI and high χ sequences. Empty refers to an empty plasmid control. *n* = 3 biological replicates of single re-coded sequences, bars represent means ± SD. (**C**) LC-MS/MS analysis of protein Co-Expression Fitness of several *E. coli* proteins upon induction of the mCherry gene re-codes. Hierarchal agglomerative clustering was performed on the mean fitness values (*n* = 4). (**D**) Quantifying soluble mCherry abundance (*n* = 4) for the same samples as (C) with IPTG induction. Abundance was determined by quantifying AUC for a unique peptide associated with each protein using LC-MS/MS and normalized to OD600. Bars represent means ± SD (**E**) Co-Expression fitness values aggregated on a box and whisker plot for all analyzed *E. coli* proteins show a significant increase in fitness for χ re-coded variants relative to CAI re-codes (*n* = 128 for each set of combined fitness values). All p values represent two tailed t tests.

We further investigated the impact of high CAI vs. high χ re-coded sequences by running targeted proteomics on select native host (*E. coli*) proteins (**Table S1**). We calculated the OD normalized ratio of host protein abundance with or without mCherry induction as a measure of Co-Expression Fitness (**Fig. 6C**). This was done for three high CAI re-coded variants of mCherry (CAI = 0.9) and three high χ variants (χ = 0.95). Using only Co-Expression Fitness data, the high χ sequences cluster independently from the high CAI sequences, indicating that the host genes respond differently to the two types of coding schemes. Overall, expression of high χ sequences has a smaller effect on expression of housekeeping genes relative to high CAI sequences and the control sequence with a weak RBS has the best overall fitness. The secondary metabolic protein IspA is generally less affected than those involved in primary metabolism. We also quantified OD normalized soluble mCherry protein abundance (**Fig. 6D**) and found that high χ re-codes yield similar albeit slightly lower overall protein compared to CAI counterparts. Despite having similar expression level, high χ re-codes have higher Co-Expression Fitness in aggregate (**Fig. 6E**).

The utility of χ as a re-coding strategy is expected to be greatest for systems where multi-gene expression is required, and when host growth and physiological health is a priority to achieve a desired outcome. Furthermore, we expect χ to translate to new proteins other than CFP and mCherry from which it was derived. To validate χ in a biocatalysis context, we re-coded three different enzymes using high χ and CAI schemes holding the first 51 bases constant and evaluated their total activity and Co-Expression Fitness with YFP (**Fig. 7, Table S2**). We selected the *Anabaena variabilis* phenylalanine ammonia lyase (AvPAL) for conversion of L-phenylalanine to *trans*-cinnamic acid (tCA) (**7A**), *E. coli* β-galactosidase (EclacZ) for conversion of *ortho*-nitrophenyl-β-galactoside (ONPG) to *ortho*-nitrophenyl (ONP) (**7B**), and *Bacillus coagulans* L-arabinose isomerase (BcLAI) (**7C**) for conversion of galactose to tagatose. For AvPAL and BcLAI, substrates were added at the time of induction, and conversion was evaluated by measuring product concentration at the end of the assay. For EclacZ, activity level was measured by lysing cells after growth was complete, then evaluating enzyme activity. In all three cases, we observe a significant improvement in Co-Expression Fitness when re-coding with high χ over high CAI, thus confirming that χ re-coding weights extend to new proteins. Furthermore, for AvPAL and LAI we observe higher formation of tCA and D-tagatose respectively, indicating more total production of functional enzyme over the course of the assay. High activity from the weak RBS for EclacZ and BcLAI is likely because the *E. coli* strain in use has native activity for these reactions. These results confirm that χ re-codes can outperform CAI in several experimental contexts.

**Fig. 7.**
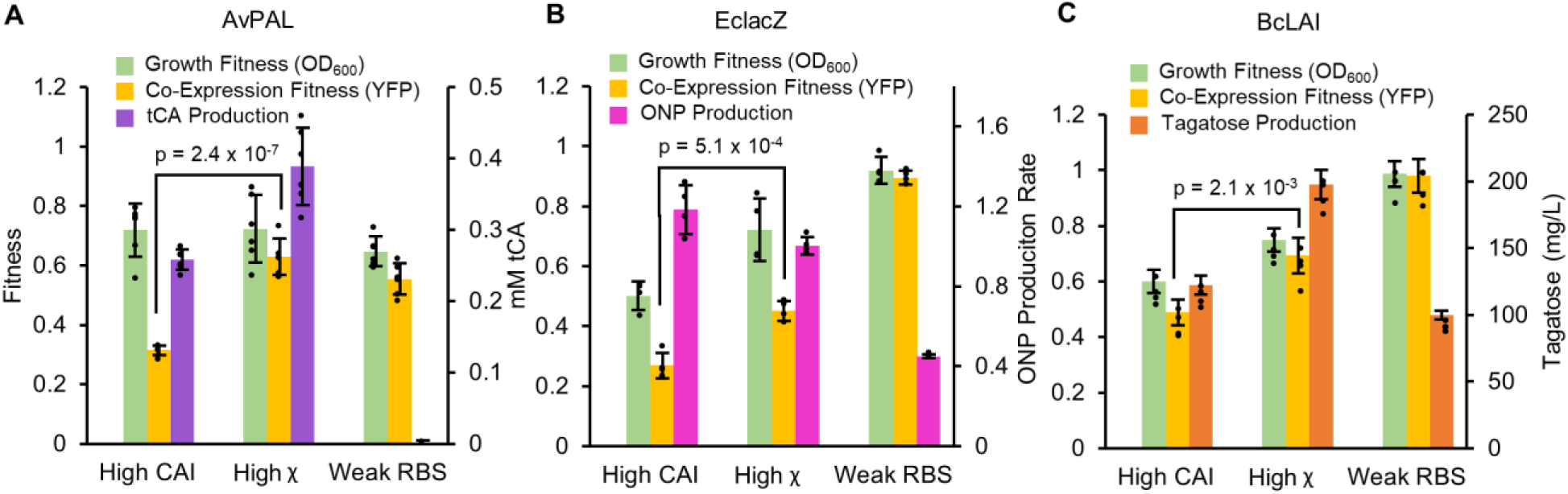
Evaluation of CHI (χ) with metabolic enzymes. (**A**) Fitness and activity data for AvPAL re-coded with high CAI or high χ. Activity is quantified by measuring endpoint accumulation of tCA. (*n* = 6) (**B**) Fitness and activity data for EclacZ re-coded with high CAI or high χ. Activity is quantified by measuring ONP production rate of endpoint lysed cultures (*n* = 4). (**C**) Fitness and activity data for BcLAI re-coded with high CAI or high χ. Activity is quantified by measuring endpoint accumulation of D-tagatose (*n* = 4). All replicates are biological replicates of single re-codes, bars represent means ± SD. All p values represent two tailed t tests.

To investigate which codon usage bias patterns for χ have the greatest contribution to Co-expression Fitness, we analyzed RSCU across all variable 59 codon dimensions (excluding stop, Trp, and Met codons) for each of the CFP and mCherry re-coded sequences (as seen in **Fig. 5B**) using PCA (**Fig. 8**). We were able to represent 46.7% of the total sequence variation in the first 3 dimensions (**fig. S20**) when analyzing the CFP and mCherry re-codes’ RSCU values along with 773 *E. coli* operons. Here again PC1 and PC2 primarily explain variation across *E. coli* sequences, but intriguingly we see a new highly orthogonal dimension in PC3 that explains variation in the χ sequences, and PC1 vs. PC3 best differentiate the χ re-coded sequences from natural *E. coli* operons. The χ sequences generally have intermediate to low values on the CAI scale with low overall CAI variation, meaning they would not have been predicted to express well using CAI (**Fig. 8A**). This is somewhat surprising given that many of the re-codes with moderate to high χ (0.8−0.95) still exhibit relatively high expression compared with the high CAI control as demonstrated in **Fig. 5E**. When mapping χ values to the data, we see that χ describes variation along PC3 very well (**Fig. 8B, fig. S21**). *E. coli* operon sequences do not vary significantly on the χ scale, implying that the re-coded sequences explore novel coding schemes orthogonal to natural sequence space. Examining the loadings for the 3 most biased natural codons, we find that the high χ sequences are using synonymous variations for Arg, Leu, and Pro that differ as expected from highly expressed genes. We conclude that competition for tRNA isoacceptors in high demand by highly expressed essential genes primarily drives competition for translation elongation resources, and avoiding specific codons that are over-represented in such native genes provides a novel strategy to improve the Co-Expression Fitness of heterologous genes.

**Fig. 8.**
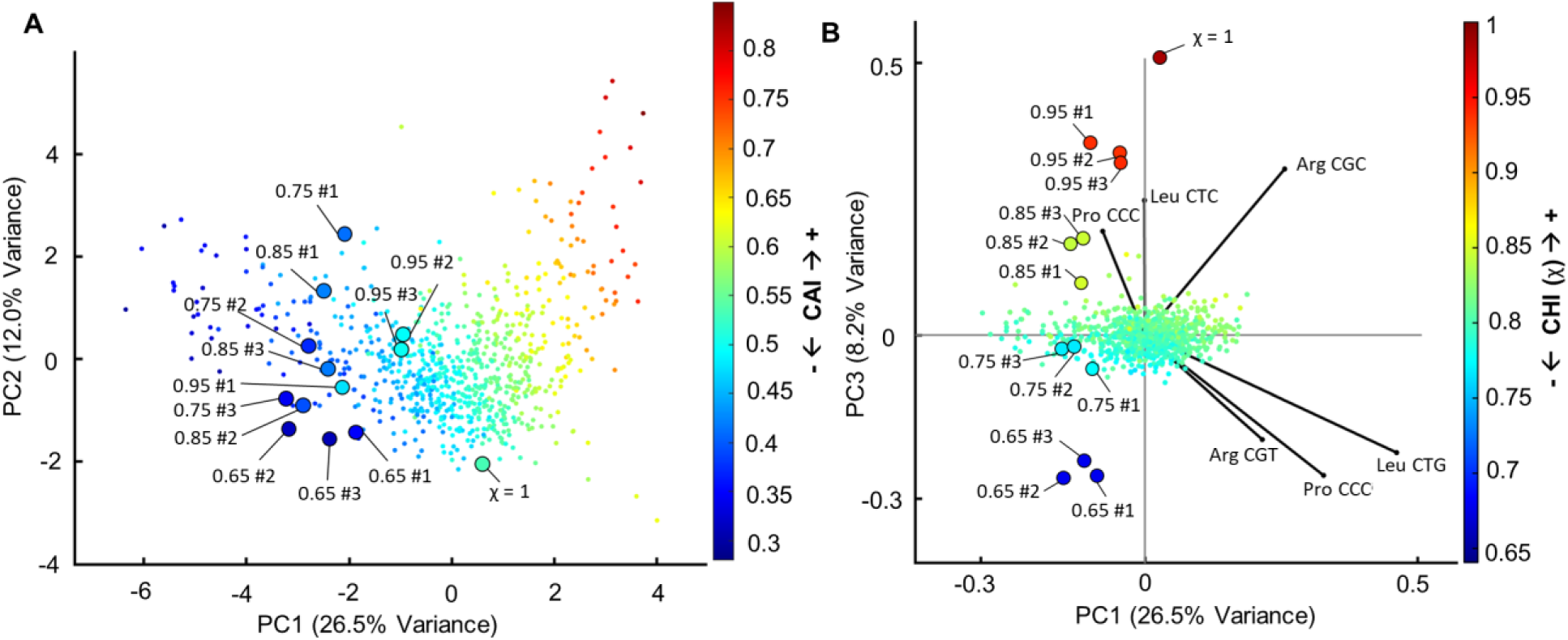
PCA analysis of χ and CAI metrics. (**A**) PCA of *E. coli* operons as well as 13 χ re-coded sequences representing RSCU for 61 codons, with CAI value mapped to individual points showing PC1 vs. PC2. (**B**) Same PCA with χ mapped to individual points instead of CAI showing PC1 vs. PC3, and displaying loading vectors for Arg/Pro/Leu codons that are driving differences between χ and CAI.

Given the breadth of existing knowledge regarding codon optimization (see (*50*)), we also evaluated how χ compares with other reported CUB strategies such as the tRNA adaptation index (tAI) (*7*) and normalized translation efficiency (nTE) (*6*). These approaches weight codons based on their co-adaptation to the tRNA pool or the tRNA supply vs. codon demand respectively. We calculated the expected RSCU of a perfectly adapted gene sequence using these various scales to assess their degree of similarity (**fig. S22**), and found that stAI (species specific TAI using *E. coli* specific weights) (*26*) correlates the closest with χ (Pearson’s r = 0.393, p = 0.002), but does not provide as much differentiation between codons available for each amino acid. We suspect the primary differentiator of the χ re-coding strategy relative to tAI or nTE is that it provides empirical insight into which specific codons have excess capacity for translation as opposed to an approach relying solely on genomic statistics and approximations. Further analysis of the χ re-coded sequences did not reveal any consistent correlation with secondary structure or GC content between CFP and mCherry re-codes, supporting the notion that specific codon use is likely driving sequence behavior (**fig. S23**). We also re-coded 10 random genes with 3 free commercial re-coding algorithms to analyze whether any of them exhibit exploration of χ related CUB strategies, and found that they generally vary along classical *E. coli* CUB and seek to adapt to host codon use without optimizing in the χ sequence space (**fig. S24**).

In theory, χ could also correlate with CUB in phages that infect *E. coli* and have co-adapted to maximize gene expression without overwhelming host resources. There have been reports of not only co-adaptation to tRNA pools (*51*, *52*), but also translational selection for CUB dissimilarity between viruses and hosts to avoid excessive competition for tRNAs (*53*). We examined codon usage in 12 common coliphages known to infect *E. coli* to examine whether CUB in such parasitic viruses may have evolved to harmonize with bacterial hosts as a means to allow better co-utilization of shared translational resources (**fig. S25**). Our analysis indicates that phage genes generally tend to avoid CUB at high values of CAI (>0.7) and exhibit a slightly higher mean χ than *E. coli* genes. This suggests that it may be more productive in the phage life cycle to avoid excessive similarity and competition with their host, but there is another unique aspect of the CUB in χ that was not strongly selected for in phages. It is possible that the translational resource demand from an overexpressed protein on a multi-copy vector is higher than natural genes have encountered, and is thus under a higher level of translational selection resulting in novel types of advantageous CUB reflected by χ that cannot be inferred from natural sequence space.

## DISCUSSION

Protein synthesis is one of the most resource intensive cellular processes, which has yielded significant CUB observed in nature, especially in single cellular microorganisms often used as expression hosts (*38*). Most conventional codon optimization strategies operate under the key assumption that translational selection in naturally evolved systems provides CUB that is relevant for the overexpression of heterologous genes. This may be partially true, but realistically, the overexpression of genes can push host resource demand beyond levels required for native gene expression (*54*), resulting in translational selective pressures that organisms haven’t evolved with. Protein expression must also be considered in the context of increasingly complicated engineered systems, and often in synthetic biology and metabolic engineering efforts, overexpression is not nearly as important as reliable and predictable gene expression and host fitness (*55*). Here we have demonstrated both in vitro and in an *E. coli* model that translation elongation can limit protein expression, and often has profitable or catastrophic consequences on system-wide resource availability.

In our TxTL assay, we found that proteins coded with similar CAI compete for the same resources, and re-coded genes can reduce such competition. Consequently, high CAI sequences are likely ribosome-limited, demonstrating reduced synthesis rates that are also highly sensitive to competition. In certain cases, low CAI genes are monopolistic or anti-competitive with free ribosomes and are thus insensitive to increased demand from high CAI sequences, albeit at the expense of overall resources. Theoretical frameworks have been well established to explain how resource limited translation can lead to the sequestration of ribosomes, but these studies generally rely on ribosome footprinting data (*43*) and tRNA copy number (*6*, *7*) to infer codon elongation times, which are indirect measurements of ribosome flux on a given mRNA. While insightful, the accuracy of such methods has been questioned given they are indirect measurements of specific codon elongation rates (*56*).

Our experimental approach using an *E. coli* model demonstrates the sensitivity of system resources at individual codon resolution and reveals key differences between the optimal CUB for highly expressed native genes vs. overexpressed proteins. Several previous studies have investigated CUB using randomized libraries that fail to thoroughly explore the vast sequence space available when re-coding a gene (*57*). Such randomized sequences will generally regress to intermediate RSCU values for each codon, and rarely sample the extremities of the sequence space available (**fig. S26**). Similar to our study, others have found that matching codon bias closer to overall host codon use rather than to highly expressed genes can improve gene fitness (*37*), but these results have been limited to sequence designs emulating natural codon use. By systematically re-coding individual amino acids to each alternate codon in multiple proteins, we have methodically investigated how individual codons contribute to gene Expression Level and Co-Expression Fitness at further extremities of the theoretical design space than have been previously explored, revealing non-natural codon bias that has significant fitness benefits (**Fig. 8**), and is largely orthogonal to natural CUB. The avoidance of codons with very high CUB in native essential genes (e.g., for Arg/Leu/Pro), and the preference for specific alternative synonymous codons, is a novel driver of reduced genetic burden.

We used individual codon sensitivity data to create a new re-coding strategy that optimizes for fitness (CHI or χ) and demonstrate how the new codon weighting method enables the creation of unique CUB strategies that are not represented naturally in *E. coli*. Using PCA for dimensional reduction, our methodology reveals how sequences with identical CAI scores can still exhibit distinct variations in CUB that result in different phenotypes, namely improvements in Co-Expression Fitness. Remarkably, globally re-coded sequences were found to have predictable phenotypes informed from the additive effects of individual codon use, allowing us to leverage a relatively small dataset to predict phenotypes in a vast sequence space. Using χ we were able to demonstrate its utility beyond model proteins for three unique enzymatic contexts. While global sequence characteristics including GC content, structure, and a variety of sequence motifs are all known to contribute to protein expression (*2*), our results suggest that codon bias is a strong predictor of both protein expression and fitness and can be optimized independently of the UTRs or 5’ coding sequence. Accordingly, our data should be useful in fitting and refining computational models (*23*, *38*, *40–42*, *58*, *59*) in the context of protein expression and resource competition. An analysis of *E. coli* phage CUB reveals that while parasitic organisms may avoid over-use of preferred host codons, a concept that has been recently suggested (*53*), the demands of heterologous gene overexpression and resulting selective pressures are likely to have different resource demands than those of viruses, and thus may have overlapping yet still largely distinct CUB fitness landscapes.

The data-informed strategy in this study represents an approach that could be extended to other microbes including eukaryotic systems, where ongoing controversy over the impact CUB has on host-gene fitness has been unresolved (*60–64*). Since χ was derived empirically, it is unlikely to have direct utility outside of an *E. coli* context, however the approach could be applied to other organisms, especially where significant natural codon bias exists in highly expressed essential genes. While our initial study included 2 proteins (CFP and mCherry) with very different amino acid sequences, measuring Expression Level and Co-Expression Fitness for additional proteins could further refine χ, and provide additional insight for maximizing expression and fitness together. The new χ metric is more predictive of *trans* effects (Co-expression Fitness) than *cis* effects (Expression Level), thus further optimization of translation initiation and CUB that maximizes both expression and fitness is an interesting future objective. The observation that there are several sequences with relatively high expression and high fitness illustrates there are solutions to co-optimize both genetic traits. In practice, re-coding genes with high CAI will often lead to higher expression with low overall fitness, but re-coding with high χ values (between 0.9−0.95) should provide reasonably high expression with more orthogonal resource demands.

Similar data sets could also be collected for any organism where protein expression is feasible, which could also provide insights into how species differ in the role CUB plays regarding resource allocation. It is possible that with more inter-species data, organism specific χ weights could be predicted *a priori* based on the avoidance of codons overrepresented in host genes.

Practically, this study should improve the predictability and robustness of genetic engineering by enabling the co-optimization of gene expression and fitness, especially for multi-gene expression systems.

## MATERIALS AND METHODS

### Equations used to assess codon usage bias

We calculated codon adaptation following the classical method reported originally by Sharp and Li (*19*). This method relies on first calculating relative synonymous codon usage (**RSCU**) in a genetic sequence, which is defined by **Equation 1**:

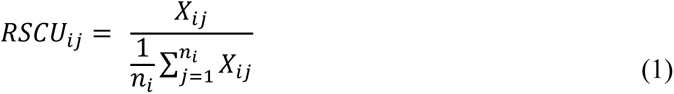

RSCU calculates the observed frequency of codon **j** belonging to amino acid **i** divided by expected frequency, where **X** is the number of occurrences for codon **j** in a given sequence. The expected frequency is simply the number of occurrences for any codon belonging to amino acid **i**, divided by the number of codons (**n**) available for that particular amino acid. RSCU is used instead of raw frequency values to normalize observed codon frequency based on the total codons available. An RSCU value < 1 indicates bias against the codon, while an RSCU value > 1 indicates a bias toward the codon, and RSCU = 1 indicates no bias. The RSCU values for each codon can be used to calculate relative adaptiveness (**W**), which is defined by **Equation 2**:

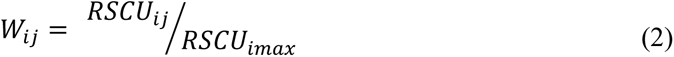

Relative adaptiveness is the RSCU for a codon **j** belonging to amino acid **i** divided by the RSCU for the codon in the set for amino acid **i** with the highest RSCU value (imax). In other words, W gives a value of 1 for codons in a target sequence that match the frequency of the most common codon in a reference sequence. W values are used in calculating the codon adaptation index (**CAI**) defined by **Equation 3**:

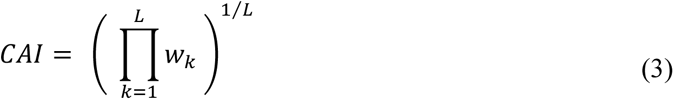

CAI is the geometric mean of the W values for each codon in a given sequence containing L codons. Importantly, the reference sequence(s) and calculated RSCU values that W values are derived from can be from any source. Unless otherwise indicated, in this study, CAI always refers to W values for a set of highly expressed set of *E. coli* genes (in K12 MG1655). Alternatively, CAI can be computed based on W values for CUB across the entire genome, sTAI weights (*26*), or χ weights (**See Data S8 and Data S9 for W values used in various calculations**). Normalized translational efficiency (nTE) was calculated as previously described (*6*) by taking the ratio of species specific TAI weights for *E. coli* (*26*) (supply) vs. the codon use across the *E. coli* transcriptome (demand) defined by **Equation 4**:

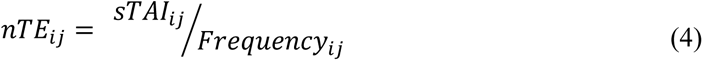

The nTE_ij_ values are analogous to W_ij_ values for the calculation of nTE, which proceeds the same as for CAI by taking the geometric mean across a sequence (as in equation 3). In this study, nTE was calculated using genomic codon frequency as opposed to codon use (originally defined as codon occurrence multiplied by RNA transcript abundance), as the two were found to be highly correlated (**fig. S27**). Lastly, the effective number of codons (**ENC**) is often used as a measure of codon bias in a sequence, and is calculated using **Equation 5**:

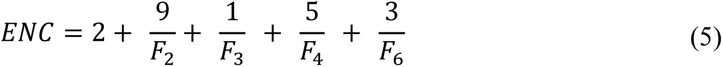

ENC can take a value from 20, in the case of extreme bias where one codon is exclusively used for each amino acid, to 61 when the use of alternative synonymous codons is equally likely. The value F is the average probability that two randomly selected codons for an amino acid with n number of synonymous codons will be identical (*65*).

### Data sources used in analysis

Genomic codon usage for *E. coli K12* MG1655 and *E. coli* MRE600 were assessed by analyzing codon bias from published annotated genomes obtained from NCBI under the accession numbers NC_000913.3 and CP014197.1 respectively using MATLAB. Phage analysis was done with annotated phage genomes from NCBI, and accession numbers are listed in **fig. S25**. Exact codon frequencies and relative adaptiveness values (W) used in this study for calculating CAI in reference to highly expressed genes CUB, entire genome CUB, sTAI, or nTE, can be found in **Data S9**. Calculations for nTE were aided by transcript abundance from a publically available dataset (GEO accession GSE59377) (*66*). The W values for χ and associated information from the study can be found in **Data S8**. W values for highly expressed genes were originally downloaded online from GenScript, and were cross referenced to published values (*48*). The sTAI codon weights were downloaded online from a publically available database (http://tau-tai.azurewebsites.net/) (*26*). The tRNA copy numbers referenced in this study (**fig. S13**) were downloaded from the Genomic tRNA Database (http://gtrnadb.ucsc.edu/) (*67*).

### Ribosome flow model

The implemented ribosome flow model (RFM) (**fig. S1**) was adapted from Zur et al. using open source Matlab® code (*42*). In this model, an mRNA is divided into n number of chunks, where each chunk is 9 codons (27 bases), approximately the footprint of an *E. coli* ribosome. Translation time of each chunk is based on local λ, which is a sum of the individual times it takes to translate each codon in a chunk. Codon times used are available in **Data S1**. Ribosome collisions are also accounted for in the model as a function of the ribosome density in adjacent positions. In this model, the protein production rate is the rate of translation of the final position on the mRNA. For this application, steady state ribosome densities were computed for CFP and YFP re-coded to use preferred (high CAI) or rare (low CAI) codons. To demonstrate the relationship between initiation rate and translation rate for different sequences, steady state protein production rates are calculated for different initiation rates.

### Gene design and re-coding

All genetic re-coding designs and analysis were executed in Matlab® using custom functions. Code is made available online at https://github.com/nair-lab. A full list of amino acid and DNA sequences used in this study can be found in **Data S10.** CFP and YFP were initially cloned through site directed mutagenesis of an existing super-folder GFP protein based on previously reported sequences. (*44*, *45*) For the systematic analysis of codon use design, CFP or mCherry were re-coded starting from highly biased sequences using the most preferred codon for each amino acid (CAI = 1 and ENC = 20), not taking into account the first 17 codons. The first 17 codons were held constant for all re-codes and were based on previously used sequences that functionally expressed well. A Matlab® script was then used to systematically design sequences where every instance of an amino acid was mutated to a single alternate synonymous codon. In the design of sequences with novel re-coding schemes, a greedy algorithm was used (**fig. S17**), that functions by randomly mutating a codon to a synonymous alternative, then evaluating whether the new sequence is closer to the target CAI (or in this specific instance χ value). To re-code CFP and mCherry to a desired χ value, a starting sequence was first randomized to ensure there was no initial bias, and then the algorithm was followed to the target χ value. We generated several unique output sequences with the same χ value but different coding sequences, then selected 3 sequences for each value of χ tested making sure they were substantially different from each other based on hierarchal clustering done in Matlab®.

### Plasmids and strain construction

Most plasmids were cloned from existing vectors with restriction enzyme sites already present (**fig. S7, S28, S30 Data S10**), which also contained 5’ and 3’ UTRs. Genes were generally custom ordered synthesized as full length double stranded DNA fragments with AarI restriction sites on the 5’ and 3’ termini. A type IIS restriction enzyme cloning approach with AarI was used to insert synthesized double stranded DNA gene fragments into the desired vector. In the case of pBAC-RARE2 (**fig. S7, Data S10**), the rare tRNA genes were amplified from a commercially available vector called pLysSRARE2 available from Rosetta 2(DE3)pLysS Competent Cells (Novagen).

The tRNA genes were cloned into a bacterial artificial chromosome (BAC) present at approximately one copy per cell, and expressed under their native promoters. All constructs were sequence verified from clonally pure DNA using Sanger sequencing across the gene and UTRs. The screening strain used to assess Co-Expression fitness was engineered from *E. coli* K12 MG1655 (CGSC#: 6300) modified with endA and recA gene knockouts (F- λ- rph-1 ΔendA ΔrecA). The YFP reporter was integrated in an intergeneic region (∼3,938,000 bp) between the rsmG-atpI genes using λ-Red based homologous recombination of the YFP CAI = 0.96 sequence, which was under the control of a strong constitutive promoter (FAB46) and RBS (BCD7) based on a previous study, (*46*) and a 5’ insulator and 3’ terminator (**fig. S29**, **Data S10**). The method of integration and marker excision method has been previously reported (Datsenko and Wanner) (*68*). Briefly, a linear cassette consisting of the gene, UTRs, and an attached kanamycin resistance marker was amplified by PCR with ∼500bp of homology to the desired locus on either end.

Chromosomally integrated clones were identified by colony PCR and sequence verified via Sanger sequencing of the PCR product including several hundred bases of chromosomal DNA and the entire integrated heterologous expression cassette. Sequence verified clones had the integrated kanamycin marker removed through the previously described FLP-FRT site specific recombinase method and were again Sanger sequenced for final verification.

### in vitro transcription-translation (TxTL) assay

The TxTL assay was carried out using the NEB PURExpress® kit (E6800). This assay relies on T7 polymerase, and consists of purified reconstituted components. Accordingly, CFP, YFP, and mCherry expression cassettes were first cloned into a pBAC vector with a T7 promoter and strong RBS (BCD7) (**fig. S30 A** and **B, Data S10**). The genes were also flanked by an insulator and terminator sequence on the 5’ and 3’ UTR respectively. Once clonally pure and sequence verified, expression cassettes were amplified by PCR (from the beginning of the insulator to end of the terminator) and normalized in concentration using UV-vis spectroscopy at λ = 260nm. A master mix was first prepared according to the PURExpress® published protocol, which was kept on ice until use. Reactions were scaled down to 5 µL final volume and carried out in Corning® low volume 384-well white flat bottom polystyrene TC-treated microplates (part # 3826). Reactions were initiated by the addition of DNA using a multi-channel pipette (n=2 per condition), followed by immediate transfer to a Tecan Infinite® M1000 microplate reader. A DNA concentration of 20ng/µL each was found to generally maximize competition between two genetic cassettes (**fig. S30 C** and **D**). Assays were run for 2.5hr. at 37°C with fluorescent reads every 5 minutes of each protein being analyzed (CFP: Ex. 435nm, Em. 470nm, YFP: Ex. 510nm, Em. 530nm, mCherry: Ex. 585nm, Em. 612nm). Reported reaction rates reflect the maximum rate observed for each individual replicate, which often occurred between 1-2 hours of incubation.

### Quantitative reverse transcription PCR (qRT-PCR)

To measure the relative mRNA levels for different samples, qRT-PCR was carried out for in vitro samples gathered in TxTL reactions, as well as for in vivo samples gathered from bacterial cells. For in vitro samples, TxTL reactions were stopped after 2.5hr., then directly treated with DNAse I-XT (NEB M0570) at 37°C for 30 minutes to remove DNA template, then RNA was purified using a Monarch® RNA Cleanup Kit (NEB T2030). For in vivo samples, RNA was extracted first using a Monarch® Total RNA Miniprep Kit (NEB T2010) with on-column DNAse digestion. cDNA was synthesized using a LunaScript® RT SuperMix Kit (NEB E3010), and real time PCR was run on a Bio-Rad CFX96 Touch Real-Time PCR Detection System using an iTaq Universal SYBR Green Supermix (BioRad #1725121). In vitro samples analyzed with or without reverse transcriptase showed a maximum of 5.6 x 10^-5^ percent signal from un-degraded DNA. In vivo samples analyzed with or without reverse transcriptase showed a maximum of 6.34 percent signal from un-degraded DNA. Real time quantification conditions were optimized, and standard curves were run for each cDNA species to measure reaction efficiency, as well as additional reactions to confirm primer specificity (**Data S11**). cDNA from TxTL reactions were analyzed using standard curves for absolute quantification of cDNA concentration, while cell based in vivo samples were quantified using the 2^−ΔΔCt^ method with *rrsA* as a housekeeping gene.

### in vivo fitness and expression assay

To assess Co-Expression Fitness, Growth Fitness, and Expression Level, sequence verified plasmid constructs were transformed into *E. coli* K12 MG1655 (F- λ- rph-1 ΔendA ΔrecA) with the chromosomally integrated YFP reporter. Unless noted otherwise, overexpressed proteins were under control of the Trc promoter with a strong RBS (BCD7) (**Data S10)**. 3 individual transformants were isolated and grown overnight in 400µL LB broth (BD Difco^TM^) with selective antibiotic at 37°C in 96 deep well plates (Greiner Bio-One MASTERBLOCK®, 96 Well, 2 ML Item: 780270) for 24 hr. Cultures were then split and diluted 1:40 into LB broth with selective antibiotic and with or without 500µM inducer (IPTG) in black 96 well clear bottom micro-titer plates (Thermo product: 165305). Plates were incubated for 8 hours with shaking at 37°C in a Tecan Infinite® M1000 microplate reader with monitoring every 5 minutes for OD600, as well as fluorescence (CFP: Ex. 435nm, Em. 470nm, YFP: Ex. 510nm, Em. 530nm, mCherry: Ex. 585nm, Em. 612nm). Data were analyzed by comparing independent induced vs. uninduced cultures in terms of fluorescence and growth. To account for lag phase and differences in rates within a single term, the background subtracted area under the curve (AUC) was used for each respective signal using a Matlab® numerical integrator. The timespan evaluated was bounded by the time it took any sample to reach the upper limit of detection for fluorescence, which often took between 4-6 hours. In most cases, the mean of 3 replicates was compared (fold change) relative to a control sequence (e.g., the high CAI starting sequence).

### in vivo functional enzyme assays

Functional enzyme assays for AvPAL, BcLAI, and EclacZ were carried out similarly to those for the in vivo fitness and expression assay with slight modifications. Cells were grown overnight for 24hr. in pre-culture and then split and diluted 1:40 into different media with selective antibiotic and with or without 500µM inducer (IPTG) in black 96 well clear bottom micro-titer plates. For AvPAL activity, overnight pre-cultures and production cultures were carried out in modified media consisting of 137 mM NaCl, 2.7 mM KCl, 10 mM Na2HPO4, 2 mM KH2PO4, 2 mM MgSO4, 2g/L glucose, 20 mM NH2SO4, 30mM L-phenylalanine, 5g/L casamino acids (Bacto), pH 7.4. After 6 hours monitoring growth and YFP fluorescence, cells were pelleted by centrifugation and tCA was measured by taking absorbance at 290 nm by UV-vis spectroscopy and fitting to a standard curve. For BcLAI activity, the standard in vivo fitness and expression assay was used with LB supplemented during the production culture with 10g/L D-galactose.

Note that growth and product formation data were collected from *E. coli* K12 MG1655 with additional galK and galM gene deletions (F- λ- rph-1 ΔendA ΔrecA ΔgalK ΔgalM) for BcLAI only, while Co-Expression Fitness data were still collected from the standard in vivo fitness and expression assay reporter strain (with integrated YFP). After 6 hours monitoring growth or YFP fluorescence, cells were pelleted by centrifugation and supernatants were analyzed using LC/MS with HILIC chromatography to measure D-tagatose formation with a standard curve. For lacZ activity, the standard in vivo fitness and expression assay was used without modifications. After 6 hours monitoring growth and YFP fluorescence, 10µL of cells were diluted into 90µL BugBuster® (Millipore) reagent and lysed for 10 minutes. Lysed cells were then diluted with 900µL of saline solution, then 10µL of diluted lysate was added to 90µL of substrate solution, consisting of 60 mM Na_2_HPO_4_, 40 mM NaH_2_PO_4_, 1 mg/mL o-nitrophenyl-β-D-Galactoside (ONPG), 2.7 μL/mL β-mercaptoethanol, and 2mM MgSO4. Reaction rate was monitored online by absorbance at 420nm by UV-vis spectroscopy, and activity was quantified by taking the slope of the resulting linear response.

### LC/MS detection of proteins

Cell based samples were first analyzed by measuring OD_600_ and were then prepared by centrifugation of 100-200 µL culture. The supernatant was discarded, and pellets were resuspended in BugBuster® protein extraction reagent (Millipore). Protein from the lysed cell suspension was then extracted by chloroform/methanol precipitation. Extracted protein was resuspended in 50mM ammonium bicarbonate in 10% acetonitrile with 1mM CaCl_2_. For TxTL samples, they were immediately resuspended in ammonium bicarbonate solution without protein extraction. Samples were then digested using sequencing Grade Modified Trypsin (Promega) for 12 hours at 37°C. Reactions were terminated by the addition of 1M formic acid, and then analyzed by LC/MS. Quantitative targeted proteomics were carried out using an Agilent 6470 triple quadrupole mass spectrometer with reverse phase UPLC chromatography. Samples were run on an Agilent Zorbax Eclipse Plus C18 column (2.1 x 150 mm) using a standard gradient with 0.1% Formic acid in water/acetonitrile. Representative ions for each peptide/protein were monitored for abundance and area under the curve was quantified for each peptide species (**fig. S31**).

### Derivation of CHI (χ)

The codon health index, (abbreviated CHI represented by the Greek letter χ) was derived empirically using data from **Fig. 4B**. Each data point (***p***) representing normalized Co-Expression Fitness for every codon re-coded in CFP and mCherry was quantified by taking the Euclidean distance (***d***) from the origin (***o***) to the coordinates of each data point. This is described by **Equation 6**, where **c** is the normalized Co-Expression Fitness for CFP, and ***m*** is the normalized Co-Expression Fitness for mCherry:

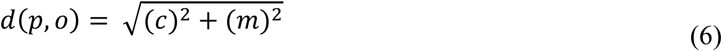

The χ scale is thus entirely based on experimental data quantifying the impact of each individual codon on Co-Expression Fitness, and weights codons higher when they similarly improve Co-Expression with CFP and mCherry overexpression. In the same way that CAI is calculated, relative adaptiveness (Wi) scores were determined for every codon by normalizing the raw Euclidean distances from each amino acid codon set to the codon with the highest fitness (as in **Equation 2**). Weights were then used to calculate χ from gene sequences similar to CAI according to **Equation 3**. Distance values are given in **Data S8**, and visualized in **fig. S15**.

### Additional data analysis

Except in the case of measured reaction rates, all data were collected from distinct samples. Mean, standard deviation, linear regression, correlation analysis, dimensional reduction, and associated statistics were calculating using built in functions in Matlab® or Microsoft Excel. Error bars in all plots represent standard deviation. Principal component analysis and hierarchal clustering were always carried out on an m x n matrix of RSCU values with codons in 61 rows and n number of gene sequences in columns. For RNA folding calculations, the minimum free energy was calculated for sequences using the Vienna RNAfold Version 2.5.1 software (*69*).

## GLOSSARY

RSCU: Relative synonymous codon usage
Wij: Relative adaptiveness (weight)
CAI: Codon adaptation index
ENC: Effective number of codons
CUB: Codon usage bias
TAI: tRNA adaptation index
sTAI: Species-specific tRNA adaptation index
nTE: Normalized translational efficiency
RFM: Ribosome flow model
CFP: Cyan fluorescent protein
YFP: Yellow fluorescent protein
TxTL: in vitro transcription-translation
UTR: Untranslated region
AUC: Area under the curve
Fitness: Performance of induced culture ÷ Performance of uninduced culture
Growth Fitness: AUC of growth curve (induced) ÷ AUC of growth curve (uninduced)
Co-Expression Fitness: AUC of YFP fluorescence (with induced CFP or mCherry) ÷ AUC of YFP fluorescence (with uninduced CFP or mCherry)
Expression Level: AUC of fluorescence from induced overexpressed protein (CFP or mCherry)
PCA: Principal component analysis
CHI (χ): Codon health index
MFE: Mean free energy
qRT-PCR: Quantitative reverse transcription PCR
BAC: Bacterial artificial chromosome
AvPAL: Phenylalanine ammonia lyase enzyme from *Anabaena variabilis*
BcLAI: L-arabinose isomerase enzyme from *Bacillus coagulans*
EclacZ: β-galactosidase enzyme from *E. coli*
tCA: *trans*-cinnamic acid
ONPG: *ortho*-Nitrophenyl-β-galactoside
ONP: *ortho*-nitrophenyl

## Author contributions

A.M.L. performed the experimental work and data analysis. A.M.L. and N.U.N. conceived the study, planned the experiments, and wrote/edited the manuscript.

## Data and materials availability

Additional data are available upon request. Additional supplementary Matlab® code can be found at https://github.com/nair-lab/CHI.

