## Supplementary Information for "Specific Codons Control Cellular Resources and Fitness"

**This PDF file includes:**

Figs. S1 to S31
Tables S1 and S2
Captions for Data S1 to S11
References (*6*, *7*, *41*, *43*, *48*, and *66*)

**Other Supplementary Materials for this manuscript include the following:**
Data S1 to S11

**KEYWORDS:** Codon bias; resource competition; tRNA; protein expression; genetic burden; translation

**GLOSSARY**

- **RSCU**: Relative synonymous codon usage
- **$W_{ij}$**: Relative adaptiveness (weight)
- **CAI**: Codon adaptation index
- **ENC**: Effective number of codons
- **CUB**: Codon usage bias
- **TAI**: tRNA adaptation index
- **sTAI**: Species-specific tRNA adaptation index
- **nTE**: Normalized translational efficiency
- **RFM**: Ribosome flow model
- **CFP**: Cyan fluorescent protein
- **YFP**: Yellow fluorescent protein
- **TxTL**: in vitro transcription-translation
- **UTR**: Untranslated region
- **AUC**: Area under the curve
- **Fitness**: Performance of induced culture ÷ Performance of uninduced culture
- **Growth Fitness**: AUC of growth curve (induced) ÷ AUC of growth curve (uninduced)
- **Co-Expression Fitness**: AUC of YFP fluorescence (with induced CFP or mCherry) ÷ AUC of YFP fluorescence (with uninduced CFP or mCherry)
- **Expression Level**: AUC of fluorescence from induced overexpressed protein (CFP or mCherry)
- **PCA:** Principal component analysis
- **CHI (χ)**: Codon health index
- **MFE**: Mean free energy
- **qRT-PCR:** Quantitative reverse transcription PCR
- **BAC:** Bacterial artificial chromosome
- **AvPAL:** Phenylalanine ammonia lyase enzyme from *Anabaena variabilis*
- **BcLAI:** L-arabinose isomerase enzyme from *Bacillus coagulans*
- **EclacZ:** β-galactosidase enzyme from *E. coli*
- **tCA:** *trans*-cinnamic acid
- **ONPG:** *ortho*-Nitrophenyl-β-galactoside
- **ONP:** *ortho*-nitrophenyl

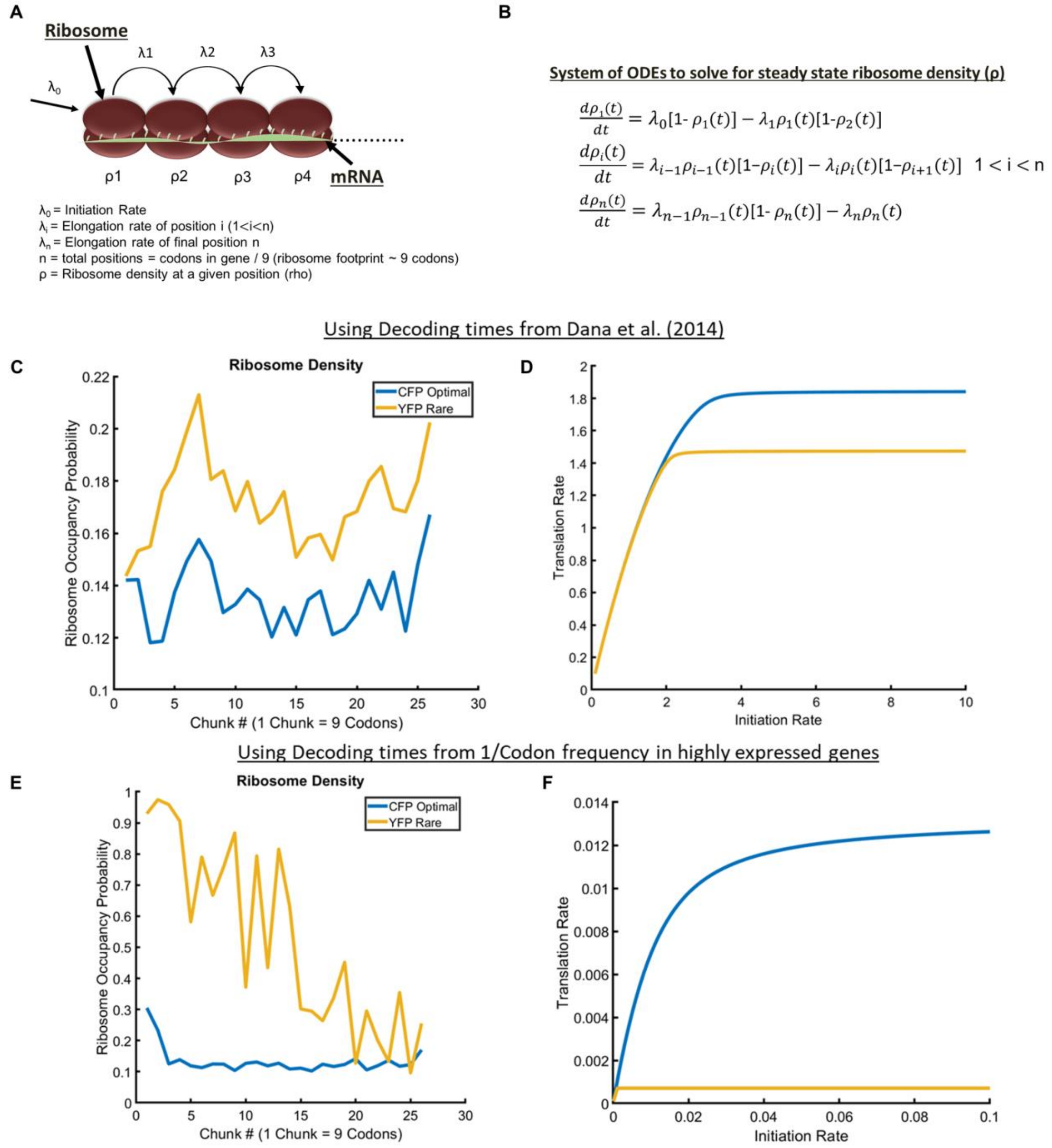


**Fig. S1.**

**Ribosome flow model (RFM) of CFP and YFP.** (**A**). Depiction of the RFM used to evaluate steady state ribosome occupancy probabilities and protein translation rates adapted from Reuveni et al. (*41*). For extensive details on the RFM, please refer to the original publication. In this model, ribosomes initiate translation according to the $\lambda_0$ rate, and then proceed to subsequent positions based on the sum of elongation times for 9 individual codons, which corresponds to the approximate footprint of a ribosome. (**B**) Steady state ribosome occupancies are calculated numerically in Matlab® by solving a system of ODEs, which varies depending on the length of the gene. Protein translation rate is the rate of translation of the final position, which is simply the ribosome density (rho or ρ) multiplied by the elongation rate at the final position ($\lambda_n$). (**C**) Calculated steady state ribosome occupancy values for codon optimized CFP

(CAI = 1 with reference to highly expressed *E. coli* genes) vs. rare codon rich YFP (CAI = 0.16 with reference to highly expressed *E. coli* genes) using codon elongation times from Dana et al. (*43*) illustrates higher occupancy on the rare codon rich sequence. An initiation rate of $1^{s-1}$ was used to generate all the ribosome density plots. (**D**) Calculated translation rate vs. initiation rate for the optimal CFP vs. rare codon YFP indicates that poorly optimized sequences become elongation limited at lower translation initiation rates. (**E** and **F**) Since exact elongation times have only been estimated and reported variations in elongation time are relatively small, the analysis was repeated using codon elongation times with more extreme variation. Here, the analysis was repeated using 1/codon frequency in highly expressed *E. coli* genes as an analog of codon elongation times (see Supplementary Data S1 for details on codon times). In this case, elongation times are more extreme for rare codons, and cause the YFP sequence to be largely insensitive to translation initiation rate due to severe elongation limitation. These results illustrate that while genes with rare codons may sequester more ribosomes and limit system resources, they are predicted to vary less in expression, or be more insulated from variation in ribosome availability, since translation initiation does not limit their expression as much as codon-optimized genes.

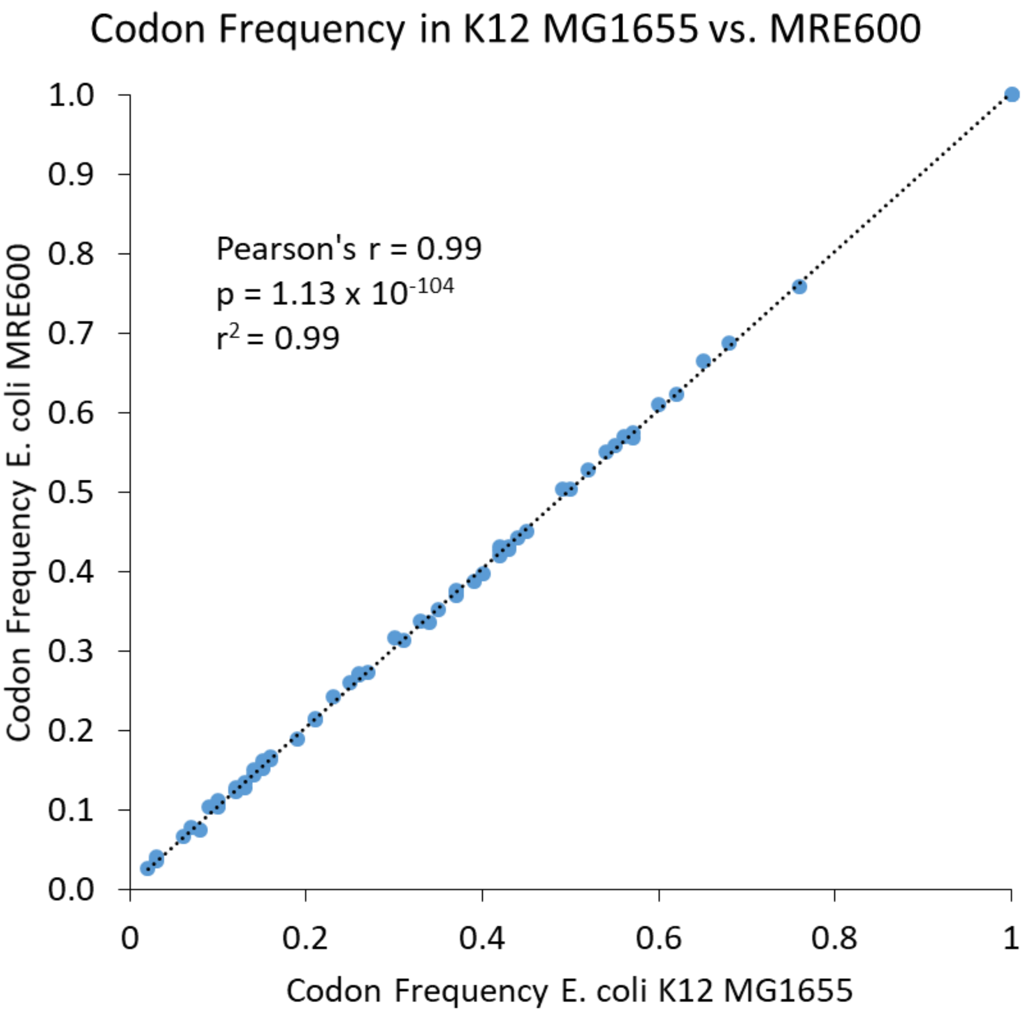


**Fig. S2.**

**Codon usage data related to Figure 1 and the TxTL assay**. Codon usage bias (CUB) reported here as frequency for all 64 codons in *E. coli* MRE600 vs. K12 MG1655 to analyze similarity. Codon use does not significantly differ between the two strains, indicating the tRNA profile in MRE600 is likely similar based on the same observed translational selection. Each point represents the frequency of an individual codon ($n$ = 64) in either of the 2 strains. A full set of annotated protein coding sequences for MRE600 and MG1655 were downloaded from NCBI for this analysis (CP014197.1 and NC_000913.3, respectively). Pearson's r, and linear regression $r^2$ values are calculated from 64 individual codon frequency data points.

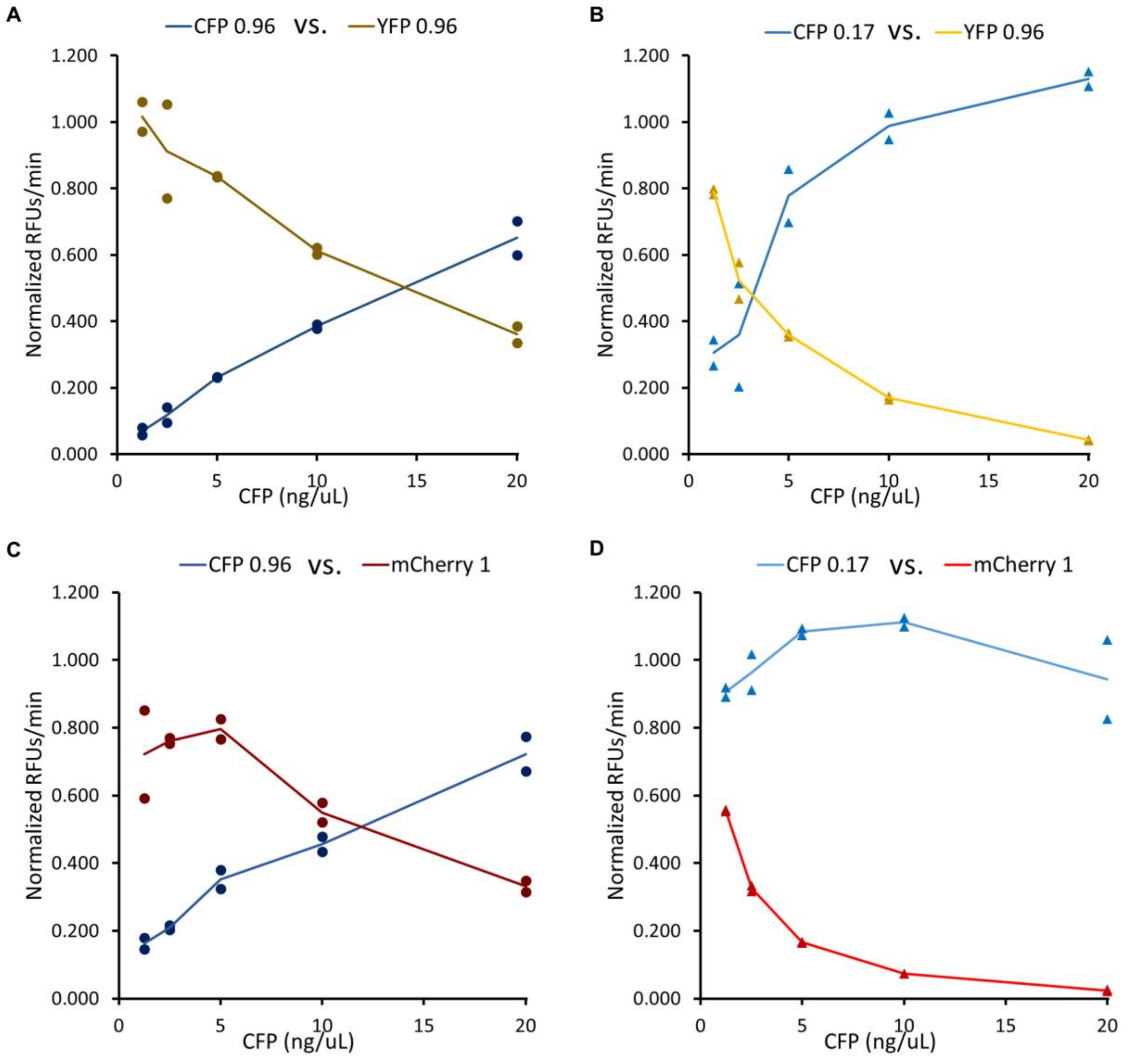


**Fig. S3.**

**Supplementary in vitro reaction rate data related to figure 1.** Relative protein synthesis rates for CFP/YFP pairs (**A** and **B**) and CFP/mCherry pairs (**C** and **D**) normalized to each gene expressed in isolation. Data represent CFP with either high CAI (0.96) or low CAI (0.17) titrated against YFP or mCherry with high CAI (0.96 or 1.00 respectively). Sequences with high CAI exhibit relatively linear tradeoffs in protein expression rate, while titration of a low CAI sequence causes non-linear reduction in YFP or mCherry protein expression. $n$ = 2 TxTL reaction replicates of single re-codes for each concentration level shown on the graphs. Solid lines are connected to sample means.

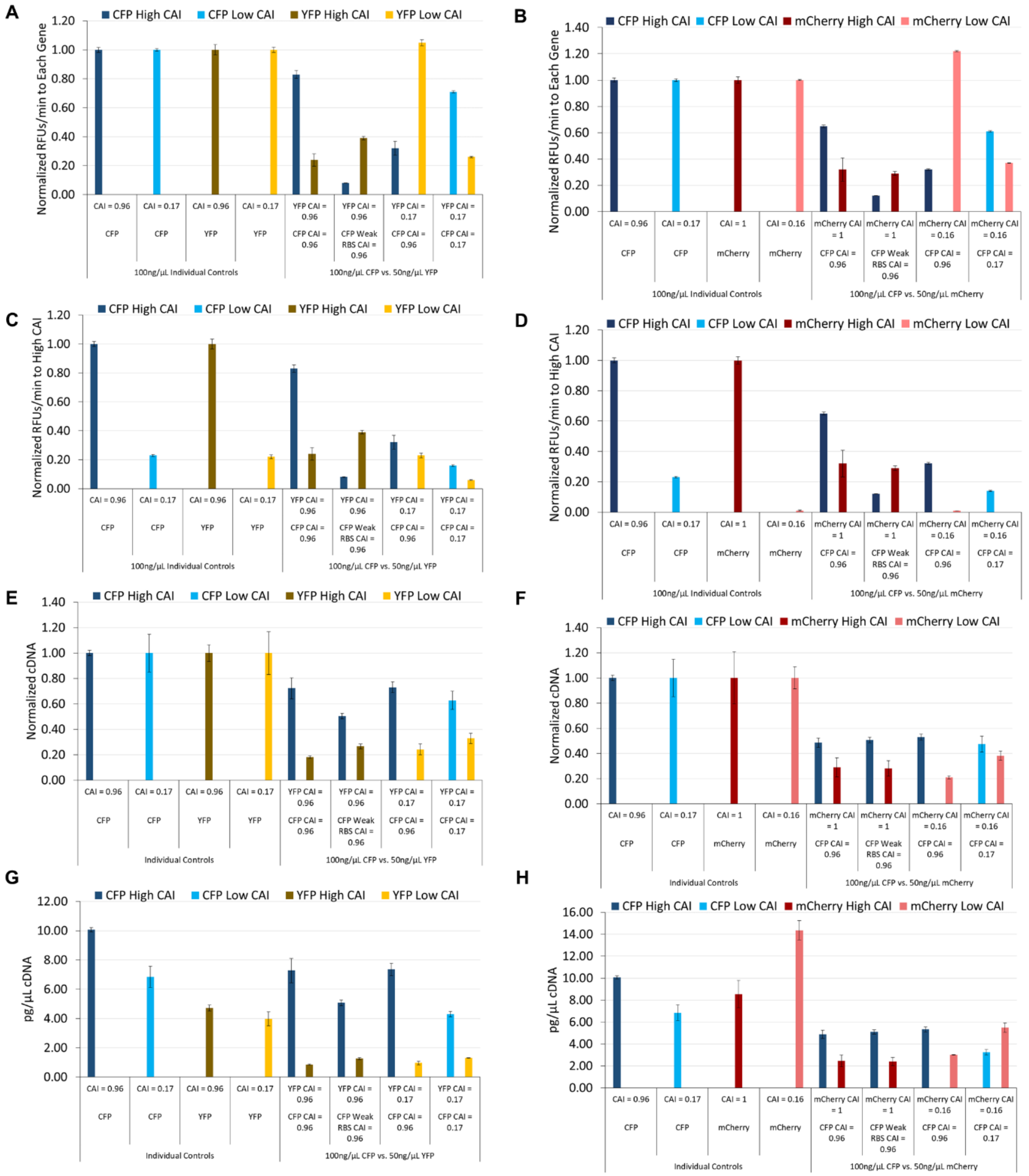

**Fig. S4.**

**Supplementary reaction rate and cDNA data related to figure 1.** (**A** and **B**) Relative protein synthesis rates for CFP/YFP pairs (**A**) and CFP/mCherry pairs (**B**) normalized to each gene expressed in isolation. Protein synthesis in rare codon rich YFP or mCherry sequences (low CAI) is not affected by competition with high CAI CFP sequences. (**C** and **D**) Relative protein synthesis rates for the same CFP/YFP pairs normalized instead to the high CAI sequences for each protein. Low CAI sequences consistently express at a slower rate. (**E** and **F**) Normalized cDNA levels for each reaction measured after 2.5 h. cDNA levels for competition reactions were measured and normalized to cDNA for individual genes expressed in isolation. (**G** and **H**) Absolute concentration of measured cDNA in the same reactions for CFP/YFP pairs

(**G**) and CFP/mCherry pairs (**H**) shown in the other panels. While different sequences exhibit moderate fluctuations in cDNA, rare codon rich sequences do not reduce mRNA transcription in the same way they consistently reduce protein translation. A condition with CFP expressed from a weak RBS is included here and indicates there is competition that occurs at the level of transcription to a certain extent, but competition at the level of translation is apparent between re-coded sequences that vary in relative protein expression, but have similar cDNA levels. $n$ = 3 reaction replicates for each data point, all bars represent means ± SD.

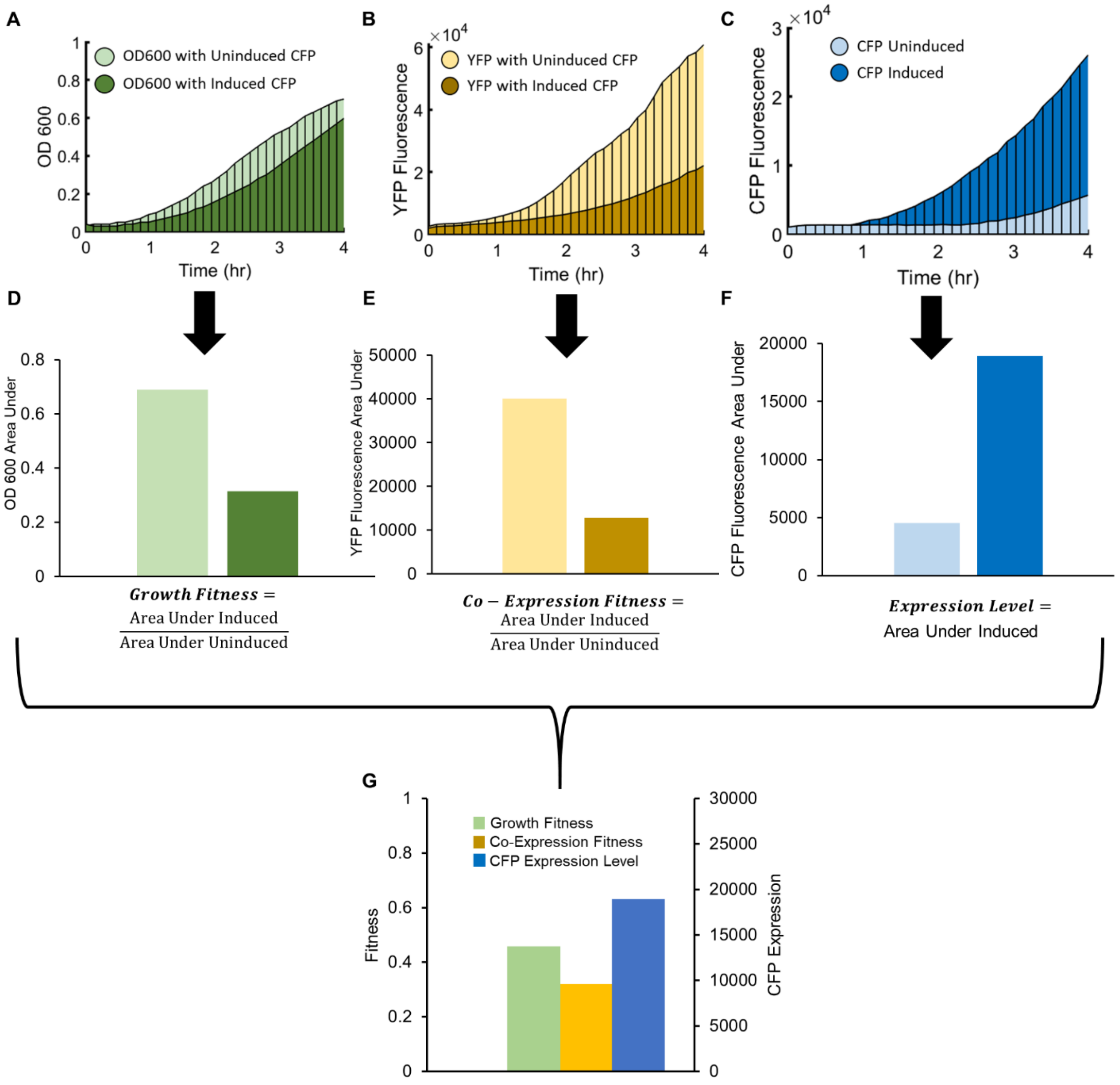


**Fig. S5.**

**Data processing workflow for growth and fluorescent data related to Figure 2.** (**A**–**C**) Signals are gathered by taking the area under the curve (AUC) of growth (**A**) quantified by OD600, or fluorescence YFP (**B**) or CFP (**C**), using a numerical trapezoidal integrator in MATLAB. The timespan is generally defined by the maximum amount of time before any of the signals saturate in any of the samples being measured within an experiment. AUC values are represented here by bar plots (**D**–**F**). These are further processed (**G**) to calculate fitness values (i.e., AUC induced ÷ AUC uninduced for OD600 or YFP signal yielding Growth Fitness or Co-Expression Fitness respectively). The absolute AUC for CFP of the induced culture is used to calculate Expression Level. Growth defects as well as a reduction in YFP expression are generally observed when CFP is induced.

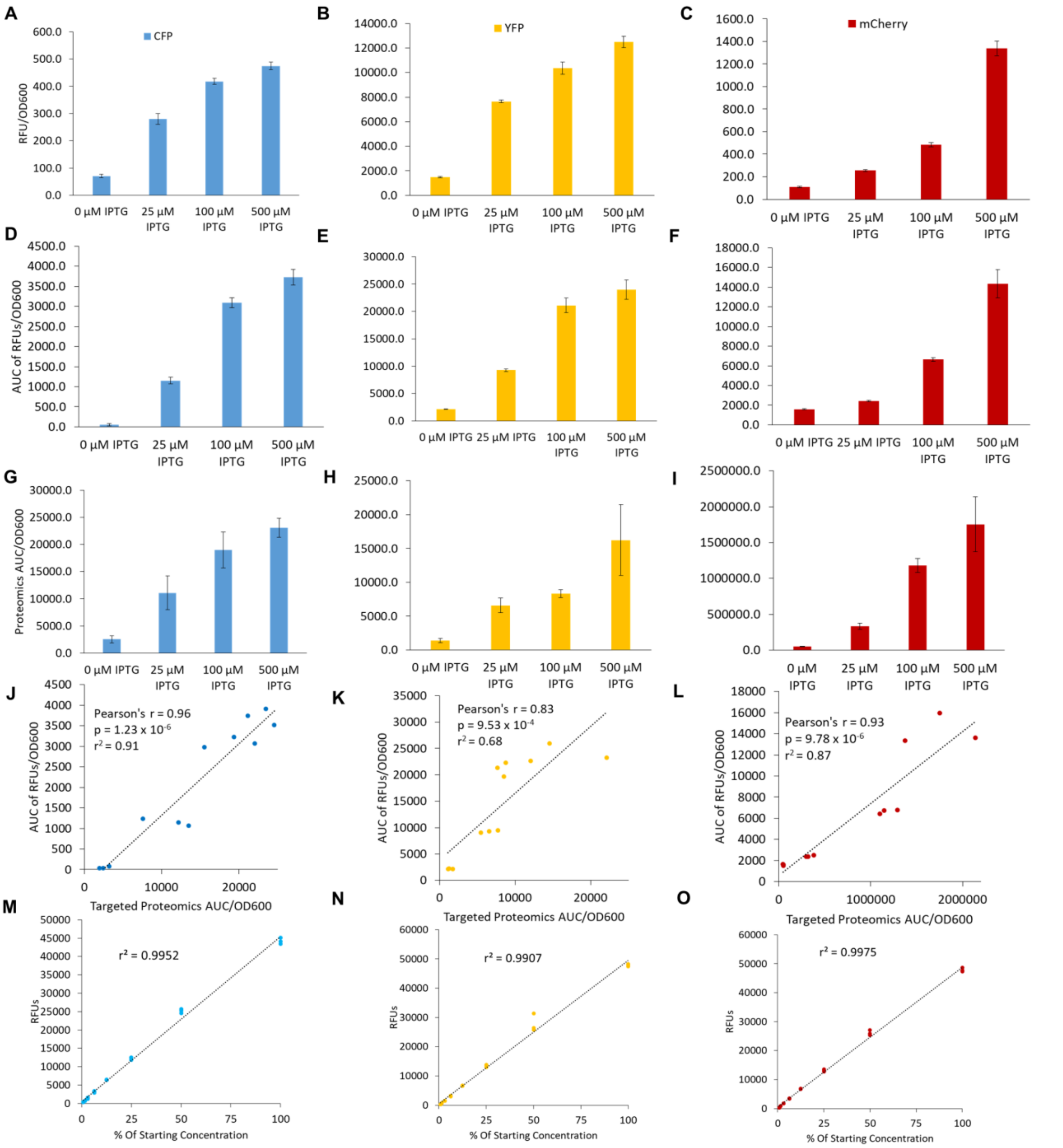


**Fig. S6.**

**Assay validation data related to Figure 2.** All data are for the high CAI sequences, and are OD normalized. Final fluorescence (**A – C**), area under the curve of fluorescence data (**D – F**), and protein level (**G – I**) correlate well when validated using LC-MS/MS, (**J – I**) verify the relationship between area under the curve representing total protein amount measured using targeted proteomics, and the area under the curve of RFU data taken from a plate reader. This supports our assumption that fluorescence data are representative of total protein level. Taking fluorescent measurements of diluted endpoint samples indicate that fluorescent protein measurements are linear relative to protein level over a broad range (**M – O**). *n* = 3 reaction replicates for each data point, all bars represent means ± SD, correlation plots represent individual data points.

A

| AA | Codon | Percent Use in Genome | tRNA Recognition Genes (Red on pBAC-RARE2) |
|---|---|---|---|
| Arg | AGG | 3.13% | argW (CCU), argU (UCU) |
| Arg | CGC | 40.63% | argY (ACG), argQ (ACG), argV (ACG), argZ (ACG) |
| Arg | CGT | 38.54% | argY (ACG), argQ (ACG), argV (ACG), argZ (ACG) |
| Ile | ATA | 7.07% | ileY (CAU), ileX (CAU) |
| Ile | ATC | 42.42% | ileV (GAU), ileT (GAU), ileU (GAU) |
| Ile | ATT | 50.51% | ileV (GAU), ileT (GAU), ileU (GAU) |

B

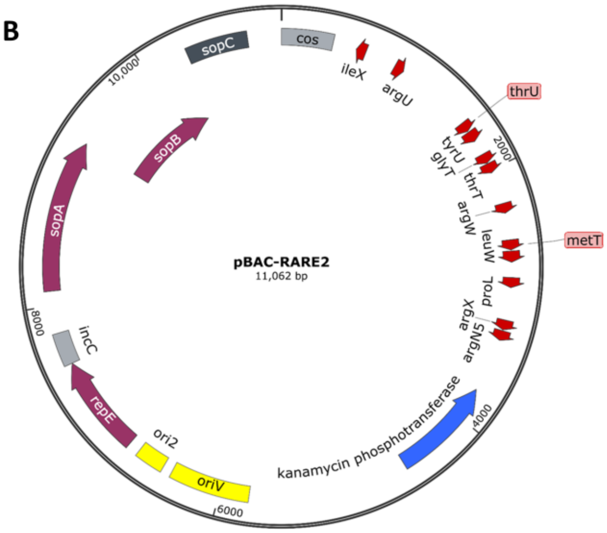


**Fig. S7.**

**Data related to Figure 2.** (**A**) Table of codons incorporated into rare codon re-coded CFP sequences represented in Fig. 1H. For each codon, the percent use in the *E. coli* K12 MG1655 genome is shown, as well as known tRNA genes that can decode each respective codon. tRNA genes that are present on the complemented pBAC-RARE2 are highlighted in red. (**B**) A vector map of pBAC-RARE2 to over-express 12 rare tRNA genes. Individual tRNA genes present on the vector are shown in red. A complete sequence for pBAC-RARE2 can be found in **Data S10**.

A

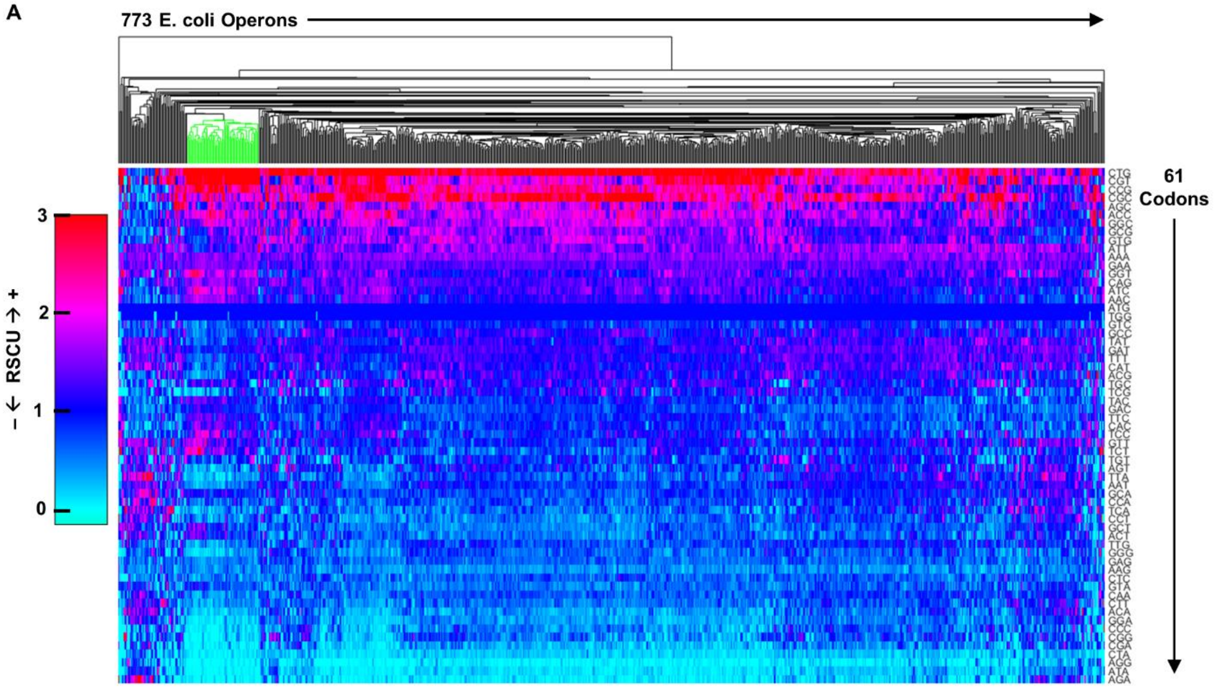


B

| Operon Names (Green Group) | |
|---|---|
| manXYZ | sdaCB |
| ptsHI-crr | rpsF-priB-rpsR-rplI |
| rplNXE-rpsNH-rplFR-rpsE-rpmD-rplO-secY-rpmJ | rpsP-rimM-trmD-rplS |
| rpsMKD-rpoA-rplQ | sspAB |
| cadBA | ybaB-recR |
| rpsJ-rplCDWB-rpsS-rplV-rpsC-rplP-rpmC-rpsQ | recAX |
| ackA-pta | seqA-pgm |
| rplKAJL-rpoBC | rlmE-ftsH |
| rpsLG-fusA-tufA | secMA-mutT |
| focA-pflB | upp-uraA |
| pyrG-eno | hcp-hcr |
| aspA-dcuA | feoABC |
| epd-pgk-fbaA | artPIQM |
| gapA-yeaD | dapA-bamC |
| cmk-rpsA-ihfB | metNIQ |
| accBC | brnQ-proY |
| frdABCD | speAB |
| holC-valS | kbl-tdh |
| glyQS | clpX-lon |
| uxuAB | glmUS |
| hslVU | edd-eda |
| iscRSUA | mreBCD |
| deoCABD | bamB-der |
| guaBA | acrAB |
| cydAB | tolB-pal-cpoB |
| atpIBEFHAGDC | secE-nusG |
| pdhR-aceEF-lpd | ibaG-murA |
| ahpCF | frwCB |

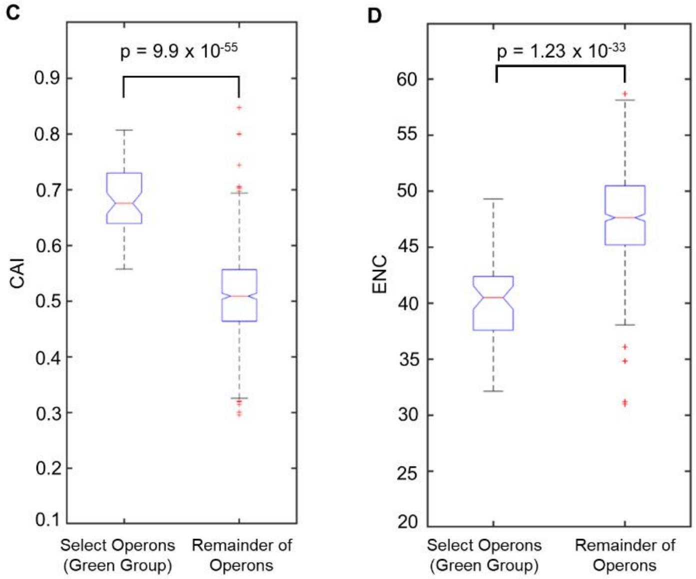


**Fig. S8.**

**Data related to Figure 3.** (**A**) Hierarchal clustering and analysis of RSCU values in 773 *E. coli* operons. Data were generated by arranging RSCU values for 773 *E. coli* operons with more than 1 protein coding sequence into a matrix of 61 rows (one for each codon) and 773 columns (one for each operon). Hierarchal clustering was done in Matlab®, and a unique group of operons with distinct CUB is highlighted (colored green). Note that RSCU values greater than 3 are displayed as the same color as RSCU = 3. (**B**) List of operons/gene names present in the highlighted group. Many genes with distinct CUB are ribosomal proteins or involved in central carbon metabolism. (**C** and **D**). Box and whisker plots of select operons from the highlighted group in green ($n$ = 56 operons) vs. the remainder ($n$ = 717 operons) for CAI values (**C**) or ENC values (**D**). An outlier is a value that is more than 1.5 times the interquartile range away from the bottom or top of the box. A lack of overlapping notches indicates >95% confidence in differences between medians. We observe a significantly higher CAI (median of 0.68 vs. 0.51) and lower ENC (median of 40.5 vs. 47.6) in select operons (green group) relative to the remainder, suggesting that this group has higher CUB overall, and generally uses distinct codons from the rest of the operons in *E. coli*. All p values represent two tailed t tests.

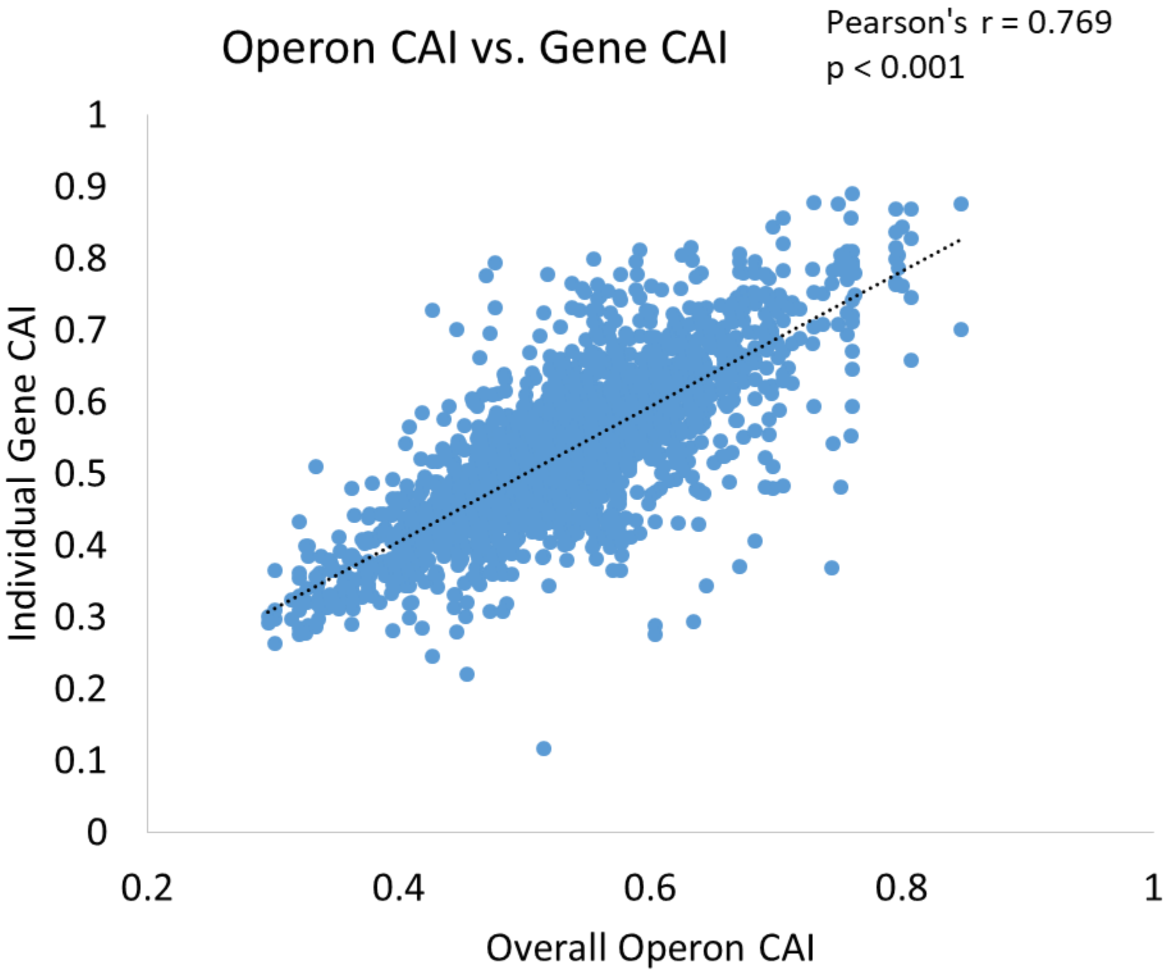


**Fig. S9.**

**Data related to Figure 3.** Analysis of CUB in *E. coli* operons vs. individual gens within those operons shows that genes in the same operon tend to have similar CUB. Data above are from 773 annotated *E. coli* operons with at least 2 protein coding genes (total of 2466 genes). CAI refers to codon adaptation index using highly expressed *E. coli* genes as a reference set (see methods). Pearson's r and p value were calculated from 2466 data points.

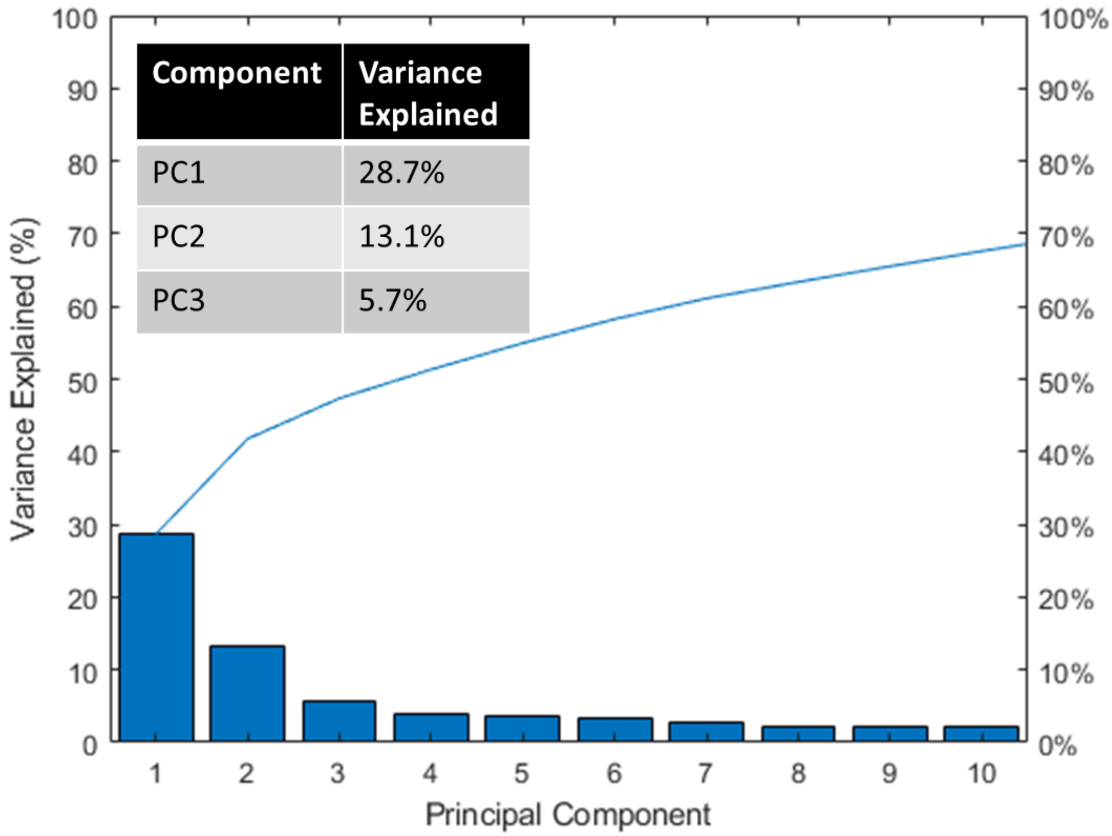


**Fig. S10.**

**Data related to Figure 3 PCA analysis.** Pareto plot showing variance explained by principal component analysis of 773 *E. coli* operons examined in Figure 3. PC1 with 28.7% of total variance was largely found to explain CAI (where CAI refers to codon adaptation index using highly expressed *E. coli* genes as a reference set).

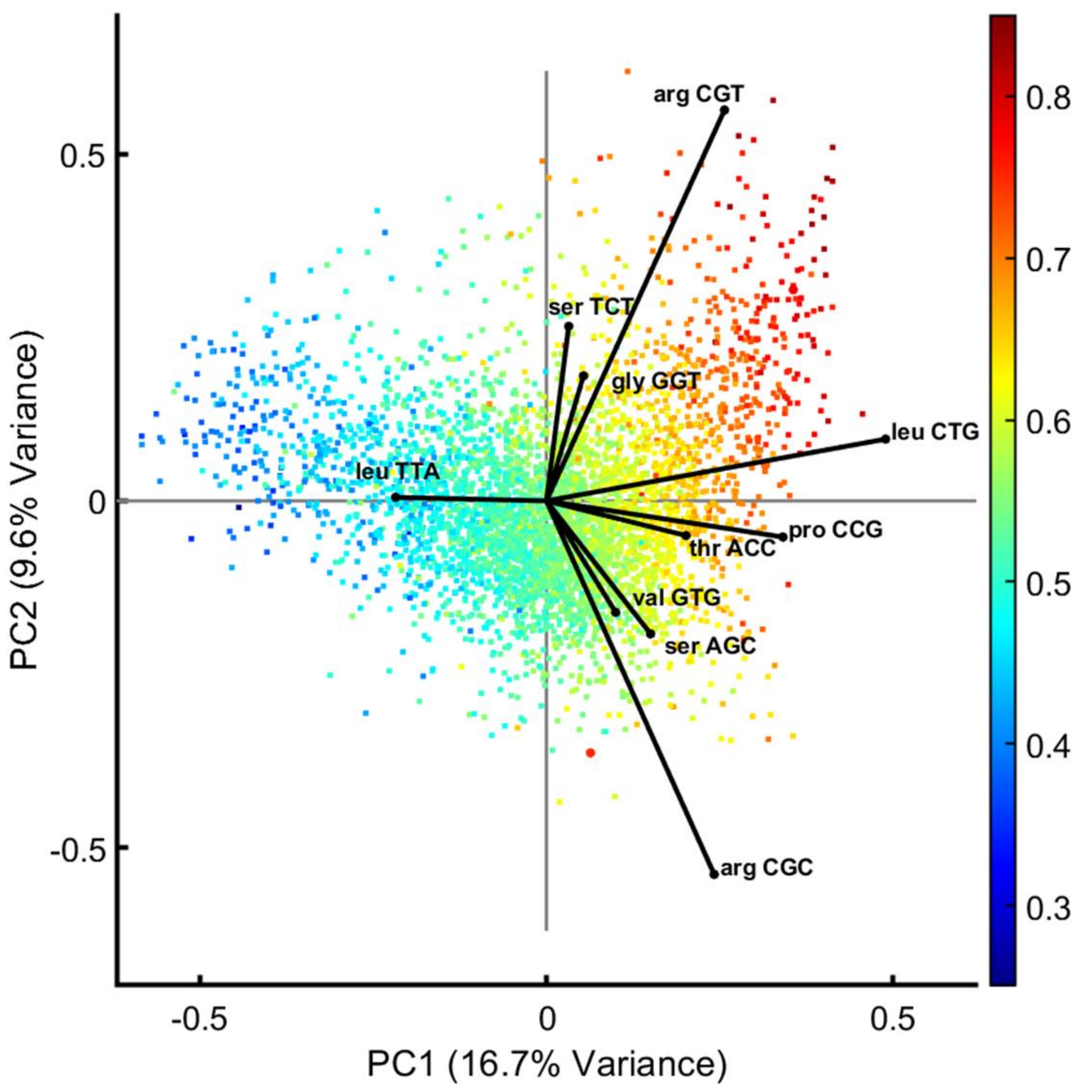


**Fig. S11.**

**Data related to Figure 3 PCA analysis.** PCA analysis of RSCU in 4,311 E. coli genes (as opposed to operons) with loadings mapped for the 10 codons with the highest contribution to variance. CAI is mapped onto individual genes and indicated in the figure legend. PCA on individual genes as opposed to operons reveals similar CUB trends across the E. coli transcriptome.

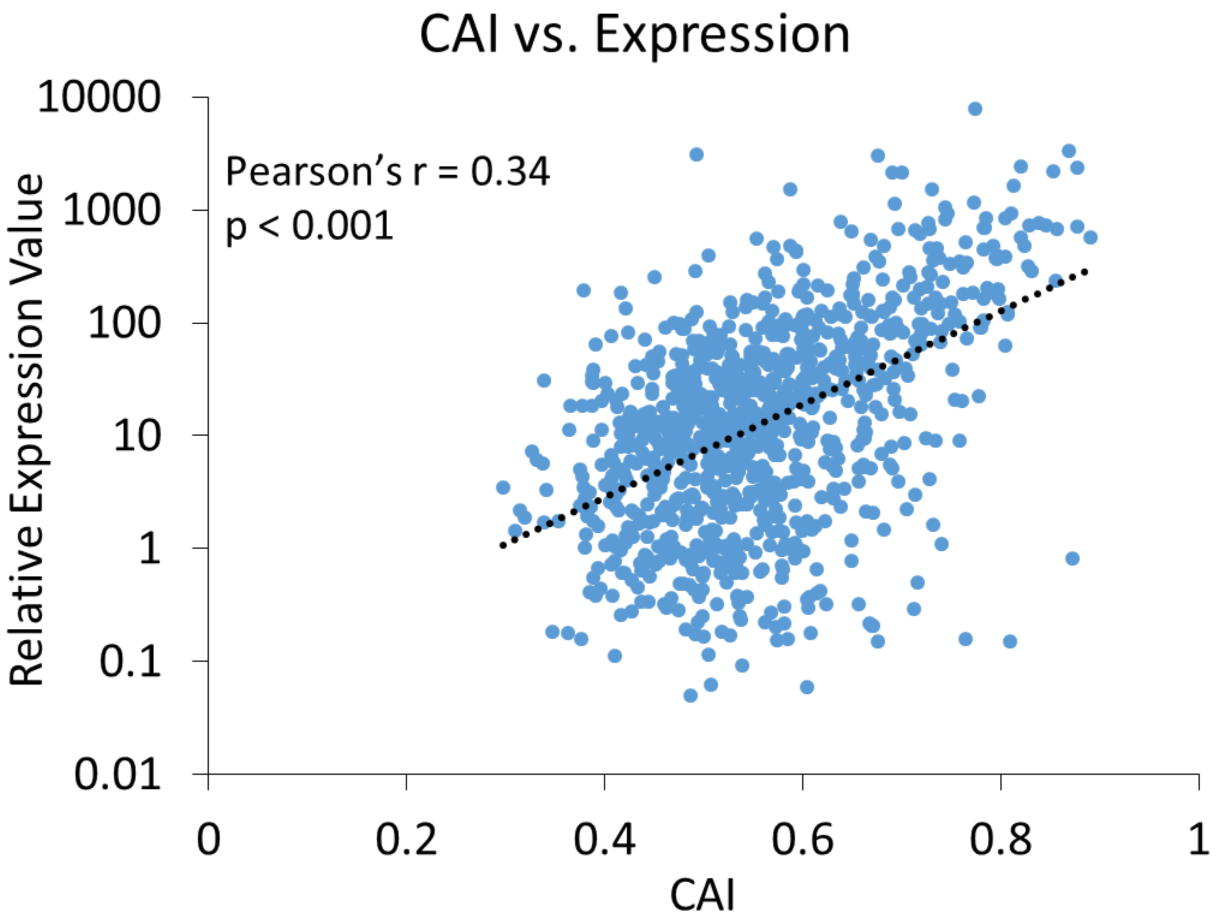


**Fig. S12.**

**CAI vs. expression level correlation data related to figure 3.** Expression data are from Taniguchi et al. (*48*) Here we correlate calculated CAI value (where CAI refers to codon adaptation index using highly expressed *E. coli* genes as a reference set) with published expression data for genes where expression level was reported (total of $n$ = 1014 genes represented here). Note that Pearson's r is calculated based on raw protein level (as opposed to log values).

Anticodons Available in E. coli K12 MG1655 (shaded cells)

Canonical Wobble — Inosine Wobble — Lysidine Wobble

| AA | Codon | Anti | Copy # | AA | Codon | Anti | Copy # | AA | Codon | Anti | Copy # | AA | Codon | Anti | Copy # |
|---|---|---|---|---|---|---|---|---|---|---|---|---|---|---|---|
| F | TTT | AAA | | L | CTT | AAG | | I | ATT | AAT | | V | GTT | AAC | |
| F | TTC | GAA | 2 | L | CTC | GAG | 1 | I | ATC | GAT | 3 | V | GTC | GAC | 2 |
| L | TTA | TAA | 1 | L | CTA | TAG | 1 | I | ATA | TAT | | V | GTA | TAC | 5 |
| L | TTG | CAA | 1 | L | CTG | CAG | 4 | M | ATG | CAT | 2* | V | GTG | CAC | |
| S | TCT | AGA | | P | CCT | AGG | | T | ACT | AGT | | A | GCT | AGC | |
| S | TCC | GGA | 2 | P | CCC | GGG | 1 | T | ACC | GGT | 2 | A | GCC | GGC | 2 |
| S | TCA | TGA | 1 | P | CCA | TGG | 1 | T | ACA | TGT | 1 | A | GCA | TGC | 3 |
| S | TCG | CGA | 1 | P | CCG | CGG | 1 | T | ACG | CGT | 2 | A | GCG | CGC | |
| Y | TAT | ATA | | H | CAT | ATG | | N | AAT | ATT | | D | GAT | ATC | |
| Y | TAC | GTA | 3 | H | CAC | GTG | 1 | N | AAC | GTT | 4 | D | GAC | GTC | 3 |
| Stop | TAA | TTA | | Q | CAA | TTG | 2 | K | AAA | TTT | 6 | E | GAA | TTC | 4 |
| Stop | TAG | CTA | | Q | CAG | CTG | 2 | K | AAG | CTT | | E | GAG | CTC | |
| C | TGT | ACA | | R | CGT | ACG | 4 | S | AGT | ACT | | G | GGT | ACC | |
| C | TGC | GCA | 1 | R | CGC | GCG | | S | AGC | GCT | 1 | G | GGC | GCC | 4 |
| Stop | TGA | TCA | | R | CGA | TCG | | R | AGA | TCT | 1 | G | GGA | TCC | 1 |
| W | TGG | CCA | 1 | R | CGG | CCG | 1 | R | AGG | CCT | 1 | G | GGG | CCC | 1 |

**Fig. S13.**

**Available tRNA/anti-codons in *E. coli* K12 MG1655 and corresponding copy # for each.** tRNA copy # data taken from the genomic tRNA database (http://gtrnadb.ucsc.edu/). Codon-anticodon interactions and figure adapted from Reis et al. (*7*) Special cases where modified tRNAs are required for codon recognition are highlighted for inosine and lysidine modifications. Generally speaking, amino acids with more than 2 available codons also have more than a single tRNA available, while most amino acids with only 2 codons share a single tRNA anticodon. In the special case of the ATA codon for isoleucine, the anticodon is identical to the methionine tRNA, and the copy # is shown only for the isoleucine CAT tRNA (denoted with *). Note that T is used here in place of U.

| Amino Acid | mCherry #/% | CFP #/% |
|---|---|---|
| Ala | 10/4.2% | 8/3.4% |
| Cys | 0/0% | 2/0.8% |
| Asp | 13/5.5% | 18/7.6% |
| Glu | 21/8.8% | 13/5.5% |
| Phe | 9/3.8% | 11/4.6% |
| Gly | 24/10.1% | 20/8.4% |
| His | 6/2.5% | 10/4.2% |
| Ile | 8/3.4% | 10/4.2% |
| Lys | 22/9.2% | 19/8% |
| Leu | 13/5.5% | 18/7.6% |
| Asn | 6/2.5% | 13/5.5% |
| Pro | 12/5% | 9/3.8% |
| Gln | 8/3.4% | 7/2.9% |
| Arg | 8/3.4% | 8/3.4% |
| Ser | 11/5% | 9/3.8% |
| Thr | 12/5.5% | 17/7.1% |
| Val | 14/6.4% | 15/6.3% |
| Tyr | 12/5.5% | 8/3.4% |

**Fig. S14.**

**Amino acids re-coded for CFP and mCherry related to Figure 4.** Differences in the number of amino acids between CFP and mCherry could contribute to noise in the codon sensitivity analysis dataset. Notably cysteine is underrepresented in CFP and absent in mCherry. The # refer to the total number re-coded, while the % refer to the number recoded as a percentage of all amino acids in the protein.

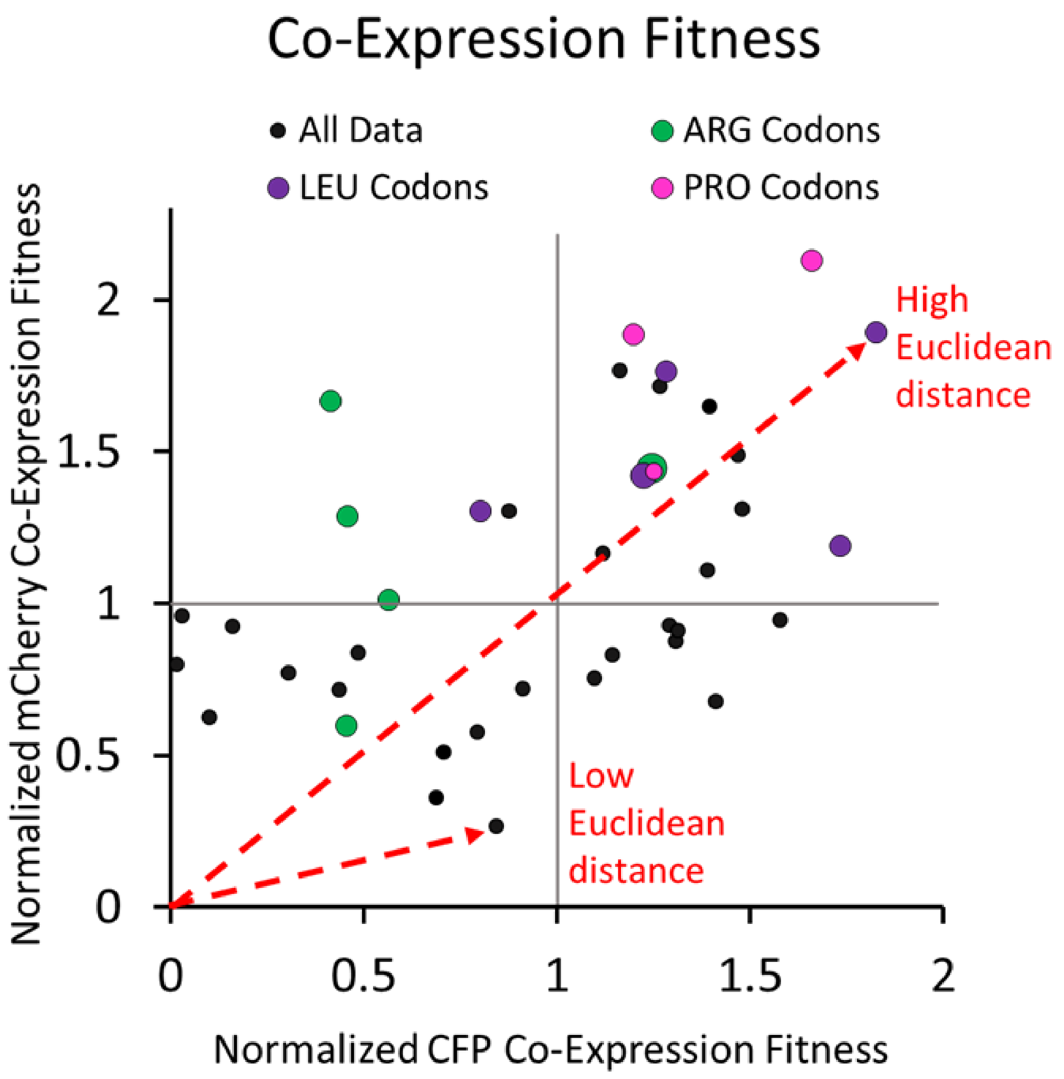


**Fig. S15.**

**Method for deriving χ.** Each data point representing normalized Co-Expression Fitness (as shown in Fig. 4B) for every codon re-coded in CFP and mCherry ($n$ = 40) was quantified by taking the Euclidean distance from the origin to the codon coordinates. These raw scores were then use to create a new table of weights for calculating χ. Raw scores and calculated weights are given Data S8.

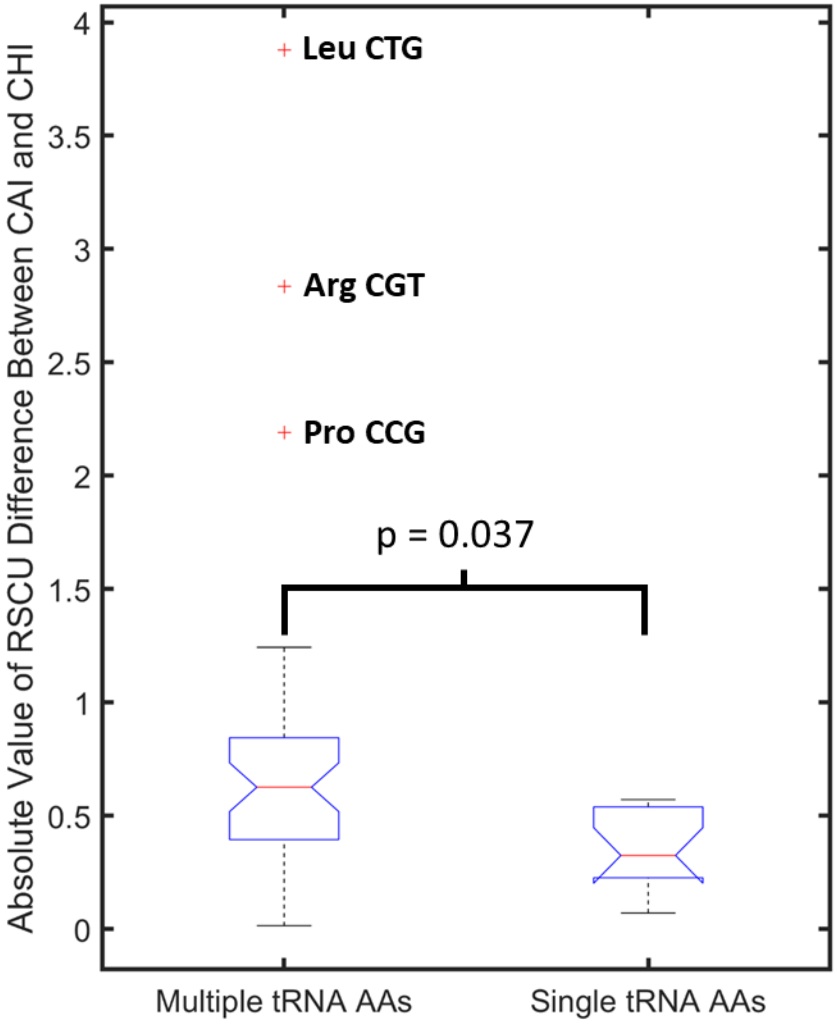


**Fig. S16.**

**Data related to figure 5a.** RSCU differences between individual codons for CAI and $\chi$ indicates significantly larger differences for amino acids with multiple available tRNAs, suggesting tRNA competition could play a role in fitness improvements observed for $\chi$ re-coded genes. In this analysis, each amino acid is grouped based on whether there exists a single tRNA available (16 total codons) or multiple (43 total codons) (see **Fig. S9**, there are 10 AAs with multiple tRNA anticodons in *E. coli*). The RSCU difference between CAI and $\chi$ is calculated based on the difference for each codon between expected RSCU values for a perfectly adapted sequence on either scale (i.e., the difference between the RSCU value on either scale represented in **Fig. 5A**). An outlier is a value that is more than 1.5 times the interquartile range away from the bottom or top of the box.  The observed lack of overlapping notches indicates >95% confidence in differences between medians. Statistical test is a two tailed t test.

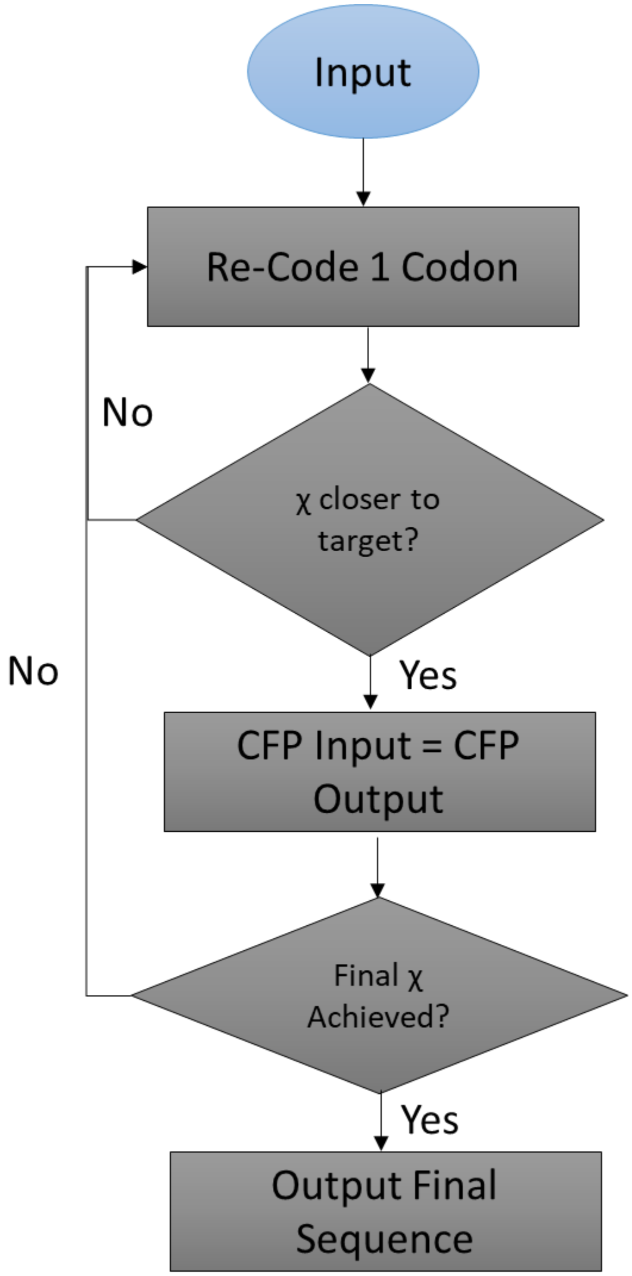


**Fig. S17.**

**Algorithm used to create χ re-codes in Figure 5.** A greedy algorithm implemented in Matlab® was used to create χ re-coded CFP and mCherry sequences. As pictured, the algorithm starts with any sequence (in this case we used a randomly re-coded version of CFP or mCherry as the input to avoid any initial bias), and randomly mutates a codon to a synonymous alternative. It then evaluates whether the new sequence is closer to the objective χ value. This is repeated until the final χ score is obtained. The same algorithm can be implemented with any objective function, e.g., for CAI, GC content, ENC, etc.

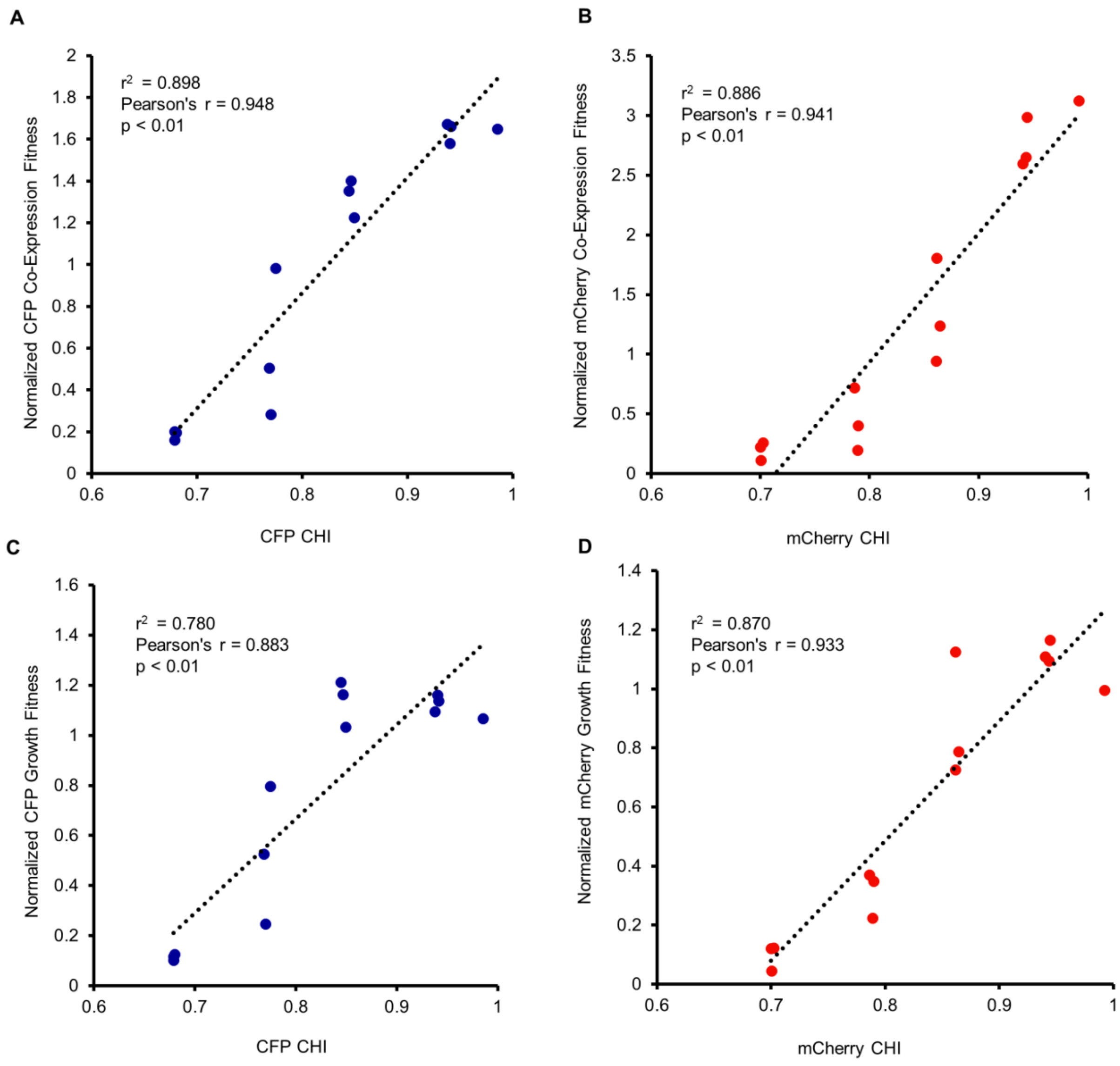


**Fig. S18.**

**Correlation data from Figure 5 C–E demonstrating a strong relationship between Growth and Co-expression Fitness with χ. (A** and **B):** Strong correlation for CFP and mCherry χ re-codes with Co-Expression Fitness (as quantified by the chromosomally integrated YFP reporter in **Fig. 2**). (**C** and **D):** Strong correlation for CFP and mCherry χ re-codes with Growth Fitness (as quantified by $OD_{600}$). In each case, results were normalized relative to the high CAI parent control. Individual data points are shown representing *n* = 13 normalized means for each correlation plot. Pearson's r, and linear regression $r^2$ values are calculated from 13 points for each plot.

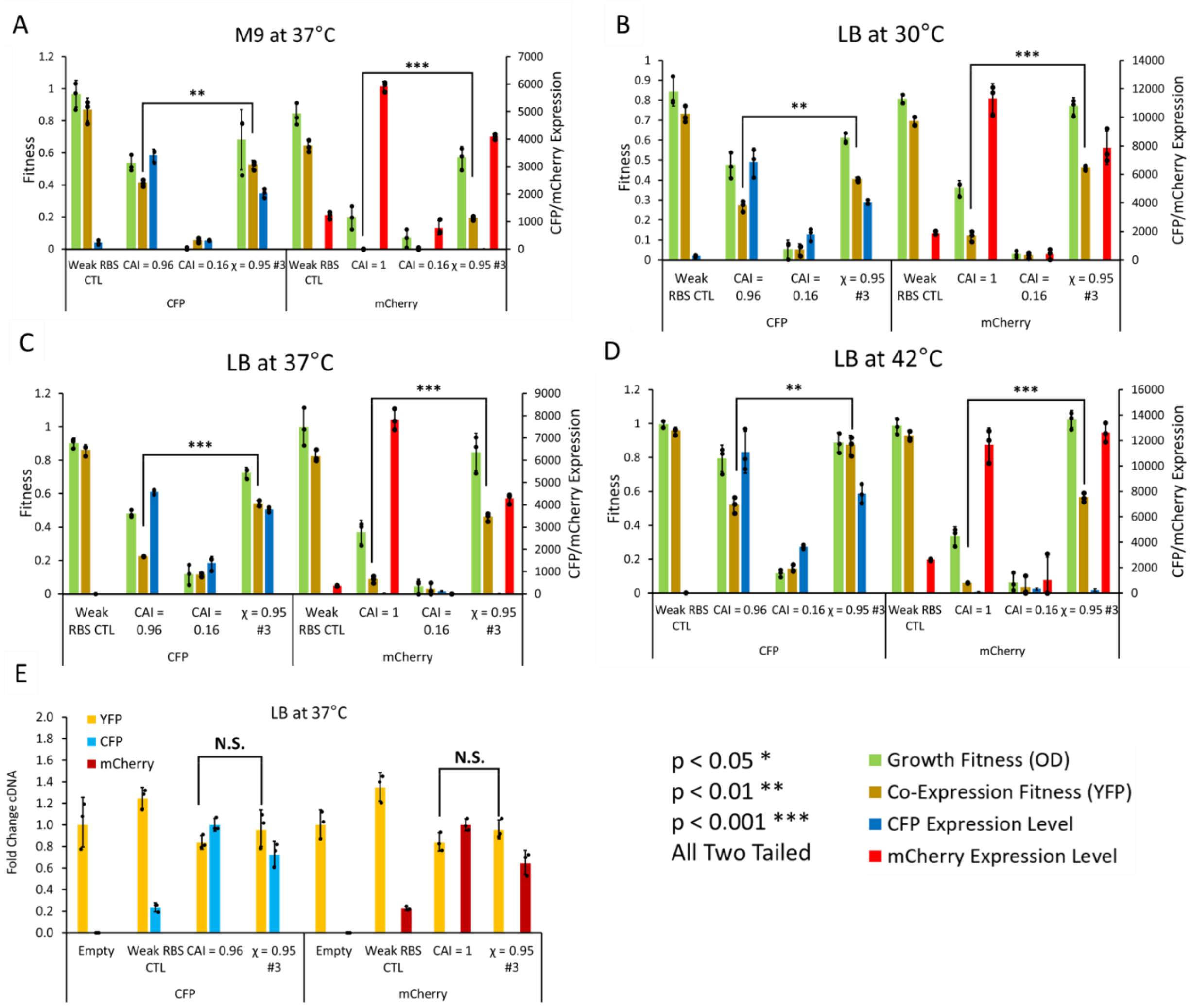


**Fig. S19.**

**Data related to figure 6 testing χ vs. CAI sequences under different conditions.** A weak RBS control was included, along with high vs. low CAI sequences, and the top performing χ = 0.95 #3 sequence using either M9 minimal media at 37°C (**A**), or varying temperatures between 30°C and 42°C in rich LB media (**B–D**). (**E**) qRT-PCR analysis of YFP, CFP, and mCherry fold change in cDNA levels from the 37°C LB condition. Changes in Co-expression fitness are not accounted for by RNA expression differences. Low mCherry and CFP RNA expression level for the weak RBS control is likely accounted for by higher degradation rates for poorly translated mRNA. All statistical tests are two tailed t tests, $n = 3$ biological replicates for each data point, all bars represent means ± SD. N.S. refers to $p > 0.05$. High CAI and high χ CFP and mCherry cDNA levels can be directly compared despite having different amplicons, given their similar amplification efficiency (**Data S11**).

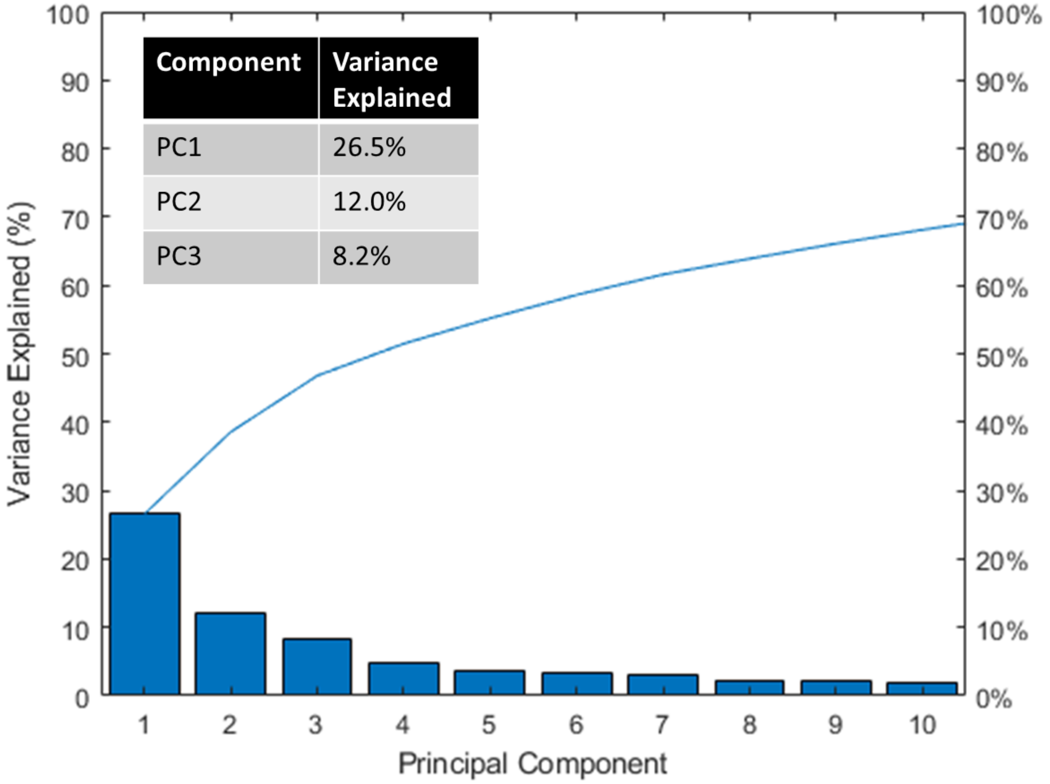


**Fig. S20.**

**Data related to Figure 7 PCA analysis.** Pareto plot showing variance explained by principal component analysis of 773 *E. coli* operons along with the χ re-coded sequences examined in Figure 6. PC1 with 26.5% of total variance was again largely found to explain CAI (where CAI refers to codon adaptation index using highly expressed *E. coli* genes as a reference set), while PC3 with 8.2% of total variance explained the χ sequences very well.

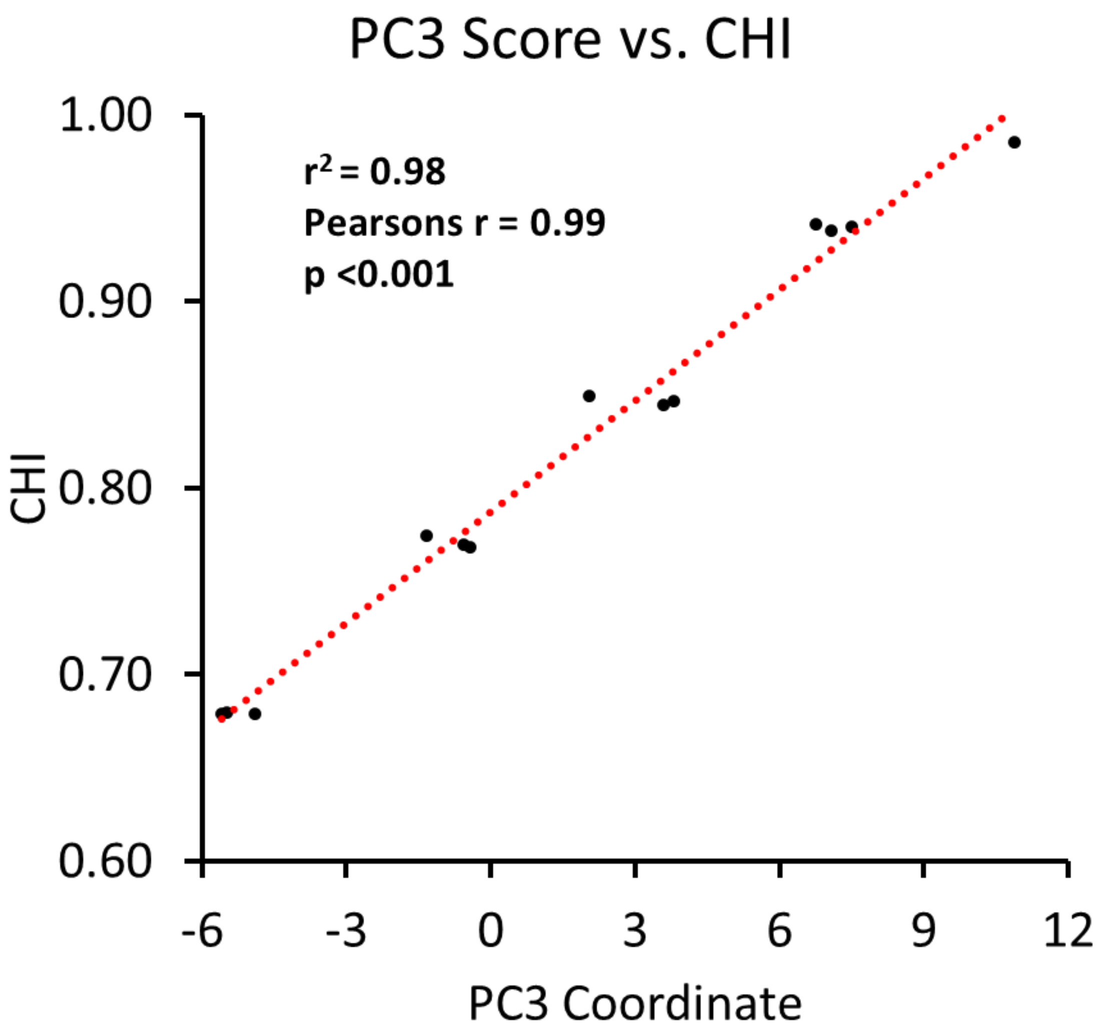


**Fig. S21.**

**Data related to Figure 7 PCA analysis.** Principal component 3 explains CHI ($\chi$) re-coded sequence variation from Figure 7 very well, as there is a very strong correlation. Pearson's r, and linear regression $r^2$ values are calculated from $n = 13$ individual data points representing single re-coded sequences.

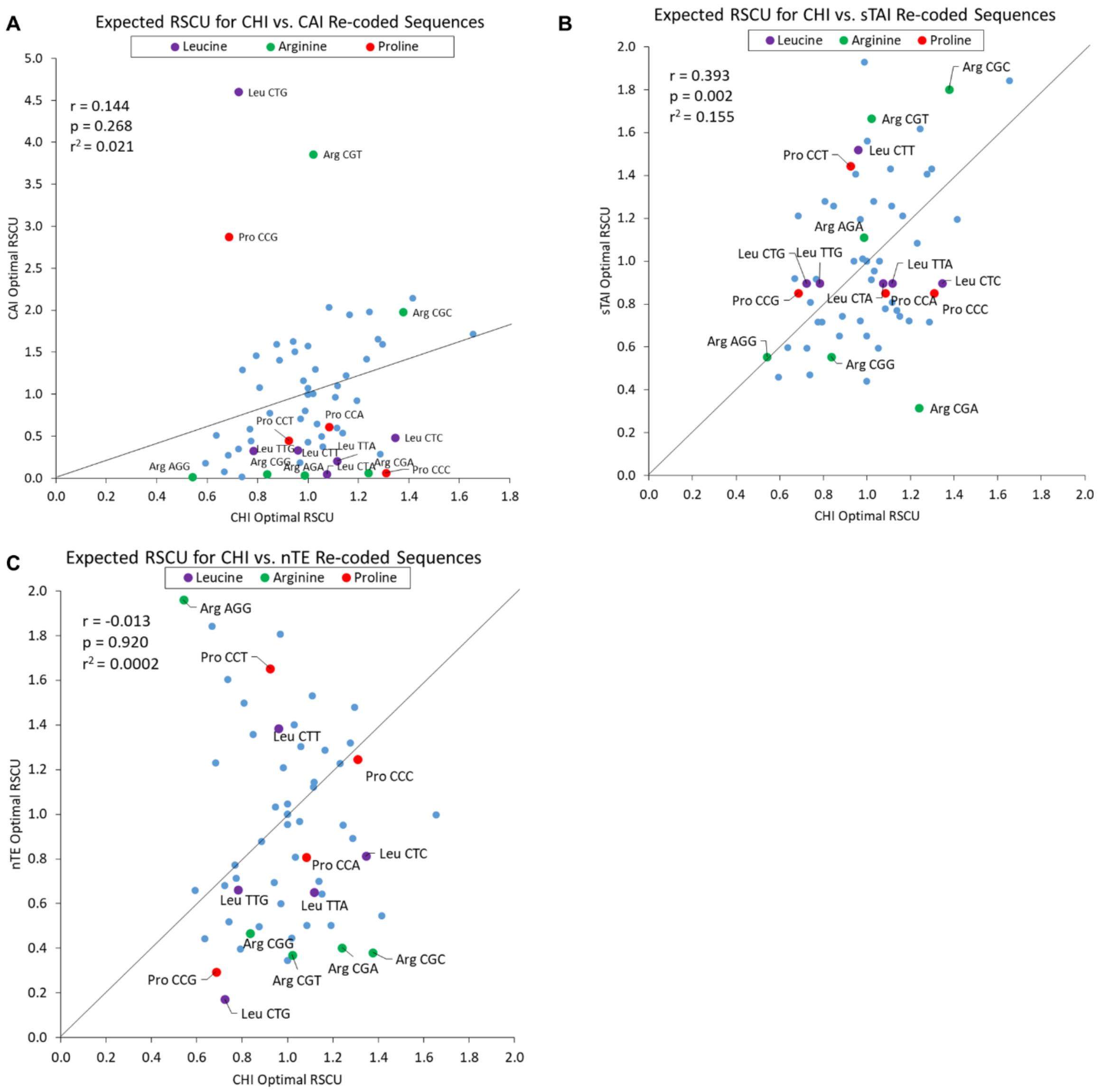


**Fig. S22.**

**Comparison of CHI (χ) with other codon usage bias indices.** Each correlation analysis is done using Pearson's r between χ and another index based on predicted RSCU values for each codon in a perfectly adapted sequence for *n* = 61 codons (excluding stop codons). (**A**) **χ vs. CAI.** The 3 outlier Arg, Leu, and Pro codons are clearly favored with CAI (where CAI is in reference to highly expressed *E. coli* genes). There is general a poor correlation between the two scales (**B**) **χ vs. sTAI.** The correlation between χ and sTAI is better, but there is generally less differentiation between codons for sTAI (e.g., the sTAI derived RSCU for several Leu and Pro codons is the same). (**C**) **χ vs. nTE.** χ and nTE are not well correlated. Despite considering supply vs. demand, nTE appears to exaggerate differences between codons resulting in the over-avoidance of some and over-favoring of others relative to χ. Pearson correlation coefficients and linear regression $r^2$ values were calculated from n = 61 codon RSCU values in each plot. Dotted lines are equivalence lines between each axis.

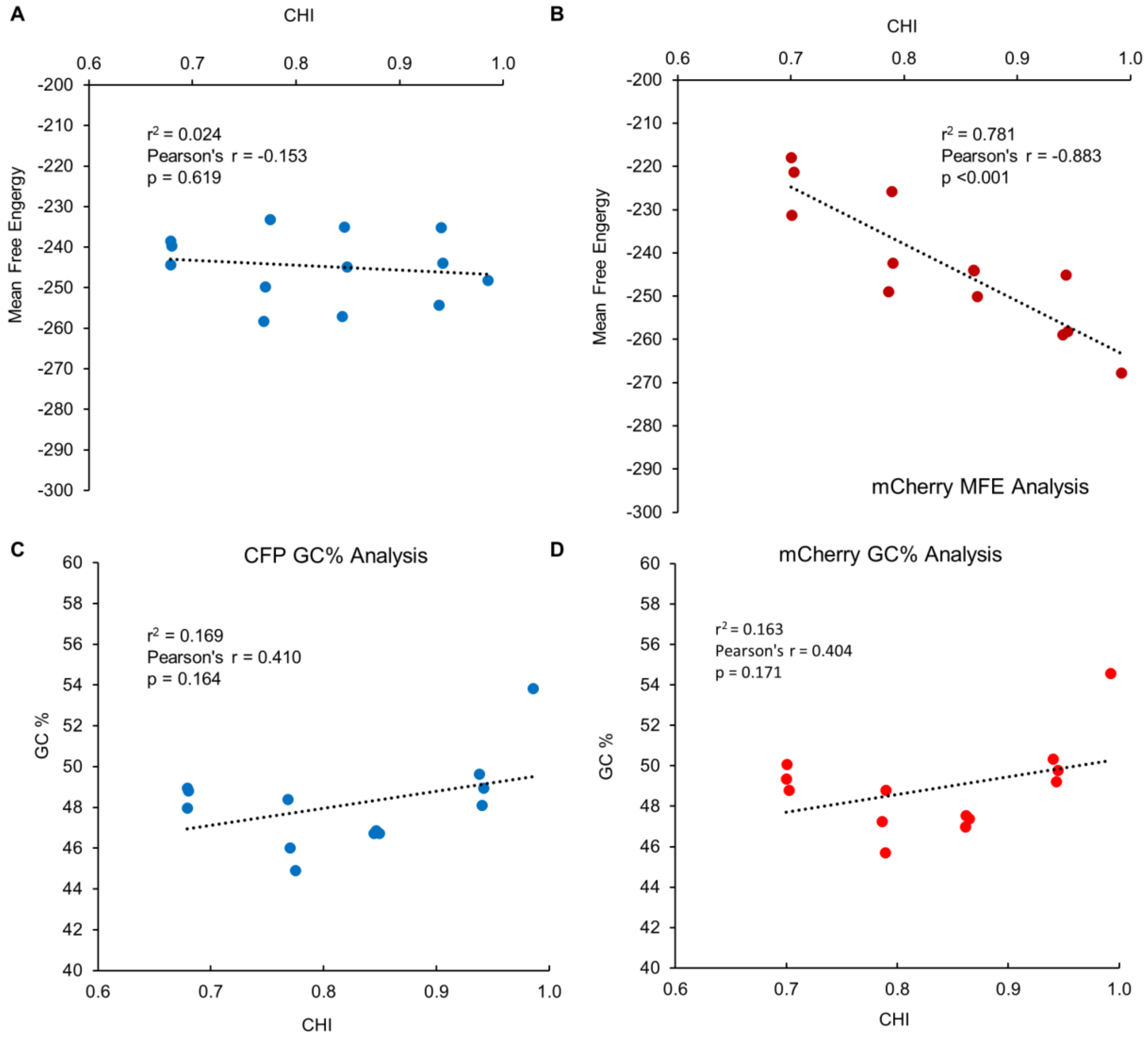


**Fig. S23.**

**Correlations between MFE (mean free energy) or GC content of CHI (χ) re-coded sequences.** (**A** and **B**) Correlation between χ and MFE for CFP and mCherry re-coded sequences. There is no correlation for the CFP re-codes, but there is a negative correlation for mCherry recodes. This implies that for mCherry specifically there is likely additional structure forming due to the incorporation of χ favored codons, but the trend is not generalizable as it is not observed for CFP. (**C** and **D**) Correlation between χ and GC content for CFP and mCherry re-coded sequences. In both cases there is no correlation, and GC content does not substantially vary as a function of χ value. Pearson's r, and linear regression $r^2$ values are calculated from $n$ = 13 individual re-codes for each plot.

**A**

| Vendor/Gene | CHI | CAI_Highly Expressed E. coli Genes | CAI_All E. coli Genes |
|---|---|---|---|
| Genewiz_Gene 1 | 0.844 | 0.789 | 0.995 |
| Genewiz_Gene 2 | 0.843 | 0.792 | 0.991 |
| Genewiz_Gene 3 | 0.814 | 0.804 | 0.987 |
| Genewiz_Gene 4 | 0.808 | 0.751 | 0.970 |
| Genewiz_Gene 5 | 0.819 | 0.766 | 0.973 |
| Genewiz_Gene 6 | 0.804 | 0.747 | 0.968 |
| Genewiz_Gene 7 | 0.817 | 0.751 | 0.976 |
| Genewiz_Gene 8 | 0.843 | 0.779 | 0.978 |
| Genewiz_Gene 9 | 0.819 | 0.773 | 0.984 |
| Genewiz_Gene 10 | 0.810 | 0.750 | 0.968 |
| IDT_Gene 1 | 0.804 | 0.420 | 0.627 |
| IDT_Gene 2 | 0.816 | 0.484 | 0.640 |
| IDT_Gene 3 | 0.816 | 0.426 | 0.627 |
| IDT_Gene 4 | 0.817 | 0.452 | 0.633 |
| IDT_Gene 5 | 0.800 | 0.445 | 0.650 |
| IDT_Gene 6 | 0.801 | 0.440 | 0.641 |
| IDT_Gene 7 | 0.831 | 0.495 | 0.667 |
| IDT_Gene 8 | 0.850 | 0.563 | 0.717 |
| IDT_Gene 9 | 0.809 | 0.476 | 0.641 |
| IDT_Gene 10 | 0.822 | 0.463 | 0.661 |

| Vendor/Gene | CHI | CAI_Highly Expressed E. coli Genes | CAI_All E. coli Genes |
|---|---|---|---|
| Genscript_Gene 1 | 0.811 | 0.635 | 0.754 |
| Genscript_Gene 2 | 0.826 | 0.661 | 0.767 |
| Genscript_Gene 3 | 0.816 | 0.628 | 0.746 |
| Genscript_Gene 4 | 0.809 | 0.628 | 0.743 |
| Genscript_Gene 5 | 0.809 | 0.621 | 0.753 |
| Genscript_Gene 6 | 0.803 | 0.641 | 0.753 |
| Genscript_Gene 7 | 0.828 | 0.642 | 0.752 |
| Genscript_Gene 8 | 0.834 | 0.641 | 0.754 |
| Genscript_Gene 9 | 0.818 | 0.638 | 0.757 |
| Genscript_Gene 10 | 0.821 | 0.631 | 0.747 |

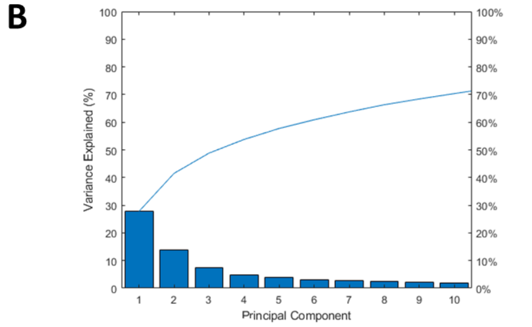


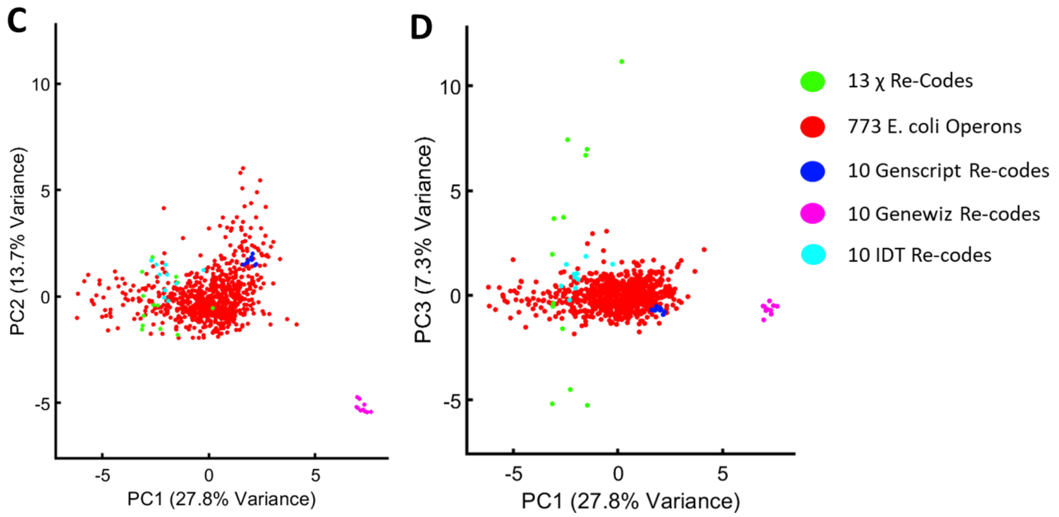


**Fig. S24.**

**CHI (χ) sequences relative to commercial re-coding algorithms.** (**A**) Table of χ and CAI values (shown here using highly expressed *E. coli* genes or the entire genome as a reference set) for 10 genes re-coded using 3 different commercially available free codon optimization tools from IDT, Genewiz, or Genscript. The functional range of χ is between 0.6-1, so none of these sequences appear to explore the χ sequence space, falling at χ ~ 0.8. IDT has the least bias towards either CAI scale. Genewiz appears to bias towards CUB very strongly with respect to all *E. coli* genes, while Genscript appears to adapt the sequence more moderately towards overall host codon usage. (**B**) Pareto plot of PCA of RSCU values of 61 codons for all commercially re-coded genes (30 total) along with 13 χ re-coded sequences (from **Fig. 5**) and 773 *E. coli* operons. 48.8% of total variance is represented by the 1$^{st}$ 3 components. (**C** and **D**). PCA analysis showing PC1 vs. PC2 or PC1 vs. PC3, with categorically labeled points. We know from previous analysis (**Fig. 3**) that PC1 largely represents CAI. It appears that Genscript re-codes most closely follow the natural sequence space of highly expressed genes. IDT re-codes align the closets with χ sequences, but do not explore the χ sequence space in any meaningful way.

**A**

| Phage | NCBI Reference | # of Genes |
|---|---|---|
| Bacteriophage HK022 | AF069308.1 | 69 |
| Bacteriophage lambda | J02459.1 | 66 |
| Bacteriophage M13 | NC_003287.2 | 10 |
| Bacteriophage mu | AF083977.1 | 55 |
| Bacteriophage N4 | NC_008720.1 | 72 |
| Bacteriophage P1 | NC_005856.1 | 110 |
| Bacteriophage P2 | NC_001895.1 | 43 |
| Bacteriophage phi X 174 | NC_001422.1 | 11 |
| Bacteriophage PRD1 | NC_001421.2 | 31 |
| Bacteriophage T3 | NC_003298.1 | 55 |
| Bacteriophage T4 | NC_000866.4 | 278 |
| Bacteriophage T7 | NC_001604.1 | 60 |

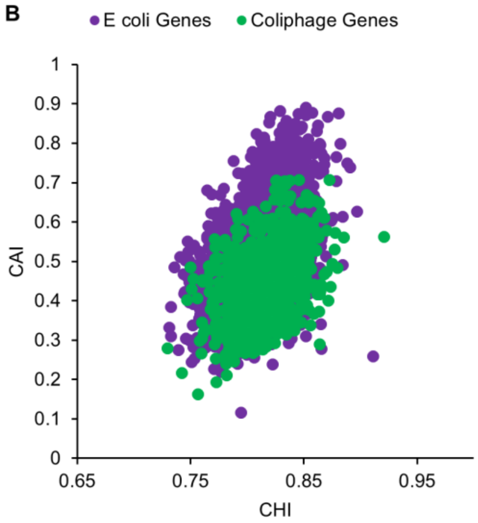


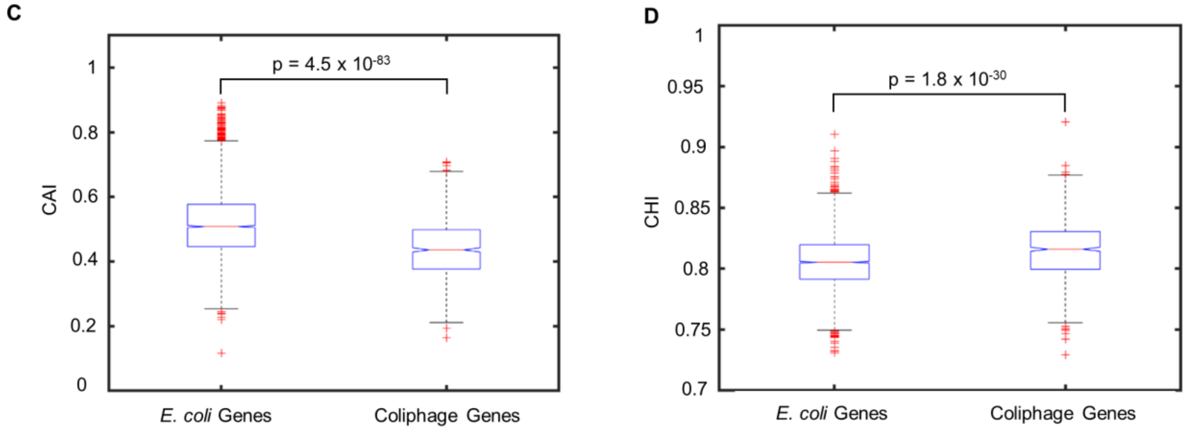


**Fig. S25.**

**CAI and CHI (χ) analysis of phage genes.** (**A**) Coliphages accessed from NCBI used in the analysis. (**B**) Calculated CHI (χ) and CAI (with respect to highly expressed *E. coli* genes) for 4311 individual *E. coli* genes or 860 coliphage genes. Phages generally do not have CAI higher than 0.7 and appear to tend slightly more towards higher χ than *E. coli* genes. (**C** and **D**) Calculated CAI (**C**) and χ (**D**) for 4311 *E. coli* genes and 860 coliphage genes. We observe a slightly higher but significant median CAI for native *E. coli* genes, and a slightly higher but significant median χ for coliphage genes. An outlier is a value that is more than 1.5 times the interquartile range away from the bottom or top of the box.  A lack of overlapping notches indicates >95% confidence in differences between medians. Statistical tests are two tailed t tests.

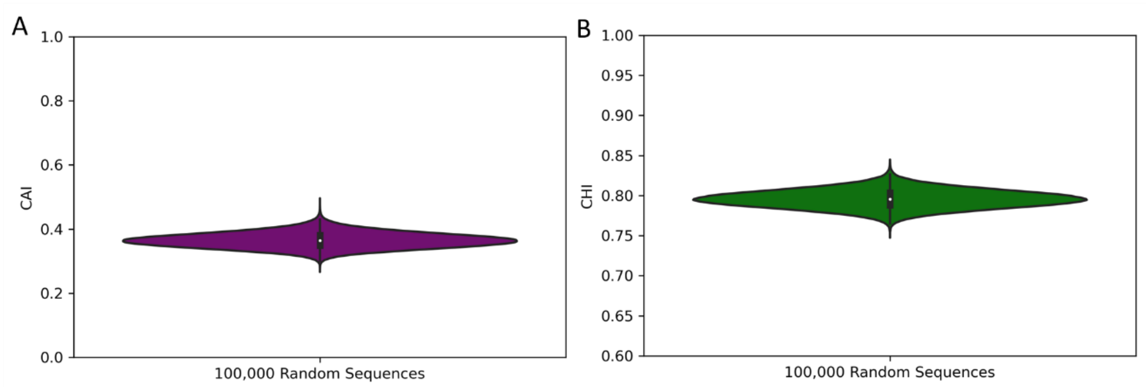


**Fig. S26.**

**CHI (χ) and CAI values of random CFP sequence recodes.** (**A**) Calculated CAI (with respect to highly expressed *E. coli* genes) for 100,000 random sequences. The median CAI is 0.36, and even the highest outlier does not exceed 0.5. This illustrates why randomizing sequences is unlikely to affect global sequence bias towards specific codon use. (**B**) Calculated CHI (χ) for the same random sequences. We see a median χ score of 0.80, and the highest outlier does not exceed 0.85, again illustrating a lack of global variation in specific codon use towards higher χ from randomized sequences.

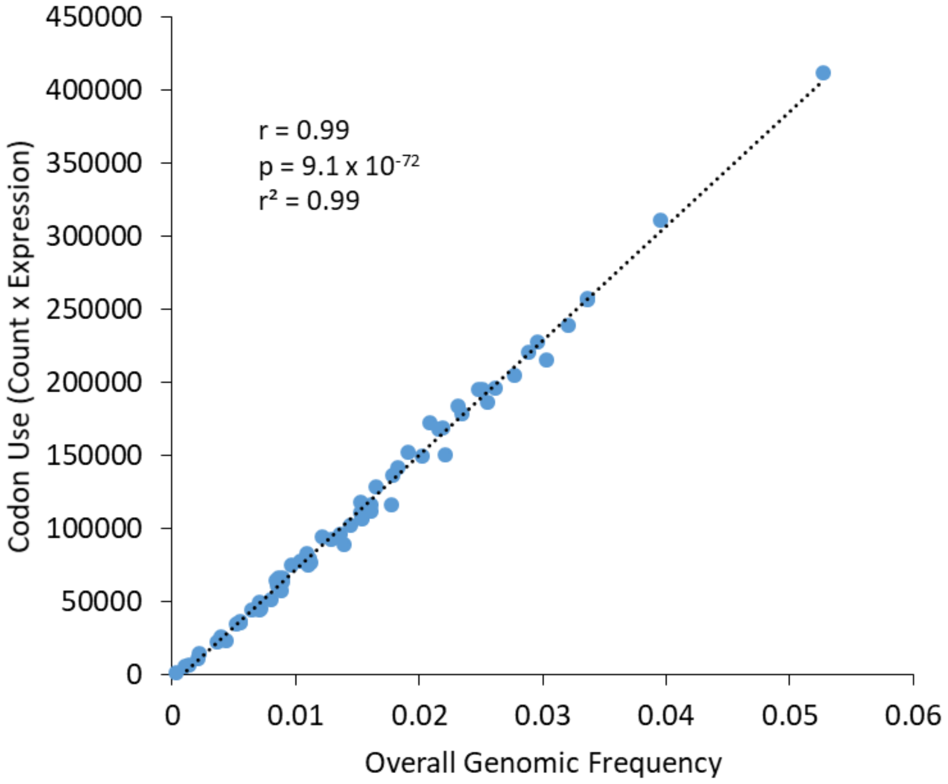


**Fig. S27.**

**Codon use vs. codon frequency in *E. coli* K12 MG1655.** Genomic frequency was used for nTE calculations in place of codon use (see methods) because they are very well correlated, or expression of genes in *E. coli* with different CUB does not appear to change codon demand in this case. Codon use was calculated as previously described (*6*) by taking the total codon count across all *E. coli* genes multiplied by their corresponding transcript abundance from a publically available dataset (GEO accession GSE59377, “rpoB_wt_lb”) (*66*), which correspond to an exponentially growing culture of *E. coli* in LB broth. Pearson correlation coefficient and linear regression $r^2$ values are calculated from $n$ = 64 codons.

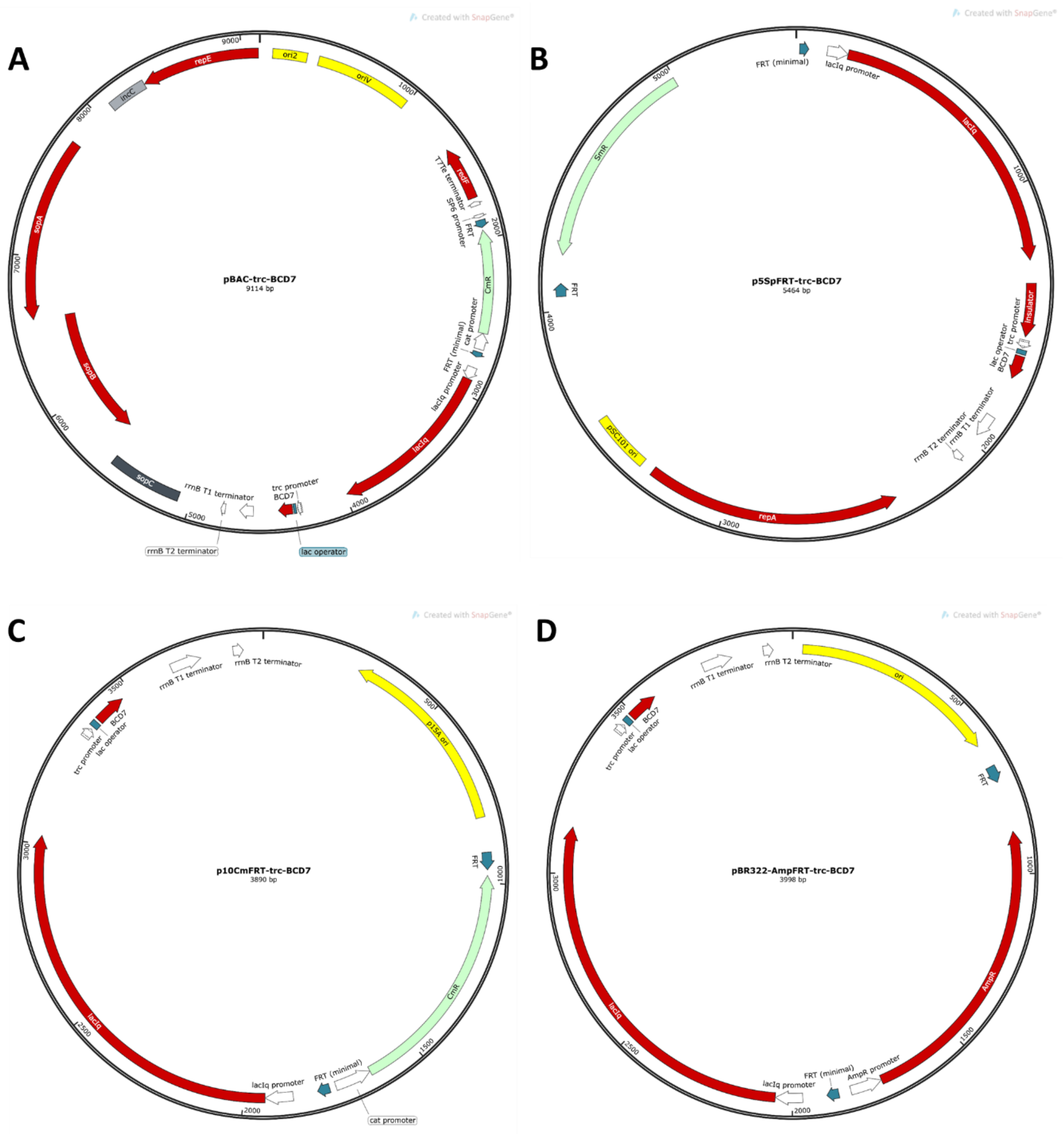


**Fig. S28.**

**Vector maps for expression plasmids used in this study.** Each vector consists of a unique origin/copy number control mechanism based on the origin of replication and in some cases accessory proteins. Copy numbers are approximated based on known values. (**A**) pBAC (F1 origin, ~1 copy); (**B**) p5 (pSC101 origin, ~5 copies); (**C**) p10 (p15A origin, ~10 copies); (**D**) p20 (pBR322 origin, ~20 copies). Full length vector sequences can be found in **Data S10**.

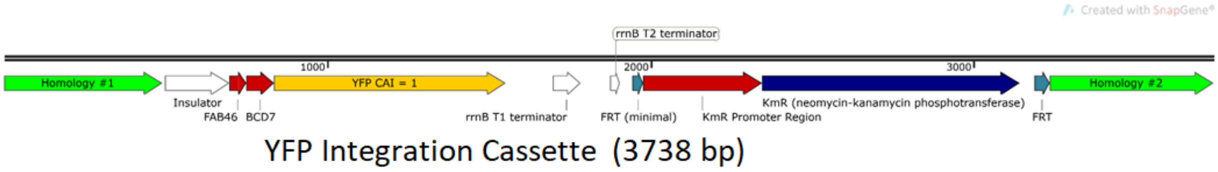


**Fig. S29.**

**Chromosomal integration cassette for YFP reporter.** The YFP cassette was integrated in the *E. coli* chromosome in between the genes *rsmG* and *atpI*. It consists of (from 5' to 3') a homology region of ~500 bp, an insulator sequence, followed by the FAB46 strong constitutive promoter, BCD7 (a strong RBS), and the YFP CAI = 0.96 (~1) sequence, and the *rrnB T1/T2* terminator. Attached to the 3' end is a kanamycin marker flanked by FRT sites for site specific marker excision with the FLP recombinase, and finally another ~500 bases of homology on the 3' end. The complete sequence can be found in **Data S10**.

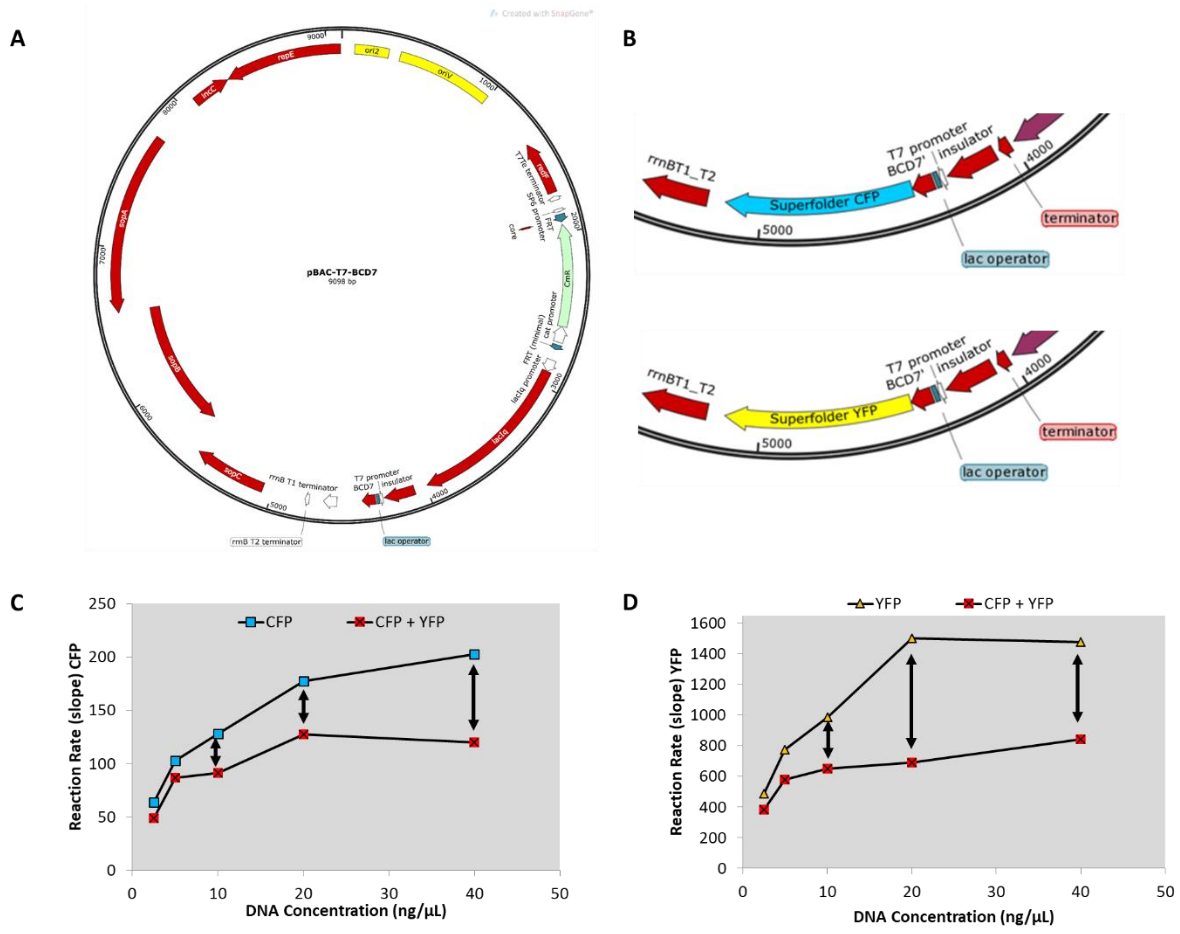


**Fig. S30.**

**In vitro transcription-translation (TxTL) assay design.** (**A**) Cloning vector used to generate the linear cassettes used in the TxTL assay. The vector includes a T7 promoter, which the NEB PURExpress® kit relies on for transcription. (**B**) Representative Linear DNA cassettes used for TxTL for CFP and YFP, including an insulator, T7 promoter and lac operator (IPTG inducible), strong RBS (BCD7), gene of interest, and *rrnB* terminator. Genes were amplified from the insulator to the terminator for use in the TxTL assay. (**C** and **D**) Expression of CFP and YFP in isolation or under competition in the TxTL assay and resulting reaction rates. DNA concentration refers to the concentration of each species of DNA added (i.e., there is 20 ng/μL total for individually expressed, or 40 ng/μL total for combined). Competition is effectively maximized around 20ng/μL of each gene.

### Superfolder CFP

```
  1  mgkgeelftg vvpilveldg dvnghkfsvr gegegdatng kltlkfictt
 51  gklpvpwptl vttltwgvqc fsrypdhmkr hdffksampe gyvqertisf
101  kddgtyktra evkfegdtlv nrielkgidf kedgnilghk leynfnshnv
                  Peptide
151  yitadkqkng ikanfkirhn vedgsvqlad hyqqntpigd gpvllpdnhy
201  lstqsvlskd pnekrdhmvl lefvtaagit hgmdelyk
```

### Superfolder YFP

```
  1  mgkgeelftg vvpilveldg dvnghkfsvr gegegdatng kltlkfictt
 51  gklpvpwptl vttltygvqc fsrypdhmkr hdffksampe gyvqertisf
101  kddgtyktra evkfegdtlv nrielkgidf kedgnilghk leynfnshnv
                  Peptide
151  yitadkqkng ikanfkirhn vedgsvqlad hyqqntpigd gpvllpdnhy
201  lsyqsvlskd pnekrdhmvl lefvtaagit hgmdelyk
```

### mCherry

```
  1  mvskgeednm aiikefmrfk vhmegsvngh efeiegegeg rpyegtqtak
 51  lkvtkggplp fawdilspqf mygskayvkh padipdylkl sfpegfkwer
                                                 Peptide
101  vmnfedggvv tvtqdsslqd gefiykvklr gtnfpsdgpv mqkktmgwea
151  ssermypedg alkgeikqrl klkdgghyda evkttykakk pvqlpgaynv
201  nikldítshn edytiveqye raegrhstgg mdelyk
```

**Fig. S31.**

**Peptides used for LC/MS analysis of protein abundance.** Note that CFP and YFP could not be differentiated as they were quantified using the same peptide, and only differ by 2 amino acids. For CFP and YFP, the peptide is 122 amino acids from the N terminus, and for mCherry it is 97 amino acids from the N terminus, thus giving confidence that the peptide quantification represents abundance of the entire protein.

| Gene Name | Full Name | Pathway/Description | CAI |
|---|---|---|---|
| gapA | glyceraldehyde-3-phosphate dehydrogenase | Glycolysis, primary metabolism | 0.88 |
| pgk | phosphoglycerate kinase | Glycolysis, primary metabolism | 0.82 |
| rpsA | 30S ribosomal subunit protein S1 | Protein synthesis | 0.84 |
| eno | enolase | Glycolysis, primary metabolism | 0.89 |
| acnA | aconitate hydratase A | TCA cycle, primary metabolism | 0.52 |
| sdhA | Succinate dehydrogenase A | TCA cycle, primary metabolism | 0.72 |
| sdhB | Succinate dehydrogenase B | TCA cycle, primary metabolism | 0.52 |
| ispA | farnesyl diphosphate synthase | Isoprenoid biosynthesis, secondary metabolism | 0.50 |

**Table S1.**

**Table describing background *E. coli* proteins monitored for abundance by LC/MS targeted proteomics.** Genes were selected to represent both higher and lower CAI levels from background *E. coli* host proteins.

| | CAI | CHI (χ) | ENC | GC |
|---|---|---|---|---|
| **AvPAL High CAI** | 0.83 | 0.81 | 24.36 | 54.9 % |
| **BcLAI High CAI** | 0.91 | 0.82 | 23.83 | 51.7 % |
| **EclacZ High CAI** | 0.86 | 0.81 | 24.17 | 57.6 % |
| **AvPAL High CHI** | 0.45 | 0.93 | 32.18 | 51.1 % |
| **BcLAI High CHI** | 0.52 | 0.95 | 34.28 | 49.3 % |
| **EclacZ High CHI** | 0.43 | 0.94 | 32.56 | 54.5 % |

**Table S2.**

**Sequence statistics for new re-coded enzymes in Figure 7.** Represented for each gene is the calculated codon adaptation index (CAI), codon health index (χ), effective number of codons (ENC), and % GC content.

**Data S1. (separate file)**
**Fig. S1 Codon Elongation Times used in RFM.**

**Data S2. (separate file)**
**Fig. 1C Raw Fluorescent Data**

**Data S3. (separate file)**
**Fig. 1E and 1F Raw Fluorescent Data**

**Data S4. (separate file)**
**Fig. 1G and 1H Raw Fluorescent Data**

**Data S5. (separate file)**
**Fig. 2E Raw Growth and Fluorescent Data**

**Data S6. (separate file)**
**Fig. 2F Raw Growth and Fluorescent Data**

**Data S7. (separate file)**
**Fig. 2G Raw Growth and Fluorescent Data**

**Data S8. (separate file)**
**Co-expression Fitness Weights for CHI**

**Data S9. (separate file)**
**Codon weights, frequencies, and RSCU values used**

**Data S10. (separate file)**
**Sequences used in the study**

**Data S11. (separate file)**
**Quality controls for qRT-PCR results.** All data were generated using mean values from 2 technical replicates. (**Sheet A**) In vitro TxTL reaction primer specificity data using reaction templates, indicating cq values for all primer pairs when tested with relevant templates for each pair. All pairs are sufficiently specific for analysis. (**Sheet B**) Primer sequences used for in vitro TxTL qRT-PCR analysis, and measured efficiency values. (**Sheet C**) In vivo primer specificity data using isolated cellular RNA, indicating cq values for all primer pairs when tested with relevant templates for each pair. All pairs are sufficiently specific for analysis. (**Sheet D**) Primer sequences used for in vivo qRT-PCR analysis, and measured efficiency values. (**Sheet E**) Cq values for amplicons normalized to the same starting concentration for relevant CFP and mCherry templates. The close cq values indicate that CFP high CAI and high $\chi$ re-coded sequences can be reasonably compared given they have very similar amplification efficiency.
